# FragmentScope - exploring the fragment space with learned surface representations

**DOI:** 10.64898/2025.12.16.694391

**Authors:** Evgenia Elizarova, Irina Morozova, Ilia Igashov, Olivia Gampp, David Robin Pavel Iosub, Arne Schneuing, Kelvin Lau, Davide Ferraris, Florence Pojer, Roland Riek, Michael Bronstein, Bruno E. Correia

## Abstract

Exploring fragment chemical space for ligand design remains a major challenge in early stage drug discovery. This task is particularly challenging due to the small size, low specificity, and weak binding affinities of low molecular weight (MW) fragments. We present FragmentScope, a computational pipeline that uses learned protein surface fingerprints to guide fragment placement and small molecule generation. By using a contrastive learning model trained on protein-ligand interactions, we built a database of surface-fragment pairs, which enables fast and accurate fragment placement in the target protein pocket. We demonstrate its effectiveness on benchmark datasets, achieving robust placement accuracy. FragmentScope also enables the design of small molecules based on the predicted fragments ensuring synthetic accessibility. We experimentally validated Fragmentscope across 5 different targets with binding assays and structural characterization. Our approach shows high success rates in fragment discovery and yielded promising leads for designed ligands. FragmentScope offers a scalable, structure-guided approach for narrowing chemical space and identifying prospective scaffolds, accelerating the early stages of small molecule design.

## Introduction

Small molecules are the largest class of therapeutics, playing a crucial role in the development of novel medicines. However, designing potent small molecules remains challenging due to different complexities associated with protein targets, the vast multidimensional space of possible chemical structures and their synthetic feasibility [1]. Many proteins are difficult to drug due to a number of features including the lack of well-defined binding pockets, structural flexibility and others [2,3]. Examples include proteins involved in protein-protein interactions (PPIs), transcription factors with high structural similarity, and small GTPases without clear ligand binding cavities [4].

From a chemical perspective, challenges include navigating the vast chemical space, meeting strict physicochemical criteria, ensuring compound stability and reactivity, and optimizing synthesis. Designing a functional small molecule often requires multi-step synthetic pathways with specific reaction conditions, making the process expensive and time-consuming. An alternative approach is high-throughput screening of large compound libraries, which involves testing thousands to millions of compounds to identify potential hits. However, the success rate remains relatively low due to limited sensitivity of screening assays and the difficulty of identifying truly novel scaffolds [5,6]. Furthermore, experimental validation such as testing and confirming the binding affinity, potency, and pharmacokinetic properties of candidate molecules is a crucial, yet rather laborious and expensive step, particularly when working with large compound libraries.

Fragment-based drug design (FBDD) is one of the small molecule design approaches which have been commonly used to overcome some challenges and achieve balanced synthetic feasibility with target specificity. Unlike traditional high-throughput screening, which relies on large and complex molecules, FBDD operates on libraries of smaller, less complex molecular building blocks, known as fragments (typically with molecular weights below 300 Da). These fragments are screened against target proteins to identify weak but actionable interactions, which can then be optimized by growing, linking, or merging fragments into larger, higher affinity compounds.

FBDD has several advantages over traditional screening methods. Fragments, due to their small size and potential for combination, enable more efficient exploration of the chemical space. By occupying different areas within a binding pocket, they facilitate high ligand efficiency, meaning that each atom contributes more to the overall binding affinity. Second, fragment libraries are relatively small (hundreds to thousands of compounds), making them easier to screen and optimize, compared to potentially synthesizable compound libraries, which can reach in silico billions of molecules. However, despite high ligand efficiency, the overall low binding affinity (high micromolar to millimolar range) of fragments poses a challenge, requiring various structural biology methods, such as X-ray crystallography and NMR spectroscopy, to determine binding interactions [7,8]. Additionally, optimizing weak fragment hits into high-affinity drug candidates is a non-trivial multi-step process that often requires precise chemical modifications [9]. Even though various experimental techniques are commonly used for fragment screening, the success rate of identifying a hit remains relatively low (around 5-10%) and highly depends on the library, target and method [10,11]. Computational approaches can help to address this limitation by providing a rapid and cost-effective screening and prioritizing fragment hits before experimental validation. Such a combined approach with a computational pre-screen followed by optimisation and experimental evaluation has been applied to several targets including Colony-stimulating factor 1, VEGFR-2, Glyoxalase 1 and others [12,13].

While FBDD and traditional computational methods such as molecular docking and structure-based drug design have already contributed significantly to drug discovery [14–16], they often struggle with generating entirely novel chemical entities with high binding affinity. Recently, AI-driven approaches have emerged as promising complementary strategies. By leveraging machine learning models trained on extensive datasets, these approaches aim to predict protein-ligand interactions and facilitate small molecule design [17–23].

This paradigm is slowly but steadily reshaping early-stage drug discovery, with shifting focus on data-driven models that can extract meaningful patterns directly from structures. Rather than relying on particular physical descriptors or mechanistic models which may be limited in capturing the binding signals of small fragments. These approaches leverage large-scale data to learn predictive relationships from chemical structures. Data oriented strategy is particularly valuable when dealing with fragments, where the physical signals and patterns emerging from the chemical structures may be weak or ambiguous.

Here, we introduce FragmentScope, a new pipeline that employs fragment placement in the target protein pocket based on learned protein surface embeddings to facilitate small molecule design (Figure 1A). Trained with a contrastive learning objective on thousands of protein pocket-ligand pairs, our model enables accurate fragment positioning within binding sites based on geometric and physico-chemical similarities. Our method uses learnt fingerprints to guide fragment placement by matching both the shape and chemical features of the protein surface, helping overcome key limitations of traditional fragment-based drug discovery. Additionally, by integrating fragment placement with linker generation, our method ensures both molecular novelty and capturing of crucial protein-ligand interactions. This AI-powered approach provides a robust and efficient framework for designing novel small molecules with therapeutic potential.

**Figure 1.**
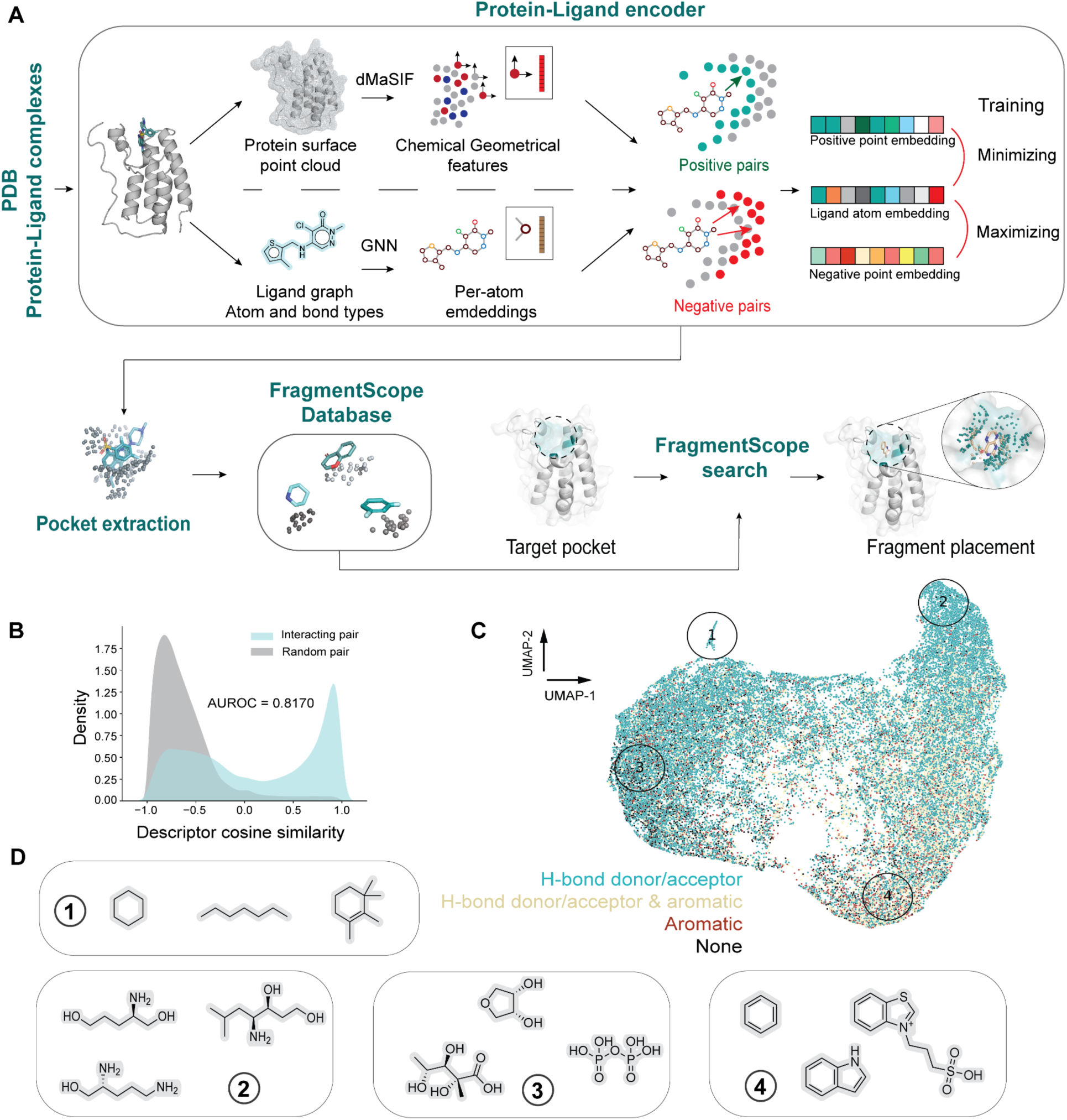
Computational framework for surface-based fragment placement. **A)** Overview of the surface based fragment placement using the protein-ligand encoder trained with a contrastive loss. Proteins are represented as a surface point cloud generated with the dMasif model expressing chemical and geometrical features. Small molecules are represented as graphs and processed with a GNN model using per-atom embeddings (see Methods ‘Protein-ligand encoder’). We trained the model with pockets from protein-ligand complexes and assembled a database of fragments, this pipeline was dubbedFragmentScope. The database is used to search and place fragments. **B)** Distributions of cosine similarities of the surface patch embeddings and fragment atom embeddings of interacting pairs (blue) and random pairs (grey), respectively. **C)** UMAP projection of the database of surface embeddings together with examples of corresponding fragments from the circled areas. **D)** Examples of some fragments in the FragmentScope database.

## Results

### In silico fragment selection with protein surface fingerprints

Navigating fragment chemical space is a challenging task due to the low MW and the chemical simplicity of fragments. Their small size results in high motion range, weak and sparse interaction patterns, and consequently low binding affinities. These factors make it difficult to distinguish true binders from nonspecific interactions, complicating both computational predictions and experimental validation. Traditional computational approaches, such as molecular docking, often struggle to efficiently explore and reliably position fragments within protein binding sites [10]. In this study, we leverage learned protein surface fingerprints to enhance fragment placement, ensuring a selection of relevant fragments within target protein pockets.

To generate these protein surface fingerprints, we built upon a previously established surface-encoding framework [24,25] and further developed it to predict protein-small molecule interactions. Our approach employs two neural networks that independently process and encode protein surfaces and small molecule fragments. The model was trained using a contrastive loss to discriminate between interacting (positive) and non-interacting (negative) atom-surface patch pairs (Figure 1A, Methods “Protein-ligand encoder”).

We processed protein-ligand complexes from the Protein Data Bank (PDB) [26], extracted corresponding pockets and created a database of pairs of small molecule fragments and their corresponding surface patches. We developed an engine called FragmentScope, that searches across that pre-built database. By doing so, one can utilize pocket surface patches as a query and efficiently search for fragments that are likely to bind to the query surface patch and provide a 3D pose for the interaction mode. The database consists of the fragments from the PDB paired with a surface patch extracted from the corresponding protein pockets, as shown in Figure 1A and Figure S1. To find and place the most relevant molecular fragment in the pocket of interest, FragmentScope represents the pocket as a three-dimensional point cloud with learned descriptors assigned to each point (Figure 1A, Figure S1) and compares it with the available pockets in the database. The most similar candidates are further aligned with the query point cloud using the 3D alignment algorithm RANSAC [27]. The resulting alignment transformations are also applied to the corresponding fragments, which places them in the query pocket. We select the final set of fragment candidates based on the computed similarity score between the aligned point clouds (see Supplementary ‘FragmenScope’).

The similarity of embedding vectors of the trained model is directly indicative of molecular interactions and can distinguish between interacting and non-interacting ones (Figure 1B). The resulting surface embeddings also encode chemical features of the bound fragments, as illustrated in Figure 1C. The labelled fragment examples show how certain chemical moieties are enriched in different parts of the fingerprint space. For instance, fragments bound to surfaces from group (2) tend to have a large number of hydrogen bond acceptors or donors whereas group (4) is mostly characterised by aromatic systems.

Two fragment databases split by molecular-weight ranges were created from known protein-ligand interactions in the PDB. The first database consists of larger, chemically richer fragments suitable as initial scaffolds (MW between 150-500 Da), while the second database contains smaller fragments designed for combinatorial linking into synthetically viable molecules (MW between 60-300 Da). Both databases leverage structural knowledge from existing complexes, providing a robust foundation for fragment placement. Further details are provided in Supplementary, section Databases.

### Prediction of fragment placement and chemical space exploration

To evaluate the accuracy of the fragment placement algorithm, we curated a set of five target proteins: K-Ras GTPase (KRAS, PDB ID: <u>8AZR</u>), Bromodomain-containing protein 4 (BRD4, PDB ID: <u>7WL4</u>), TEAD2 transcription factor (TEAD2, PDB ID: <u>8CUH</u>), Mitogen-activated protein kinase 1 (ERK2, PDB ID: <u>2OJI</u>), and SARS-CoV-2 Main Protease (SARS Mpro, PDB ID: <u>7Z0P</u>). These structures were selected due to their relevance in drug discovery and their availability in complex with small molecules.

The benchmarking process consisted of two main steps. First, we created a ‘ground truth’ dataset by extracting all publicly available ligand-bound crystal structures of homologs with at least 80% sequence similarity to the target sequences. Next, we applied FragmentScope to each target structure and shortlisted the top 200 fragment candidates along with their 3D positions (see Methods ‘FragmentScope’). Proposed fragments and their placements were then compared to the reference fragments to assess the recovery rate. For all 5 targets FragmentScope was able to retrieve ‘ground truth’ fragments (Figure 2A,B). Proteins with relatively low occurrence in the Database 1 (Figure 2A, Supplementary ‘Databases’) such as TEAD2 showed lower recovery rate. However, proteins that are structurally abundant in the Database 1 like BrD4 target - demonstrated high recovery rate of the ground truth fragments - 82.5%, as well as SARS Mpro, ERK2 - 38%, 32.5% respectively (Figure 2B). KRAS, despite the relatively high occurrence, showed low recovery rate, which could be due to the known flexibility of the protein pocket [27].

**Figure 2.**
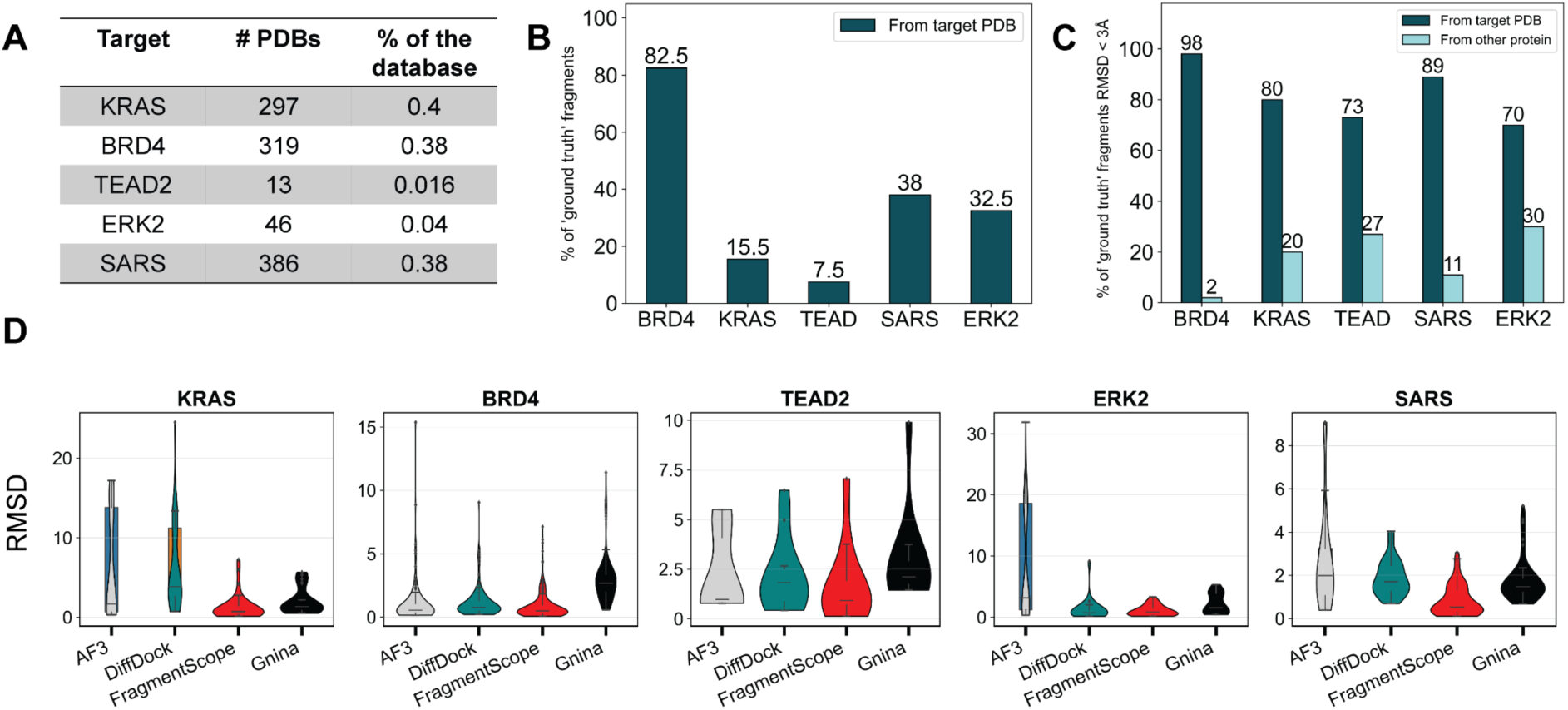
Benchmark evaluation of the surface-based fragment placement. **A)** Representation of the target structures in the Database 1. **B)** Recovery of ‘ground truth’ fragments for 5 targets used for the benchmark. **C)** Recovery of the ‘ground truth’ fragments with distinct structural origins: homologous to the target PDB structures and other complexes. **D)** RMSD distributions of the benchmark fragments placed with FragmentScope, AF3, DiffDock and Gnina.

Notably, FragmentScope also accurately retrieved chemically and structurally similar fragments from structurally distinct protein pockets even in cases of limited data availability (Figure 2C). For TEAD2 and ERK2 proteins with low structural availability in the Database 1 we observed promising recovery rate of ‘ground truth’ fragments from distinct protein pockets of 27 and 30% respectively. In contrast, most of the ‘ground truth’ fragments for BrD4 protein were from homologous structures thus only 2% of fragments were from other protein complexes. Additionally, FragmentScope is able to perform the recovery of fragments for the protein with a flexible binding pocket - KRAS with the value of 20%. This demonstrates a broader capability of FragmentScope: the learned surface embeddings capture meaningful geometric and chemical similarities that generalize beyond immediate structural homologs. Thus, the method is highly accurate when similar pocket structures exist in the database but still offers substantial value and reliable placement poses when direct matches are not unavailable. This balanced capability highlights both the specificity and generalizability of FragmentScope, making it applicable in various fragment-based drug discovery scenarios.

Fragment placement tasks in protein pockets are often addressed as docking-based virtual screening approaches applied to large fragment libraries. However, such methods in many cases have low precision, yielding many false positives and unreliable pose predictions - especially for small, low-affinity fragments.

In contrast, FragmentScope leverages learned protein surface fingerprints to guide fragment placement by identifying regions of structural similarity to known ligand-bound pockets. While our approach is limited in some cases to finding a good match, its strength lies in efficiently prioritizing structurally relevant fragments without the need to exhaustively dock large libraries. This allows us to explore the chemical space with higher precision.

FragmentScope achieves higher placement accuracy for KRAS, BRD4, SARS and ERK2 targets compared to AF3 and is also superior to physics-based docking approaches like GNINA for all test cases: KRAS, BRD4, TEAD2, ERK2, SARS (Fig. 2D). Rather than replacing docking approaches, FragmentScope offers a complementary and more precise alternative by narrowing down the list of candidate fragments through the protein surface-based search before downstream optimization or linking.

### Experimental testing of computationally identified fragments

The first application of FragmentScope was to identify novel fragments capable of binding to target pockets. This computational screening aimed to identify a set of fragments with high binding propensity and specificity, thereby increasing hit rates even in small fragment libraries. The key challenge was to accurately detect the binding interaction and to distinguish specific interactions from nonspecific, given that typical affinities for fragments can range from mM to tenths of μM in terms of K_D_ (dissociation constant), and many of them are only detected through X-ray crystallography or nuclear magnetic resonance spectroscopy (NMR). To test the computational predictions, all screened compounds were experimentally tested using Grating-Coupled Interferometry (GCI), and several also with NMR which enables residue-level detection of compound induced chemical shift perturbations in the protein structure. Thus, allowing identification of interaction sites. Specificity of the fragments was assessed by evaluating their binding to off-target proteins and at two different concentrations. The compound selection process followed the pipeline outlined below.

For four protein targets: KRAS, SARS Mpro, PGK1 and PIN1 FragmentScope was used to search a curated fragment database (Database 1), yielding 200 candidate fragments per target. All “ground-truth” fragments (those derived from the same or homologous protein-ligand crystal structures) were excluded. To enrich novel drug-like candidates, additional filters were applied. Only fragments within 7 Å of the pocket center were retained. Fragments containing long carbon chains (>4 atoms) or having rare elements (As, Cu, Se) as well as phosphates were removed. The remaining novel fragments were then matched against the commercial Enamine REAL compound Database (∼18 billion compounds) using the InfiniSee SpaceLight tool [28] with tanimoto similarity on functional-class fingerprints (fCSFPs). The selected fragments were searched in the Enamine Building Blocks database using maximum substructure similarity, resulting in 40 compounds that were purchased and tested experimentally.

All fragments were screened using GCI binding assay at two concentrations (2 mM and 500 μM). Each fragment was also tested against two off-targets (KRAS and PIN1 or SARS Mpro). Detecting hit fragments is a challenging task due to their low affinity, specificity, and molecular weight. Hence, conditions were optimized to detect weak, fast-on/fast-off protein-fragment interactions (see Methods ‘Grating-Coupled Interferometry’).

For the PGK1 target, the hit rate was the highest, with 9 of 10 fragments binding at 2 mM. Four of these retained specificity at 500 μM (Figure 3A). Few PGK1 fragments were selected for NMR validation: 160442, 21473, 23988 as a negative example (Supplementary Table 2). Noticeable shifts were observed for 23988, a structurally novel fragment for PGK1 in solution (Figure S2).

**Figure 3.**
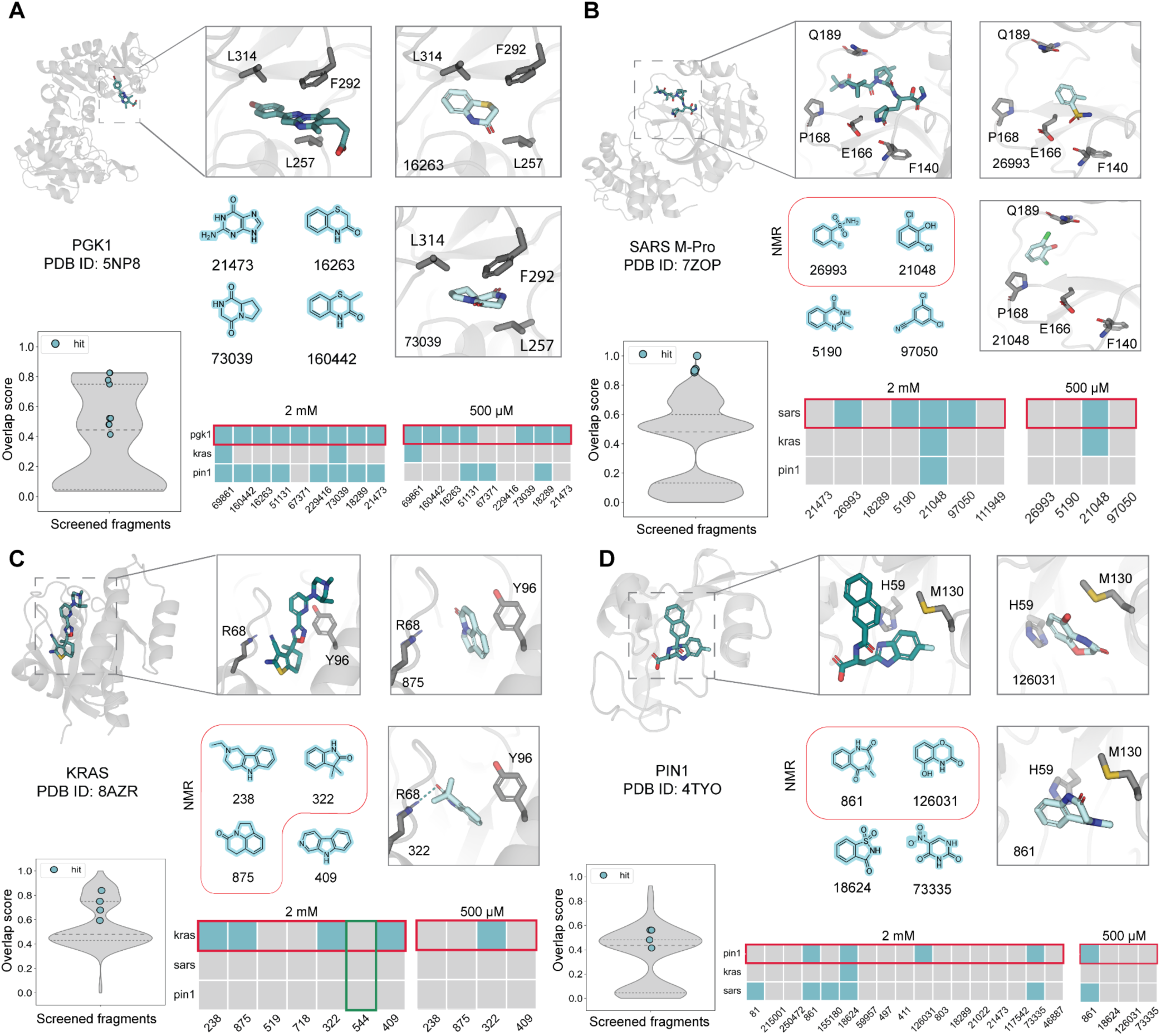
Overview of the fragment screening results for the four protein targets. Structural analysis and experimental testing of fragments targeting: **A)** PGK1 **B)** SARS Mpro **C)** KRAS **D)** PIN1. Each panel contains: 1) Crystal structure of the reference protein-ligand complex defining the binding pocket. 2) Binding of the reference compound from the crystal structure as well as predicted 3D poses of two experimentally confirmed ‘hit’ fragments placed with FragmentScope shown in sticks (cyan color). Key residues of the target pocket are shown in dark grey sticks. 3) Chemical structures of 4 ‘hits’; NMR-confirmed hits are highlighted with a red framing. 4) Histogram violin plot of the maximum substructure overlap of the screened fragments to the known molecules from the PDB, dots representing the highest overlap value for ‘hits’ fragments. 5) Binding heatmap across the primary target and two off-targets: red marks the intended target; blue indicates binding; grey denotes no binding.

Results of the screening for SARS Mpro showed that four out of seven novel fragments were hits at 2 mM. Three of them showed specificity over KRAS and PIN1. One fragment retained binding to SARS Mpro and KRAS at 500 μM (Figure 3B). Fragments 26993 and 21048 were validated orthogonally by NMR at 500 μM, as evidenced by chemical shift perturbations in the spectra (Supplementary Table 3, Figure S3). Although lack of specificity was observed at the higher 2 mM concentration, this behavior is expected due to the inherent weak and transient binding modes of fragments. Moreover, for several fragments there was no binding detected, likely due to their aggregation at high concentrations. Fragments identified as hits at both concentrations represent valuable starting points for further optimization into lead compounds. Notably, those that showed specific binding at the lower concentration (500 µM) may indicate improved specificity, offering additional advantages for selective binding to the target.

Seven fragments were experimentally tested for binding to KRAS, including one fragment (544) extracted from an existing KRAS structure with a compound bound to a Switch I/II pocket of KRAS [29]. Four of the six novel fragments showed binding at 2 mM, one fragment (322) retained binding at 500 µM, displaying selectivity for KRAS over tested off-target proteins (Figure 3C).

Interestingly, fragment 544 did not show detectable binding under the same conditions. This lack of binding is likely due to its structural simplicity that is insufficient for a detectable binding interaction, despite that it was derived from a known KRAS binding ligand. This highlights the value of combining computational predictions with experimental validation to accelerate the discovery of new chemical matter for drug design

PIN1 appeared to be the most challenging target, among 17 tested fragments, we identified four hits at 2 mM. Two retained specific binding at 500 μM. Overall, the fragment hit rates at 2 mM ranged between 23% and 100% (57% for KRAS, 100% for PGK1, 57% for SARS-CoV-2 Mpro, and 23% for PIN1) and were generally higher than those observed in experimental library screens with non-targeted fragment sets [10,30,31].

Despite the challenges to detect binding of fragments at low concentrations, orthogonal experimental validation of several fragments for KRAS and PIN1 confirmed our observations. Fragments 875 and 322 showed large shifts on both residues M67 and G75 (Figure 4A), confirming their binding to KRAS (Supplementary Table 4). Additionally, among three fragments validated to KRAS with NMR, fragment 238 showed the largest chemical shift perturbations (Figure 4A), with multiple residues in the predicted pocket being perturbed, including ARG68 and GLU62 where the largest shift was observed (Figure 4B). These results indicate the potential of this fragment to become a promising scaffold for subsequent ligand design. Experimental validation for the PIN1 fragments also confirmed binding of the 126031 fragment indicating shifts of: SER154, GLN131 within the pocket of interest (Figure 4C, Supplementary Table 5). As well as the 861 fragment was confirmed to bind to the target, shifts were observed for residues ARG69 and SER71 (Figure S4). In addition, fragment 155180 was successfully crystallized in complex with PIN1 which confirmed its binding within the predicted pocket (Figure S5). Even though the placement was not the same as we predicted (RMSD 3.98 Å), the location of the fragment within the pocket was accurate (Figure 4D). Taking into account low molecular weight and possibility of the fragment to bind to many other areas within the target pocket or other potential pockets, the crystallographic structure provided meaningful data, highlighting the potential of our approach for fragment placement.

**Figure 4.**
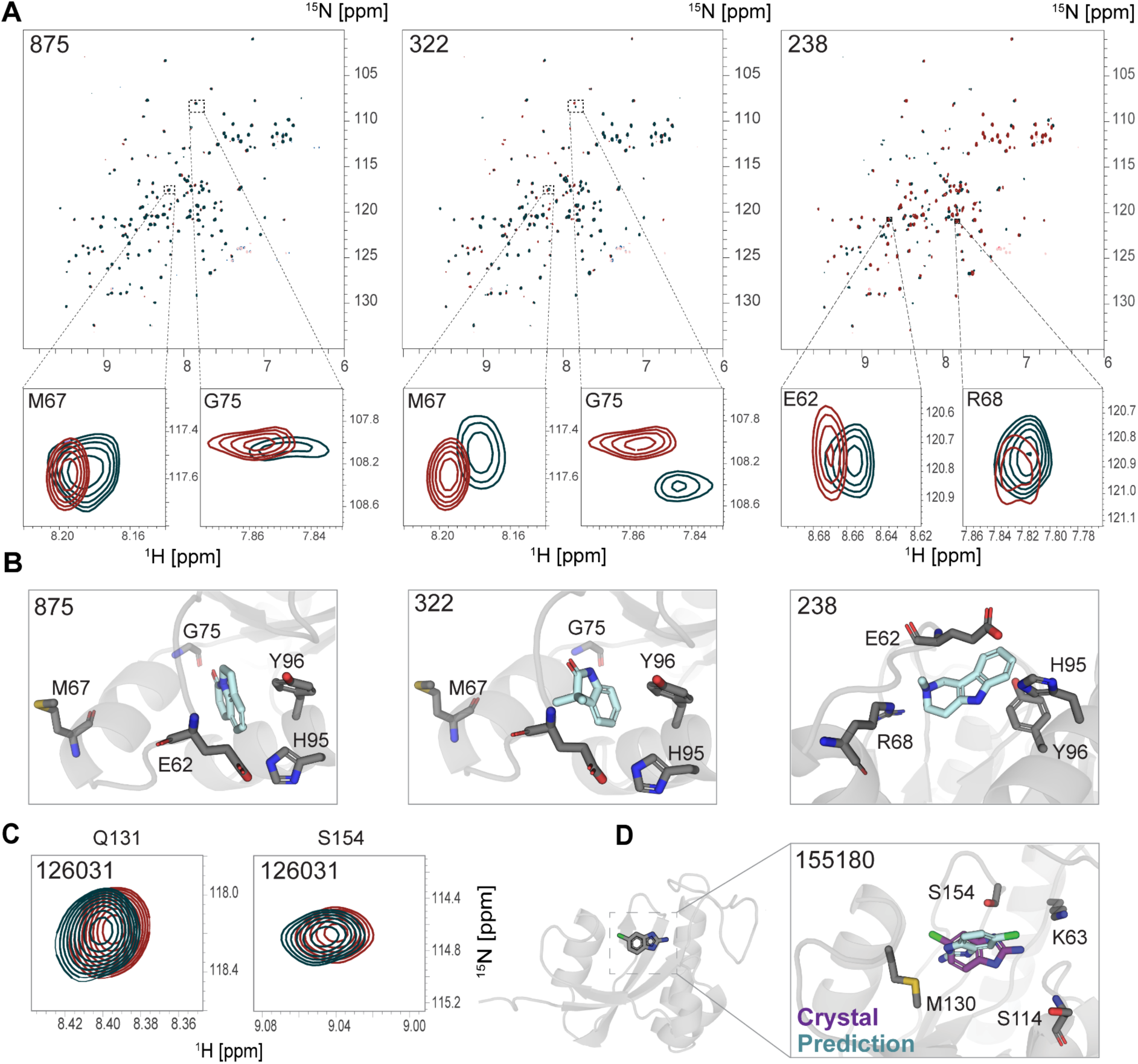
Experimental validation [15N,1H]-HSQC NMR spectra and structural analysis of FragmentScope designed fragments targeting KRAS and PIN1. **A)** [^15^N,^1^H]-HSQC overlay of apo-KRAS G12V 100 µM (teal) bound to novel fragments 875, 322 and 238 (red), binding of all fragments were confirmed. Both residues GLU62 and ARG68 peaks shift upon addition of 2 mM of fragment 238 (red). For fragments 875 and 322 both residues M67 and G75 have large shifts confirming binding (red). **B)** Predicted binding modes for designed fragments 875, 322 and 238 in the Switch I/II pocket of KRAS. **C)** [15N,1H]-HSQC overlay of 200 µM apo-PIN1 (teal) bound to 0.5 mM of the fragment 126031 (red), binding of the 126031 fragment was confirmed. Both residues GLN131 and SER154 peaks shift upon addition of 0.5 mM ligand (red). **D)** Figure shows the placement of 155180 in crystal structure (purple) and the predicted pose with the FragmentScope (cyan) bound to PIN1.

Collectively, these results show that FragmentScope efficiently enriches novel, target-specific fragments. Substructure analysis further underscored their novelty, demonstrating that most identified hits do not match known ligands found in homologous structures in the PDB. Thus, FragmentScope effectively explores previously inaccessible chemical space, identifying structurally novel scaffolds with clear potential for further therapeutic development.

### Fragment-based small molecule design

Subsequently to the fragment placing task, we aimed to design higher molecular weight compounds starting from FragmentScope fragments. By combining protein surface fingerprints with fragment placement, following linker generation, our approach provides a rational workflow for small molecule design. The ability to identify and position novel fragments based on structural similarities across protein surfaces represents a new tool in the computational drug discovery toolbox.

To show the potential of our pipeline in guiding fragment-based drug discovery, we applied our computational strategy to several therapeutically relevant targets: KRAS, BRD4, PGK1, PIN1, and SARS Mpro. Top-scoring fragment placements, identified through surface fingerprint similarity, were further processed using our fragment-based ligand generation pipeline (Figure S6). This pipeline entailed several steps: first, fragments were computationally combined based on the compatibility of their binding modes and structural orientation; second, the DiffLinker tool was employed to generate linkers that bridge the fragments preserving the original binding poses [32]; finally, the resulting molecules were filtered based on synthetic feasibility (synthetic accessibility score < 3.5 [33,34]). Small molecules that passed the filtering stage were searched in the Enamine REAL database for exact-match compounds. All compounds found in the Enamine database were purchased and further experimentally tested to test binding and specificity. They were further validated using docking to check the recovery of key interactions (see Methods Section ‘Small molecules design pipeline’) and analyse the alignment with the experimental data.

Our integrated computational-experimental pipeline enabled the transition from initial fragment hits to validated small molecule scaffolds, facilitating the discovery of novel, selective, and synthetically accessible drug lead candidates. The goal of this effort was to demonstrate that surface fingerprint-guided fragment placement can significantly narrow the chemical space while enriching compounds with binding activity.

To evaluate the generalizability of our developed pipeline, we also attempted to design full small molecules, without the intermediate fragment validation, against three structurally and functionally distinct targets: SARS Mpro, PGK1, and PIN1. Designed compounds showed measurable binding to their respective targets, with Z559 (SARS Mpro) and Z123 (PGK1) showing the most consistent binding responses and partial selectivity at sub-millimolar concentrations (Supplementary Section ‘Experimental validation of designed compounds’, Figure S7). Although some off-target binding was observed, these results demonstrate that the pipeline can generate chemically novel scaffolds with experimentally confirmed target engagement, highlighting its potential for fragment-based design beyond known chemical matter. More details in Supplementary Figures S8-S10.

Exploring further the capability of our pipeline we applied it to the oncogenic target KRAS, a GTPase long considered “undruggable” due to its shallow and highly dynamic binding surfaces. Some success has been recently achieved with the first two approved FDA drugs targeting G12C KRAS mutant - sotorasib and adagrasib [35,36]. KRAS mutations are known to cause constitutive activation of the protein, leading to uncontrolled cell proliferation [37]. Two distinct pockets have been targeted in efforts to inhibit KRAS: the Switch I/II pocket and the SIIP (Switch II Pocket) [38,39]. In our study, we focused on designing small molecules for the SIIP based on structural information from the known binder Precursor 1 (PDB ID: 8AZR) (Figure 5A) [40].

**Figure 5.**
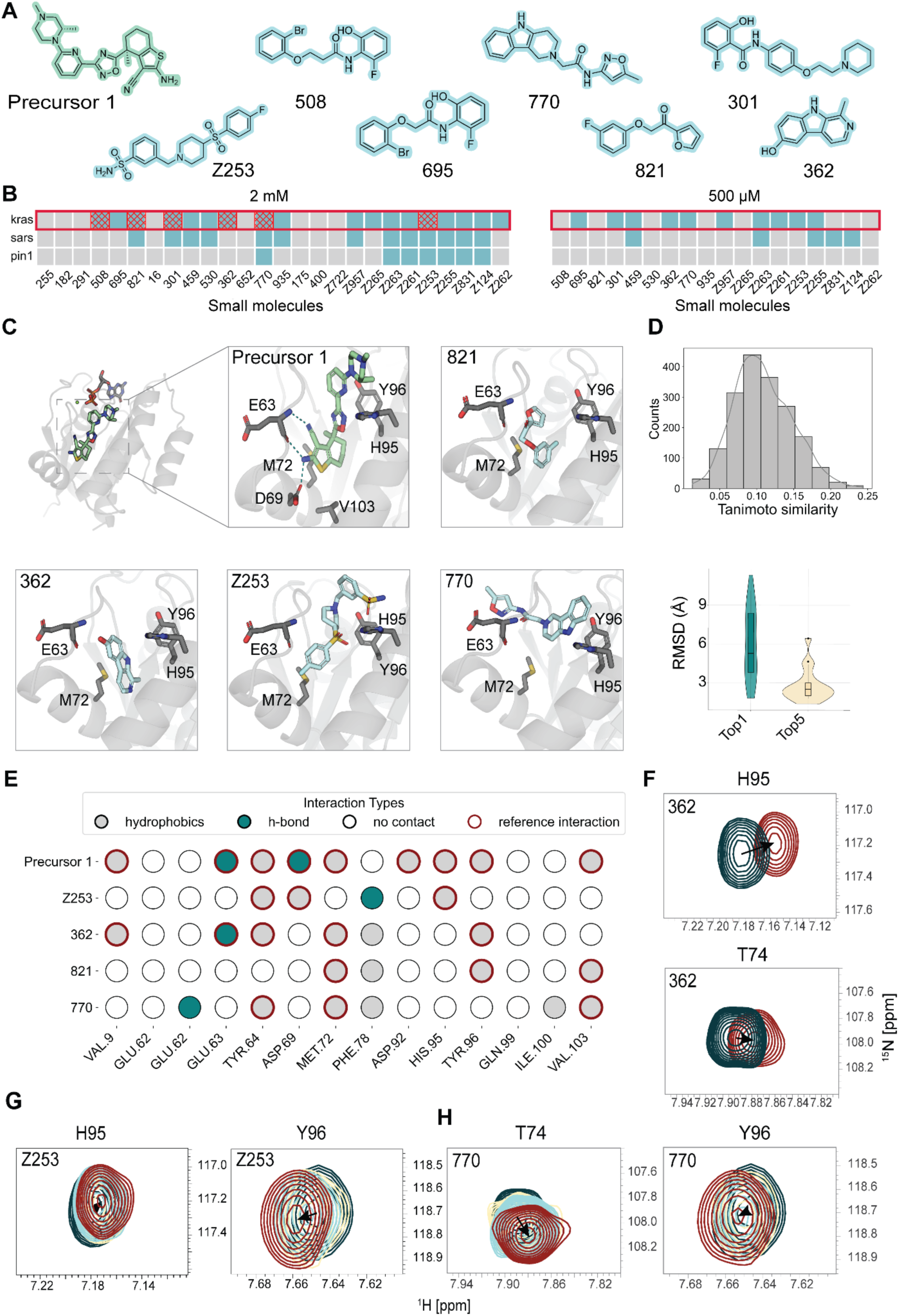
Experimental validation and structural analysis of FragmentScope designed small molecules targeting KRAS. **A)** Selected ligands for KRAS designed by FragmentScope. **B)** Heatmaps indicate binding specificity (blue - binding, grey - no binding) across KRAS, SARS, and PIN1 as determined by kinetic experiments at 2 mM (left) and 500 µM (right). NMR confirmed binders are labeled in red dashed marks. **C)** Predicted binding modes for designed ligands 821, Z253, 770, 362 and the known ligand Precursor 1 (PDB ID: 8AZR) in the Switch I/II pocket of KRAS. **D)** Tanimoto similarity distribution highlights chemical novelty of the designed compounds (left histogram), while RMSD violin plots compare docking pose accuracy (Top1 vs Best-of-Top-5) predicted by GNINA. **E)** Interaction map shows contacts for selected compounds compared to Precursor 1. **F)** The binding of ligand 362 was confirmed. Both residues TYR74 and HIS95 have peaks shift upon addition of 2 mM ligand (red). **G)** Chemical shifts perturbations for both residues - HIS95 and TYR96 confirm Z253 compound binding. **H)** The chemical shifts of the 2 residues TYR74 and TYR96 are shown with varying amounts of ligand 770 - derivative of 238 fragment. The colour coding corresponds to the various concentrations of the ligand, and was done with 80 µM KRAS.

A diverse set of small molecules was generated and tested against KRAS. These compounds fell into three main categories (Supplementary Table 1). The first one was ‘hit’ fragments derivatives: compounds selected from an Enamine-provided library of derivatives based on known KRAS fragment hits (see ‘Experimental testing of computationally identified fragments’). These were docked using GNINA, and ranked based on docking score and how well they preserved the core fragment pose predicted by FragmentScope. The second group of compounds included molecules designed with the developed pipeline based on Database 1, which consists of low MW fragments (see Methods ‘Databases’). The third group consists of the molecules also created with our pipeline but based on fragments from Database 2, larger MW fragments. In addition fragments were filtered based on their origin. We kept only those that were extracted from the ligands that bind to structurally distinct to target proteins. Designed compounds were purchased from the Enamine REAL database and experimentally tested. All details, including SMILES and source fragments, are provided in Supplementary Table 1.

Twenty-six compounds designed for KRAS were tested for binding and specificity to the target (Figure 5A, Supplementary Table 1). Experimental binding validation was performed using GCI, with an initial screening concentration of 2 mM, resulting in 18 out of 26 compounds showing measurable binding to KRAS (Figure 5B, heatmap 2mM). Some examples of the curves obtained in the experiment for several compounds such as Z255, Z261, Z957 and 770 presented in Figure S11. All the hits were identified according to our hit criteria (for details see supp. section ‘Grating coupled interferometry’). As a positive control we used BI-2865, which is an enhanced potent derivative of the precursor 1. The experimental K_D_ determined for BI-2865 positive control in our experiment was 6.9 nM, similar to that reported in literature (K_D_ = 4.5 nM) [41]. To evaluate specificity, hits were tested at a lower concentration (500 μM) against two off-targets: SARS and PIN1. Seven compounds maintained selective binding to KRAS under these conditions (Figure 5B, heatmap 500 μM), highlighting the potential for further development of compounds to become potent and selective inhibitors for KRAS-driven cancers.

It was not possible to determine precise K_D_ for the compounds given that many of them seem to range from high micromolar to low millimolar affinities (Supplementary Figure S11). Several reasons can lead to this outcome: the first one is the compound’s potency, despite the detected affinity, the compound’s kinetics can be too fast to be able to determine it accurately; mixtures of diastereomers, enantiomers which may lead to an underestimation of their potency; and high percent of salt presence such as TFA, HCI, HBr in amino compounds. To validate binding interactions observed in GCI, [¹H,¹⁵N]-HSQC NMR spectroscopy was performed on a subset of 18 compounds (Figures S12,13; Supplementary Tables 6,7). NMR spectroscopy measurements confirmed six (301, 508, 821, Z253, 362, 770) out of 18 tested compounds considered as hits based on the GCI screen (Figures 5B).

Since we can get residue level mapping of the binding interactions we can compare the predicted binding mode with the shifts obtained experimentally. Four compounds are shown in their predicted placement in Figure 5C, occupying Switch I/II pocket interacting with several key residues in the pocket, shown in sticks. Importantly, tanimoto similarity analysis (Figure. 5D, histogram) showed that the selected compounds sample a diverse chemical space from the previously known compounds targeting SIIP, underscoring their novelty and optimization potential. Interaction analysis revealed partial overlap with Precursor 1 contacts, along with new favorable interactions, such as a hydrogen bond between Z255 and TYR96, and between compound 770 and GLU62 (Figure 5E). Provided data highlights that the surface fingerprints learnt from protein-surface ligand interactions capture the key interactions of the binding mode.

For three compounds (Z253, 770 and 362) three pocket residues showed altered chemical shifts: GLY75, HIS95, TYR96. For compound 362 the largest chemical shift was observed in HIS95 supporting the binding to the target pocket (Figure 5F). Large shifts were observed for GLY75 for compounds Z253 (Figure 5G, Figure S13) and 770 (Figure 5H, Figure S13). Although the NMR titration curve for GLY75 is not fully saturated, it shows the dose-dependence from a 770 compound concentration (Figure S15). Shifts corresponding to HIS95 and TYR96 indicate the binding of the 770 compound to the target pocket (Figure 5H, S13). The titration data by GCI and NMR shows the dose dependence binding of the compound, although it was not possible to define the precise K_D_ (Figure S14).

Retrospectively analysing with docking the predicted poses for the compounds we saw top-ranked poses did not always perfectly match the predicted fragment placements; however, accuracy improved when comparing the correct pose among the top-5 predicted docking solutions rather than only the top-1 pose, indicating the robustness of the fragment-guided placement signal (Figure 5F, Supplementary Table 8).

In conclusion, to address the challenge of identifying low-affinity fragment hits, as well as ligands, two complementary approaches were employed: GCI and [¹H,¹⁵N]-HSQC NMR spectroscopy. Both methods confirmed a substantial subset of hits, with KRAS showing the highest success rate - 100 % for fragments and 37% for ligands. Additionally, specific binding residues were identified by [¹H,¹⁵N]-HSQC NMR, as designed by the FragmentScope method.

To further assess the performance of our fragment-guided molecule design pipeline, we applied it to BRD4, a well-characterized epigenetic regulator and therapeutically relevant target in oncology and inflammation [42]. A total of 28 designed compounds were computationally generated and experimentally screened for binding to the acetyl-lysine binding pocket of BRD4 (PDB ID: 7WL4) (Figure 6A, Table S1).

**Figure 6.**
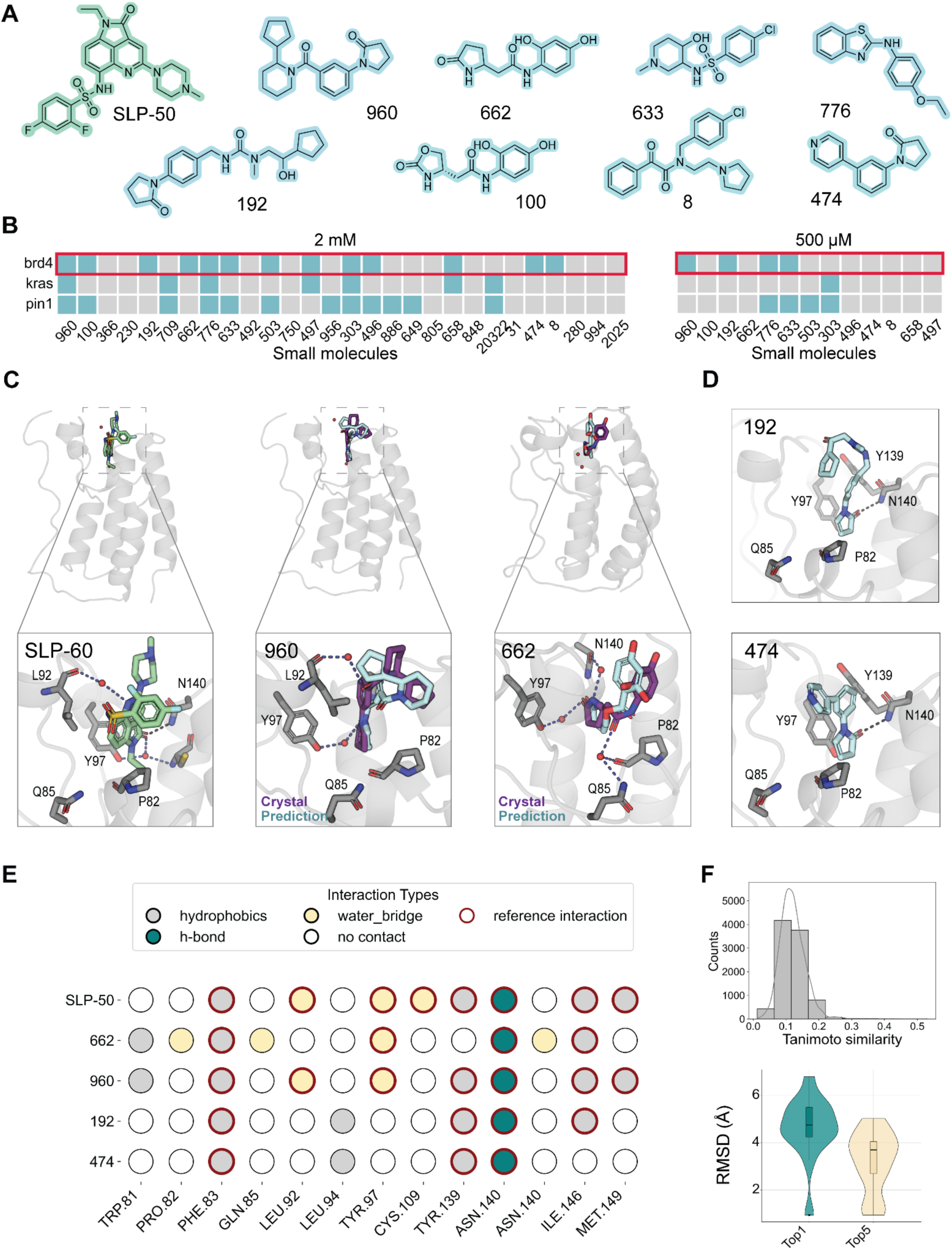
Experimental validation and structural analysis of FragmentScope designed small molecules targeting BRD4. **A)** Set of some ligands tested for BrD4 designed by FragmentScope. **B)** Heatmaps indicate binding specificity (blue - binding, grey - no binding) across KRAS, SARS, and PIN1 as determined by kinetic experiments at 2 mM (left) and 500 µM (right). **C)** Figures represent the crystal structure of precursor SLP-50 (green) and designed compounds 960, 662 in obtained crystal structures (purple) compared to the predicted with the pipeline (cyan). Predicted compounds 960 and 662 were designed to target the acetyl-lysine binding pocket of BRD4 (PDB ID: 7WL4). **D)** Predicted binding modes for designed ligands 192, 474 in the acetyl-lysine binding pocket of BRD4 (PDB ID: 7WL4). **E)** Interaction map shows contacts for selected compounds compared to SLP-50. **F)** Tanimoto histogram and RMSD plots indicate chemical diversity and pose recovery, respectively.

Experimental validation confirmed that 13 out of 28 compounds showed binding to BRD4 at 2 mM concentration (Figure S15), with positive control JQ1 compound measurements as a reference (Figure 6B, supp. Table 9). Experimental binding affinity of it to the BrD4 target was around 200 nM (the literature value is 50-100 nM [41]). Further specificity analysis against unrelated proteins (KRAS and PIN1) showed that two of these hits exhibited selective binding to BRD4 with minimal off-target interactions, reinforcing their potential as selective ligands. Compounds 960 and 662 were successfully co-crystallized with BRD4 at high resolution, enabling a direct comparison between the computationally predicted poses and experimental structures (Figure 6C, S16, S17). The RMSDs between the crystallographic and predicted poses were 3.1 Å and 2.0 Å, respectively. In both cases, the core fragment was placed deep within the binding pocket aligned closely with the computational prediction, while the peripheral fragment exhibited positional shifts which resulted in larger RMSD from the computational predictions. In addition crystallographic structures provided valuable insight into the binding interactions. Notably, compound 662 formed water-mediated hydrogen bond bridges with four amino acid residues, three of which were novel in comparison to the reference crystal structure with ligand SLP-50 (Figure 6C). Compound 960 formed two known water-mediated hydrogen bond bridges (Figure S17). Displacement of the other water molecules observed in the binding pocket for 960 and 662 compounds can be used as guidance for the following optimization of compounds. These findings validate both the predictive accuracy and practical value of the surface-guided fragment design approach in delivering synthetically accessible, structurally novel, and experimentally validated BRD4 ligands.

**Table 1.**
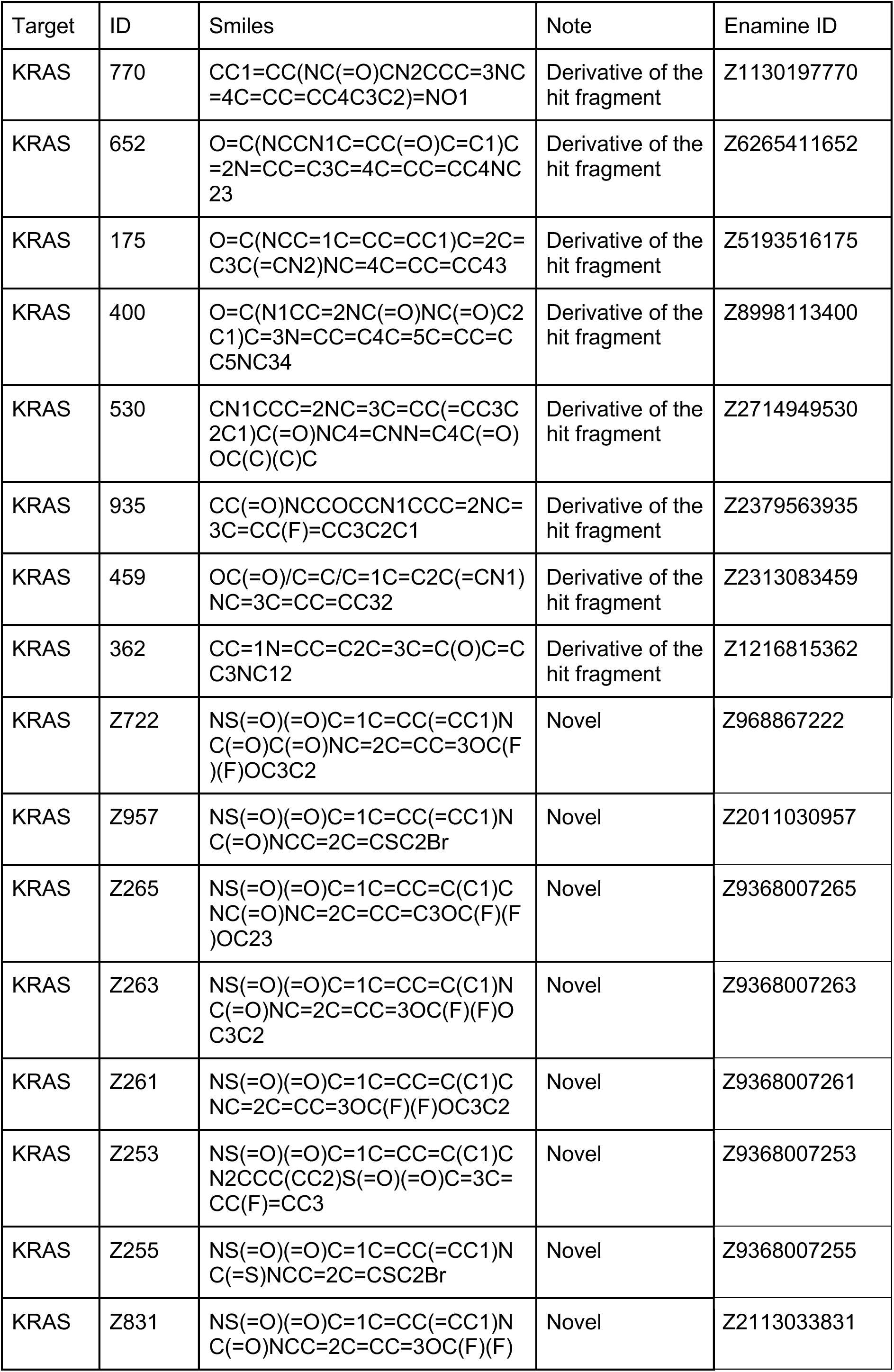
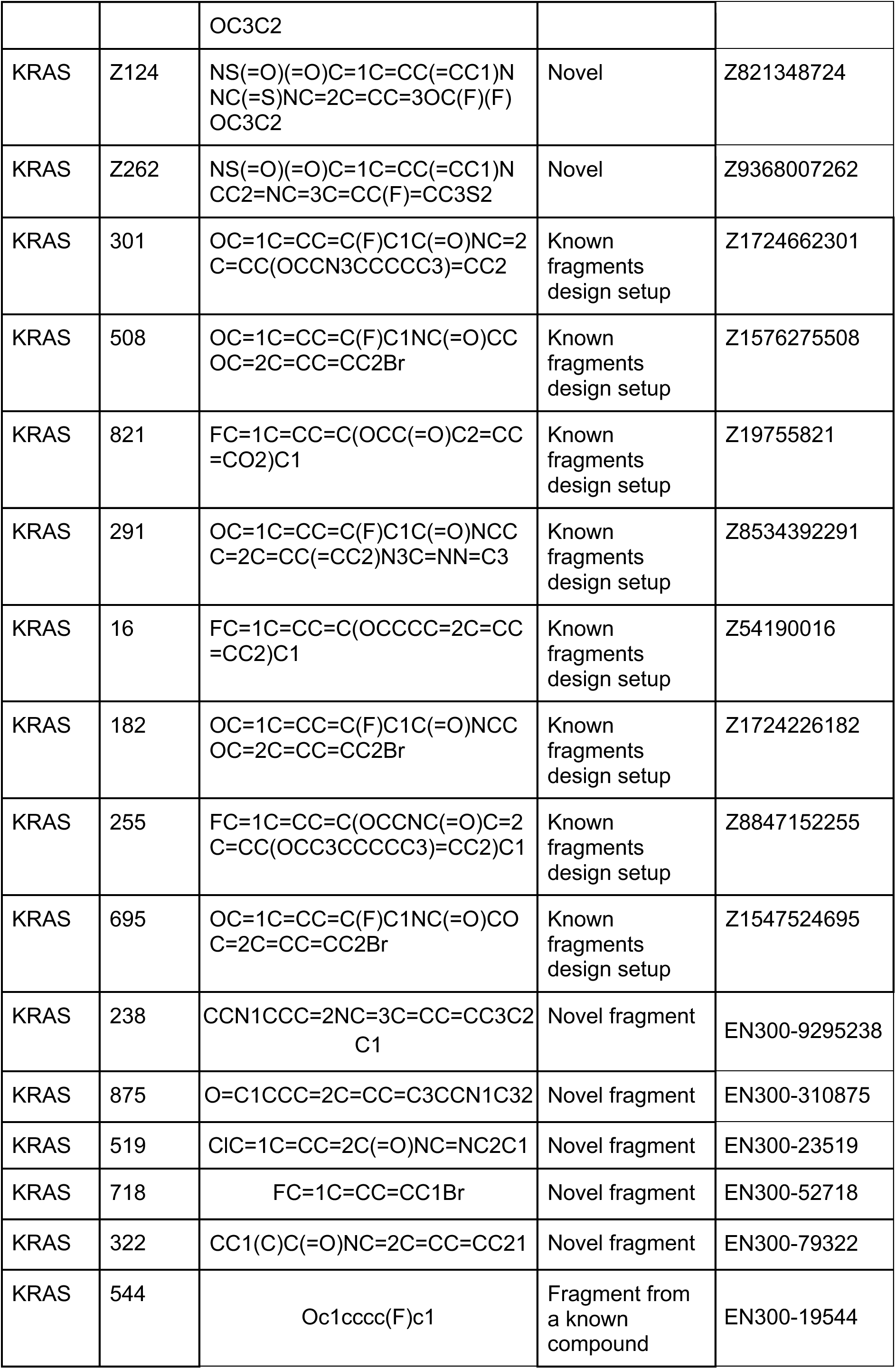
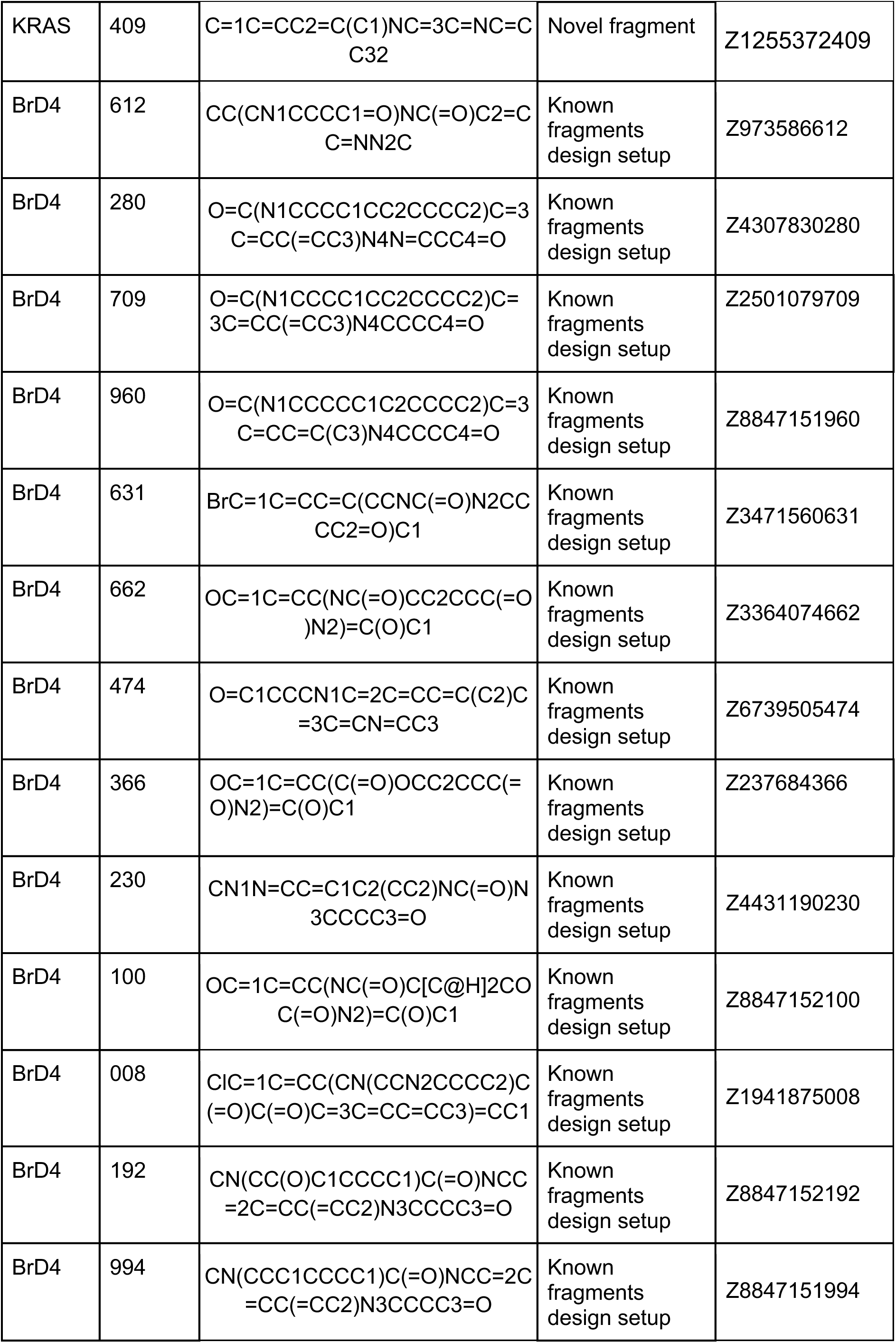
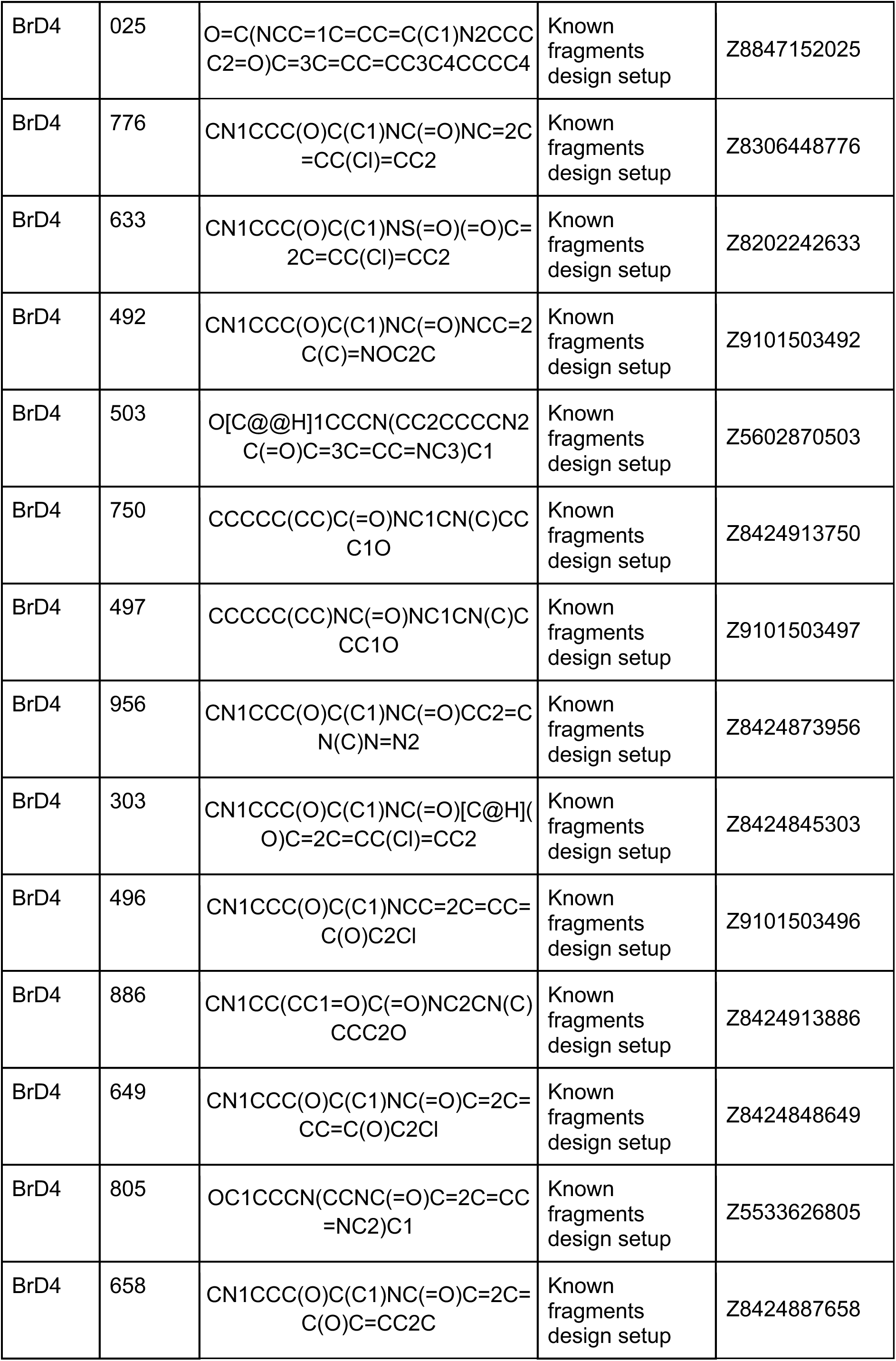
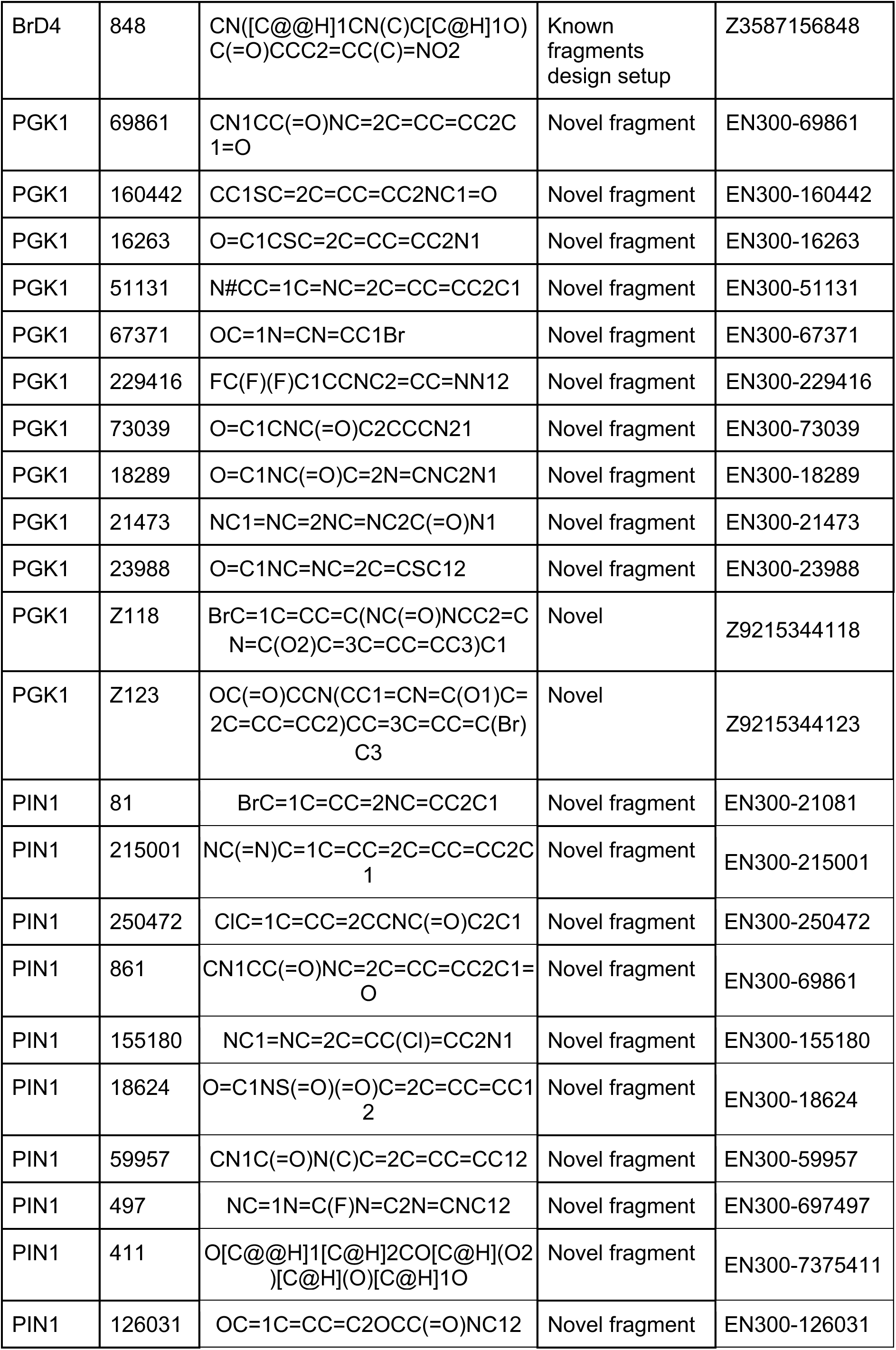
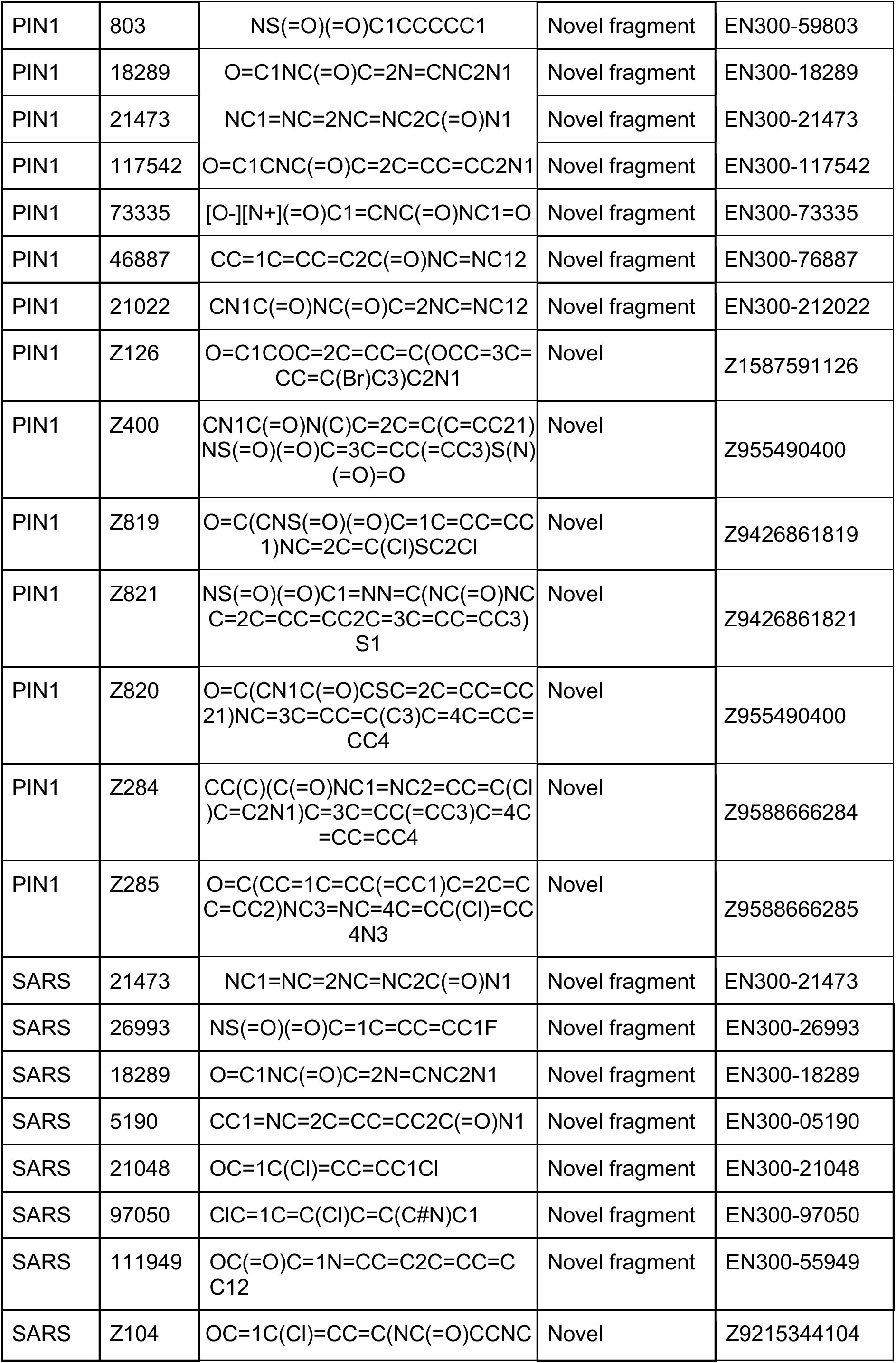
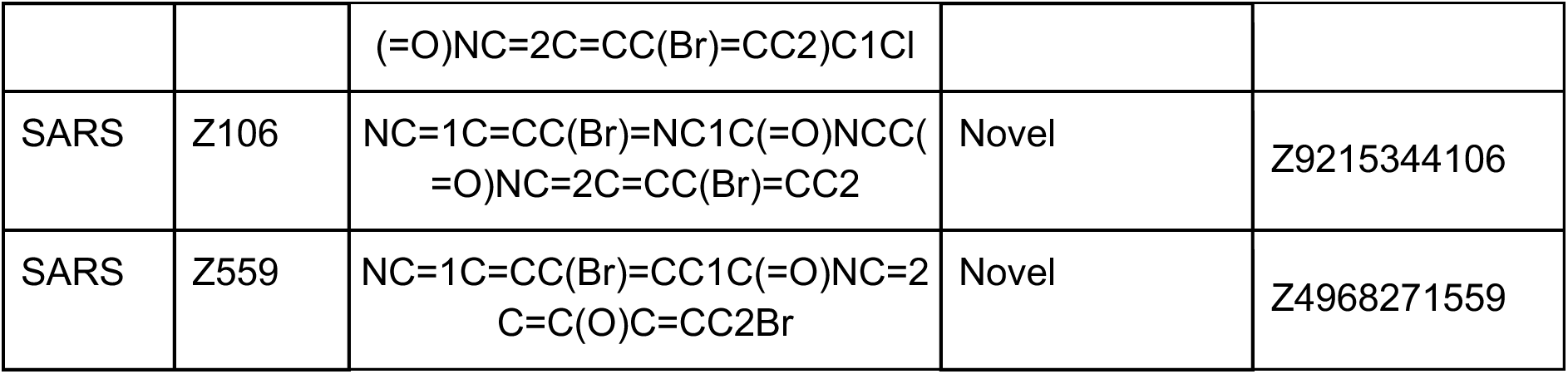
The list of compounds.

**Table 2.**
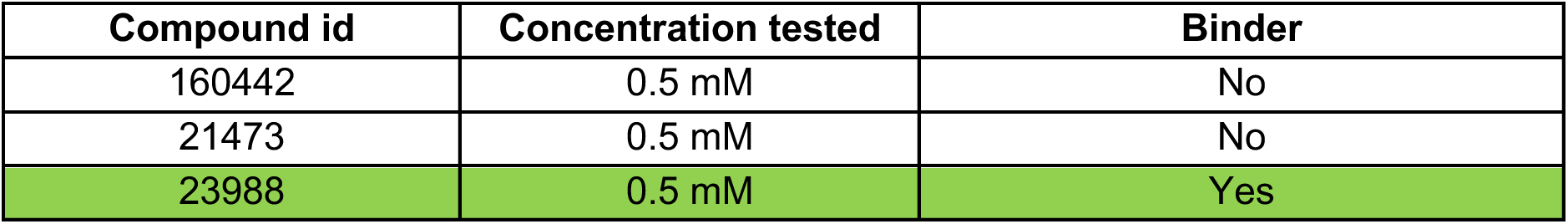
NMR tested PGK1 novel fragments.

**Table 3.**
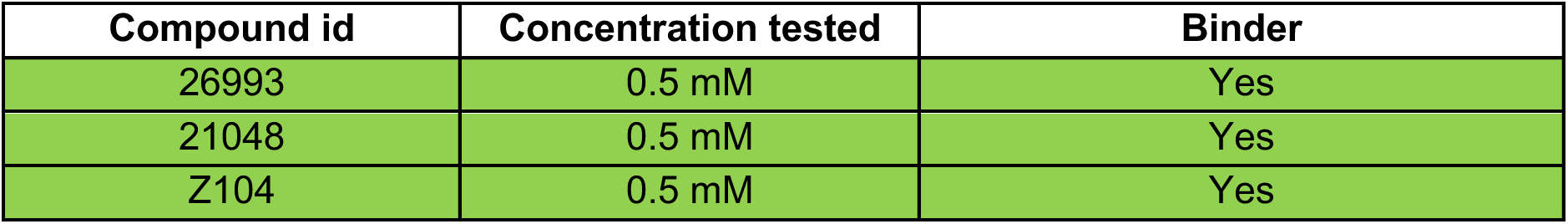
NMR tested SARS Mpro novel fragments and ligands.

**Table 4.**
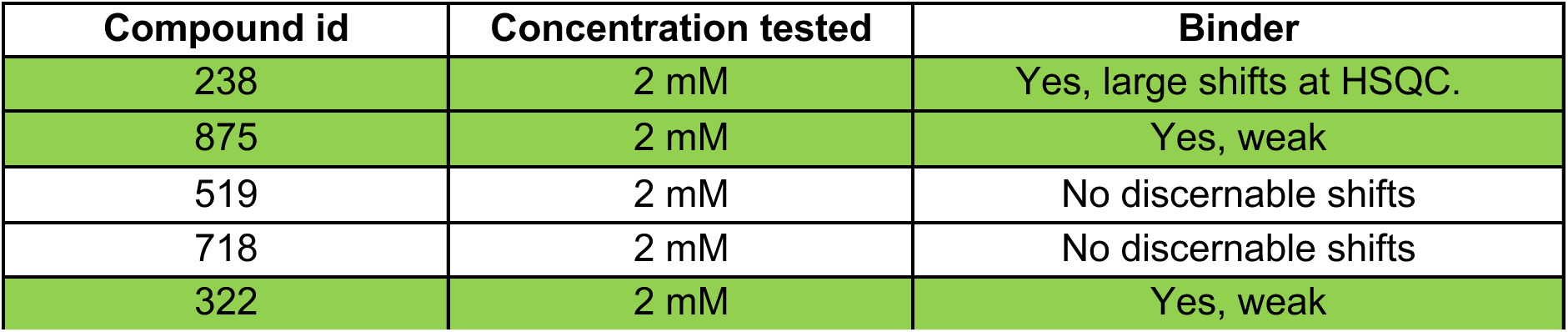
NMR tested KRAS novel fragments.

**Table 5.**
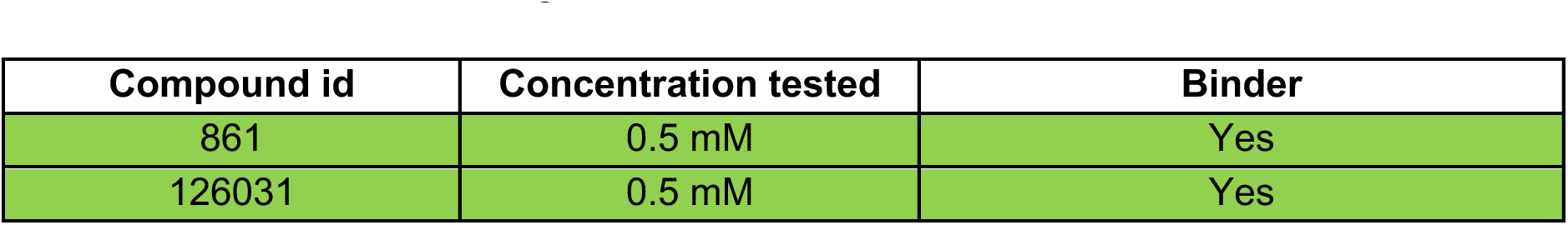
NMR tested PIN1 novel fragments.

**Table 6.**
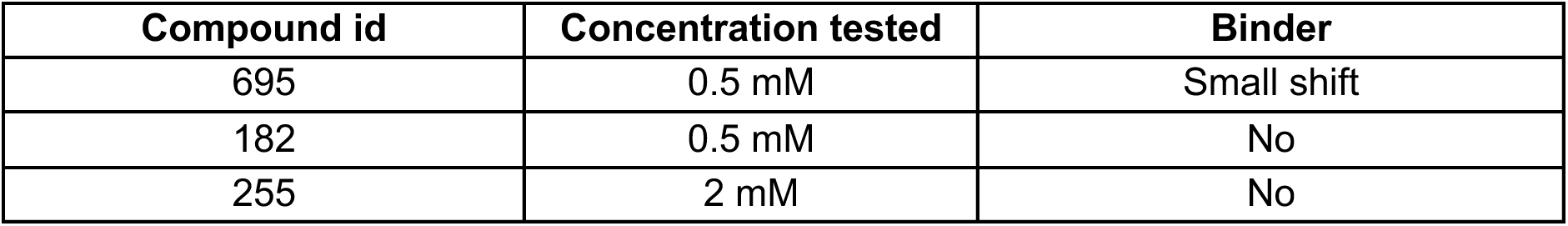
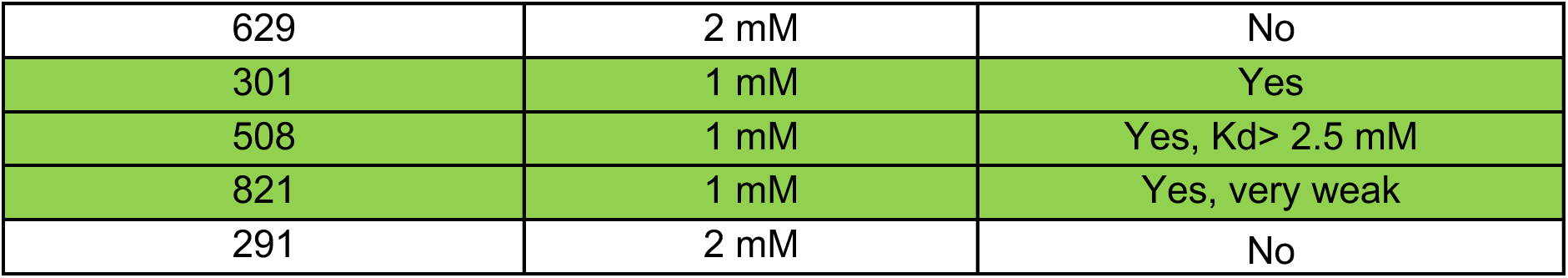
NMR tested KRAS compounds (known fragment, novel linker)

**Table 7.**
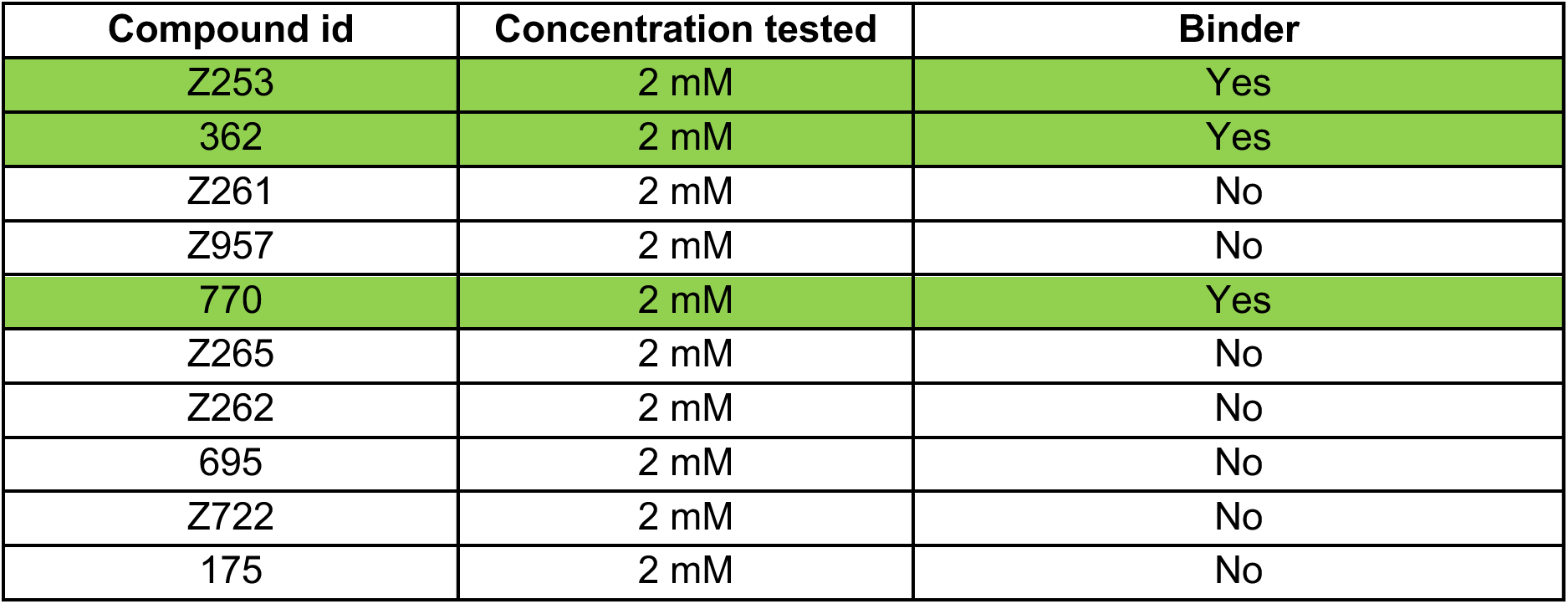
NMR tested KRAS derivatives of novel fragments, de novo molecules.

**Table 8.**
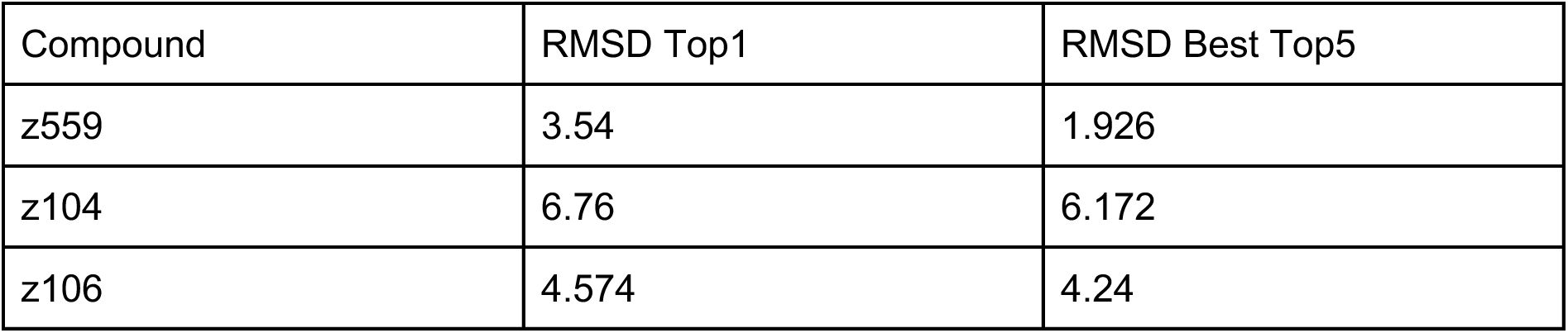
RMSD values for docking of compounds.

| **Compound** | **RMSD Top1** | **RMSD Best Top5** |
| --- | --- | --- |
| z559 | 3.54 | 1.926 |
| z104 | 6.76 | 6.172 |
| z106 | 4.574 | 4.24 |

**Table 9.** Hit criteria table for all targets: KRAS, BrD4, PGK1, SARS Mpro, PIN1. The experimental hits are defined on two criteria: 1) Observed R_max_ > 50% of R_max_ (positive control), 2) dissociation and association errors are ≤ 100%. The relative diss. and ass. errors are calculated as the 95% confidence intervals of the dissociation rate (K_D_) and association rate (K_A_) estimates divided by their values.

| Target | Compound | Rmax | Ass. Error | Diss. Error | HIT? |
| --- | --- | --- | --- | --- | --- |
| KRAS | Z957 | 152 | 27 | 7 | YES |
| KRAS | 770 | 129 | 21 | 6 | YES |
| KRAS | Z255 | 75 | 14 | 5 | YES |
| KRAS | 175 | 0 | 480 | $2.2 \times 10^8$ | NO |
| KRAS | BI-2865 | 23 | 16 | 20 | YES |
| KRAS | 238 | 212 | 34 | 22 | YES |
| KRAS | 409 | 180 | 21 | 6 | YES |
| KRAS | 322 | 34 | 25 | 9.8 | YES |
| KRAS | 718 | 6 | 63 | $5.2 \times 10^7$ | NO |
| BrD4 | 497 | 44 | 17 | 93 | YES |
| BrD4 | 960 | 28 | 4.6 | 11 | YES |
| BrD4 | 662 | 7.6 | 0 | 0 | YES |
| BrD4 | 956 | 0 | $6.2 \times 10^3$ | $4 \times 10^8$ | NO |
| BrD4 | +JQ1 | 14 | 0 | 0 | YES |
| BrD4 | 303 | 116 | 16 | 10 | YES |
| BrD4 | 633 | 122 | 48 | 17 | YES |
| BrD4 | 658 | 47 | 17 | 16 | YES |
| BrD4 | 776 | 353 | 29 | 6.7 | YES |
| BrD4 | 848 | 1 | $1.1 \times 10^3$ | $8 \times 10^7$ | NO |
| PGK1 | Z123 | 403 | 43 | 14 | YES |
| PGK1 | 442 | 34 | 36 | 9.8 | YES |
| PGK1 | 51131 | 22 | 74 | 24 | YES |
| PGK1 | CHR-6494 | 12 | 16 | 5.1 | YES |
| PGK1 | 69861 | 87 | 75 | 11 | YES |
| PGK1 | Z118 | 0 | $1.1 \times 10^3$ | 52 | NO |
| SARS | 26993 | 23 | 25 | 11 | YES |
| SARS | 5190 | 32 | 25 | 34 | YES |
| SARS | 21048 | 102 | 47 | 12 | YES |
| SARS | 97050 | 22 | 27 | 7.1 | YES |
| SARS | 18289 | 0 | $5.9 \times 10^3$ | $9.9 \times 10^2$ | NO |
| SARS | Nirmatrelvir | 79 | 42 | 42 | YES |
| SARS | Z104 | 20 | 68 | 32 | YES |
| PIN1 | 861 | 48 | 23 | 29 | YES |
| PIN1 | 73335 | 20 | 78 | 26 | YES |
| PIN1 | 803 | 4 | $1.3 \times 10^3$ | $4.5 \times 10^2$ | NO |
| PIN1 | 18624 | 22 | 55 | 18 | YES |
| PIN1 | DTM | 3.2 | 39 | 52 | YES |
| PIN1 | 126031 | 9 | 76 | 89 | YES |
| PIN1 | Z820 | 0 | $1.1 \times 10^3$ | 52 | NO |

Interaction analysis of other designed compounds such as 192, 474 also revealed the recovery of some reference interactions SLP-50 such as a hydrogen bond with ASN140. However, the overall profile is not as rich as for the reference compound (Figure 6D,E). Notably, all designed compounds are chemically diverse, spanning novel scaffolds distinct from SLP-50 or any other known ligands bound to the target pocket of BRD4 (Figure 6F, histogram), showing novelty in the generative workflow. In addition, we performed docking analysis using GNINA to computationally validate the predicted binding modes (Figure 6F). The median RMSD between the predicted and docked poses was 4.74 Å for the top-ranked pose, and 3.7 Å when considering the best pose among the top five, which is within the range of the RMSDs observed in crystal structures.

Overall, presented results highlight the power of the FragmentScope, it efficiently explores and narrows the chemical space, offering novel fragments placement that can be successfully leveraged for small-molecule design. The data we present demonstrate the novelty of the designed compounds, importantly we experimentally confirmed their predicted binding poses. Together, these data underscore the potential of our method to enhance fragment-based drug discovery.

## Discussion

Fragment-based drug discovery is a promising strategy for hit identification and optimization. It offers several key advantages such as access to vast chemical spaces through fragment combination and higher ligand efficiency compared with traditional screening libraries. However, experimentally detecting low molecular weight fragment hits is challenging due to their low binding affinity. With the advent of highly accurate deep learning prediction methods, computational approaches that can provide a means for fast and cost-effective fragment pre-selection, the experimental success rate can increase significantly.

Popular deep learning approaches typically hinge on scaling up neural network architectures enabled by increasingly large training datasets, a strategy that has so far led to limited success on protein-ligand interaction prediction tasks [43]. Small molecule interactions are difficult to predict reliably as even the smallest atomic modifications can change the energetic equilibrium from the bound to unbound state. Moreover, deep learning approaches struggle to generalize to unseen chemistry as only about 27,000 solved structures are currently available for training in the PDB [44]. Previous efforts were dedicated to create a database of crystallized protein-fragment complexes, however these data are still limited to the amount of structures [45,46].

To enhance generalisation when insufficient data is available for training large models, we present a deep learning approach grounded in geometric and physical principles. FragmentScope utilizes a protein surface representation and learns complementarity-determining features of surface patches directly in contact with a ligand. This approach reduces the complexity of the molecular system and limits computations to areas that immediately affect the intermolecular interaction. For fragment placement, FragmentScope performs a database search based on the learned features, which adds another layer of robustness as small deviations of predicted embeddings get mapped to the closest known interaction pattern from the database.

Our evaluation showed that FragmentScope consistently outperformed traditional docking methods in placing fragments accurately within known binding pockets. This accuracy was achieved due to the ability to leverage structural similarity to previously characterized protein-ligand interfaces, enabling the placement based on structural context captured by computational scoring metrics.

The key advantage of FragmentScope lies in its ability to efficiently prioritize fragments and narrow down the screened chemical space. Instead of experimentally testing thousands of fragments, our method identifies structurally relevant fragments based on learned surface similarities and places them in the pocket of interest. These computationally selected fragments act as anchor points and could provide structurally meaningful starting scaffolds for subsequent molecular design. The average success rate of FragmentScope in our study was 59% across 4 targets, which improves upon random fragment screening, which is typically reported to be 5-10% for NMR screens, depending on the target [10]. Slightly higher success rates can be expected for campaign-level hit rate identification on large-scale platforms like XChem which is in the range of 1-30% [46]. Combined approaches with computational pre-selection and following experimental validation like the SEED2XR protocol have shown hit rates of 10-40% [47,48]. Thus, FragmentScope achieves highly competitive results, underscoring its potential in narrowing down the chemical space and identifying hits.

Transitioning from fragment hits to full molecules, we computationally grew identified fragments. Our pipeline was proved to be effective, yielding experimental hit rates of 30 - 40%. While this approach inherently limits chemical novelty as it depends on a limited amount of building blocks, it still produces molecules with diverse scaffolds and low Tanimoto similarity to known binders, including fully novel compounds.

Two orthogonal methods for hit validation: GCI technology and NMR spectroscopy, different in their biophysical principle, confirmed the hits designed by the Fragmentscope method. However, as anticipated, not all could be confirmed - likely due to low binding affinities falling below the detection thresholds of the respective techniques. However, both methods demonstrated a higher success rate in validation of fragments rather than ligands for KRAS, PGK1, PIN1, SARS protein targets. K_D_ determination using the GCI method was approached with caution due to limitations in achieving binding curve saturation and the accuracy of kinetic model fitting. Hit fragments validated by two orthogonal methods: GCI and NMR can serve as effective scaffolds for the further design of high-affinity ligands.

Despite many encouraging results, our data also highlights some remaining limitations. Notably, the PIN1 experiments show that flat and charged pockets are challenging screening targets. FragmentScope’s effectiveness is inherently dependent on the diversity and scale of the underlying fragment-surface database, which was constructed from co-crystallized protein-ligand pairs. Therefore, growing this virtual screening library as more crystal structures become available will expand and diversify the repertoire of available pharmacophore patterns, and thus increase the chances of finding highly complementary fragments even for the most challenging targets.

Additionally, FragmentScope, like most fragment-based discovery approaches, struggles to achieve high levels of specificity. Future work could address this limitation by incorporating explicit negative design to select fragments not only for their complementarity to the target pocket but also for incompatibility with off-target protein surfaces. Growing the compounds, optimising them with additional chemical groups to the target pocket is another way to improve the specificity of the compounds. Finally, crystallographic and docking-based interaction profiles revealed a lack of strong enthalpic interactions in most designed compounds, an expected outcome at this early stage. However, we observed important stabilizing features, such as water-mediated hydrogen bonds, which often play key roles in fragment retention and optimization [49,50]. For instance, co-crystallized compound 662 with BRD4 formed more water bridges and hydrophobic contacts than the reference ligand SLP-50, despite its smaller size. These results suggest that fragment-guided placement, when integrated with accurate solvent modeling, could better exploit interaction modes in small molecules design.

In conclusion, FragmentScope addresses key bottlenecks in FBDD by enabling structure-guided, surface-based fragment selection. While further optimization is required to reach drug-like affinities, our approach reliably identifies novel, synthetically accessible, and selectively binding scaffolds, significantly accelerating the path from protein target pocket to chemical compound.

## Methods

### Protein-ligand encoder

To ensure meaningful and expressive protein surface descriptors, we train a protein-ligand encoder (PLE) in a contrastive learning fashion using a curated dataset of protein-ligand complexes from Protein Data Bank (PDB) [26].

To curate the dataset, we downloaded 122,012 raw X-ray, Cryo-EM and NMR structures of protein-ligand complexes from PDB with resolution below 3Å and release date before the 1st of May, 2023. We then split all proteins in chains and generated ligand-protein chain pairs resulting in 244,244 data points. We used the Ligand Expo database [51] to assign covalent bond types to all ligands and filtered out the ones from which we could not match the chemical structures with the corresponding Ligand Expo representatives. Additionally, we removed all molecules that did not pass the RDKit [52] sanitization procedure. Besides, we filtered out all ligands that were labeled as invalid in the latest BindingMOAD release [53]. Finally, we curated our own additional list of compounds that were abundant in our collection but yet of little interest (typically sugars, organophosphates and fatty acids). We also excluded pairs in which less than 50% of ligand heavy atoms are in contact (i.e. within 5Å) with the protein chain. As a result, we reduced the number of available protein-ligand pairs to 118,308. Next, we downloaded the precomputed MMseqs2 [54] 30% identity cluster for all PDB chains and used them to split the data in the train (n=97,129), validation (n=6,954) and test (n=10,831) sets. Due to the high redundancy of certain protein families, at every training epoch we iterated over available training clusters (n=4,752), and for each of them sampled one protein-ligand pair per epoch. This allowed us to train the model in a balanced way, while using all available data.

The protein-ligand encoder consists of two separate neural networks that independently operate on proteins and small molecules. For proteins, we first triangulate the surface mesh using MSMS [55] and then learn surface features by mapping types and distances of nearby atoms to every surface point (i.e. mesh vertex), followed by approximately geodesic convolutions, using the dMaSIF architecture [56]. Ligands, on the other hand, are encoded with a graph neural network (GNN) directly operating on atom types and bond types. Both surface vertices and ligand nodes are encoded in a shared 16-dimensional latent space where we can compute similarities for pairs of descriptors.

The training objective for both neural components is to increase the cosine similarity between embeddings of interacting (positive) pairs while reducing the similarity between non-interacting (negative) pairs. To make the latent space more dense and compact, we additionally regularise the norm of the embedding vectors.

Positive pairs are defined based on a distance cutoff of 3 Å. To construct a diverse and balanced set of negative pairs (equal in size to the positive set), we associate ligand node descriptors with the protein surface points randomly selected from three geometrically-distinct regions of the protein surface: pocket, non-pocket concave (i.e. pocket-like) areas, and non-pocket convex areas. To assemble negative examples using the points from the pocket itself, we consider pairs where the protein and ligand points are separated by more than 3 Å. For the non-pocket concave (i.e. pocket-like) and non-pocket convex categories, we train a gradient boosting pocket classifier on a random subset of 1,000 training protein surfaces. Each surface point is represented by a five-dimensional feature vector of curvature values computed at scales 1.0, 3.0, 5.0, 7.0, and 9.0, as in dMaSIF [56], and the model is trained to detect surface points belonging to the known pocket (based on the 3 Å distance from the ligand). The trained gradient boosting model is then used to classify all surface points as either concave or convex, enabling the selection of negative pairs from both regions to ensure geometric diversity. Overall, at each training step and for each training protein-ligand complex, we collect all available positive pairs of points, and then sample the same number of negative pairs with equal populations in all three categories.

Overall, PLE consists of 118K trainable parameters. We trained the model for 35 epochs using the mini-batch stochastic gradient descent algorithm with Adam optimizer, batch size of 32, lambda=0.1, and learning rate of 0.001. The training was performed on a single GPU NVIDIA Tesla V100-PCIE-32GB and took 3 days.

### FragmentScope

FragmentScope consists of a curated database of pocket-fragment pairs and a search algorithm that returns a ranked list of the most relevant fragments placed around the pocket of interest to maximize protein-fragment interactions.

To find the most relevant fragments for an input protein (with a known approximate query pocket), we first process the protein with PLE and retrieve the surface point cloud corresponding to the pocket. Next, we search for the most similar patches in the database. The general scheme of the search procedure is represented in Figure S2 and consists of three main steps: shortlisting, alignment and scoring. First, we perform a search over the whole library in order to shortlist candidate ligands. To reduce computational complexity and focus the search on relevant candidates, we implemented a filtering step that selects the top 200 fragment-pocket matches for each query pocket based on global surface similarity. The filtering process proceeds as follows: we first compute pairwise distances between the query pocket embedding and all candidate patch embeddings. For each candidate patch, we determine the minimum distance from each patch point to the query pocket, capturing the closest structural correspondence. These minimal distances are then averaged into a final similarity score for ranking. This scoring method effectively prioritizes patches that contain regions most structurally similar to the query pocket, without requiring a full alignment at this stage. In the second step, we align pockets of shortlisted candidates with the query pocket using Random Sample Consensus (RANSAC) [57] followed by point-to-point Iterative Closest Point (ICP) [58]. Once the shortlisted pockets are aligned with the query pocket, we rank pairs of found patches and corresponding fragments based on the recomputed similarities between surface patches. To do this, we assemble pairs of the closest surface points from the query and found patches and compute Euclidean distances between the corresponding descriptors. The final score is obtained by averaging the computed distances over all collected pairs.

### Databases

Since the FragmentScope is an engine that operates on the pre-built database. Two databases were constructed with the above described procedure. The first database was designed with the fragments of MW 150-500 Da with the total number of fragments-patch pairs n=129,481. The search across this database was aimed to find rather big fragments able to be good scaffolds for the future compounds as well as have more signal for the binding affinity. Fragments obtained from this database were tested experimentally and described in the ‘Prospective fragments screen’ Results section. The second database was created with the aim to place fragments with the following pairing and linking procedure to get full molecules. The molecular mass of the fragments was 60-300Da with the total number of fragments-patch pairs n=475,190. Designed by using the second database of compounds are described in the following ‘Fragment-based small molecules design section’ of Results.

Both databases were constructed on the complexes from a curated Protein Data Bank dataset. The filtering and selection of the complexes was conducted with the same filtering procedure described in the ‘Protein-Ligand encoder’ part Methods section.

### Grating coupled interferometry

Grating coupled interferometry method was chosen for hit identification. The binding kinetics was measured by label-free, GCI (grating-coupled interferometry) on the Creoptix WAVE system (Malvern Panalytical) using Creoptix WAVE control software (Malvern Panalytical, v.4.5.18). Protein samples were immobilised through amine coupling (carboxymethyl 5-dextran on the 4PCH chip (Malvern Panalytical). Activation of a chip surface was performed with injection of a 1:1 mixture of 100 mM *N*-hydroxysuccinimide and 400 mM 1-ethyl-3-(3-dimethylaminopropyl)-carbodiimide. All protein samples were immobilised on a chip surface in a concentration of 40 ug/ml in 10 mM of sodium acetate pH 4.5 over a 6 min period of injection. The remaining activated groups have been passivated with 1 M of ethanolamine pH 8.5 over 6 min injection. The level of protein immobilisation was 10000-15000 response units (RU).

The running buffer consisted of HPS-P+ buffer (10 mM HEPES pH 7.4, 150 mM NaCl, 0.005% v/v surfactant P20; 1 %DMSO GE Healthcare), for KRAS protein additionally contained 5 mM MgCI2 and 1 mM DTT (also for SARS, PIN1). The screening for hits was done at 2 mM and 500 µM concentrations in the following running buffer: 20 mM HEPES, 150 mM NaCI, 1 mM DTT, 0,02% P020 surfactant, 1% DMSO at 25 C. Positive controls binding was checked firstly to estimate the amount of active protein on a chip, the following positive controls KRAS: BI-2865, PGK1: CHR-6494, PIN1: DTM, BrD4: +JQ, SARS Mpro: Nirmatrelvir were chosen. The waveRAPID (repeated analyte pulses of increasing duration) kinetic assay [59] for hit screening (flow rate 400 ul/min, 5 s association time, 20 s dissociation time) was applied, in this type of kinetic all compounds have been injected at the same concentration of 2 mM or 500 uM. We applied the following filtering hit identification criteria: 1) Observed R_max_> 50% of R_max_ (positive control), 2) dissociation and association errors are ≤ 100%. The relative diss. and ass. errors are calculated as the 95% confidence intervals of the dissociation rate (K_d_) and association rate (K_a_) estimates divided by their values. WaveRAPID measurements were fitted with a 1:1 kinetic model to define affinity range.

### Protein production and purification

A list of protein sequences can be found in Supplementary Table 6. Genes encoding the 6xHis-tagged proteins of interest were purchased from Twist Bioscience, cloned into pET11 (bacterial vector) using NDeI and BlpI restriction sites by Gibson assembly, only BrD4 plasmid was kanamycin resistant - pNIC28-Bsa4 Kanr and transformed into XL10-Gold or HB101 bacteria. Plasmids were extracted using a GeneJET plasmid Miniprep kit (Thermo Fisher, for bacterial vector) and sequenced by Sanger. All Proteins were purified using bacterial expression systems. For bacterial expression, BL21(DE3) or T7 Express Competent *Escherichia coli* were transformed with the plasmid of interest and grown as a preculture overnight. Precultures were inoculated 1:50 in Terrific Broth medium and incubated at 37 °C until they reached an optical density at 600 nm (OD600) of approximately 0.6. The differences were done for BrD4 protein induction; it was induced with 1 mM kanamycin, when the optical density was 3.0 (OD600). Then, bacteria were induced with 1 mM isopropyl β-D-1-thiogalactopyranoside (IPTG) and incubated overnight at 18–20 °C. Cells were collected by centrifugation at 4,000*g* for 10 min, resuspended in lysis buffer (50 mM Tris, pH 7.5, 500 mM NaCl, 5% glycerol, 1 mg/ml lysozyme, 1 mM phenylmethylsulfonyl fluoride (PMSF) and 1 µg/ml DNase) and lysed by sonication using 30s pulses on power setting 16 with 1 min pauses on ice between pulses. Some additional reagents in lysis buffer for KRAS and BrD4 proteins were added, which contained additionally 5 mM MgCI_2_ and one EDTA-free protease inhibitor cocktail tablet (Roche Diagnostics, Mannheim, Germany). Lysates were then clarified by centrifugation at 30,000*g* for 30 min and filtered.

All 6xHis-tagged proteins were purified using an ÄKTA Pure system (GE Healthcare) Ni-NTA HisTrap affinity column, followed by size-exclusion chromatography on a Superdex HiLoad 16/600 75 pg or 200 pg depending on the size of the protein. With additional steps for SARS-CoV-2 Main Protease purification - buffer exchange on PD MidiTrap G-10 columns and with the next purification on a HiTrap Q FF column (GE Healthcare). All proteins were concentrated in the appropriate final buffer for every protein: KRAS buffer - 20 mM HEPES, 150 mM NaCI, 1 mM DTT, 5 mM MgCI_2_, BrD4 buffer - 20 mM HEPES, 500 mM NaCI, 5% glycerol, SARS, PIN1 buffer - 20 mM Tris, pH 7.5, 150 mM NaCI, 1 mM DTT and flash frozen with liquid nitrogen and frozen at −80 °C. The PGK1 protein sample was provided by a collaborator from BioNMR laboratory. Full protein amino acid sequences (see supp. Table 10).

**Table 10.**
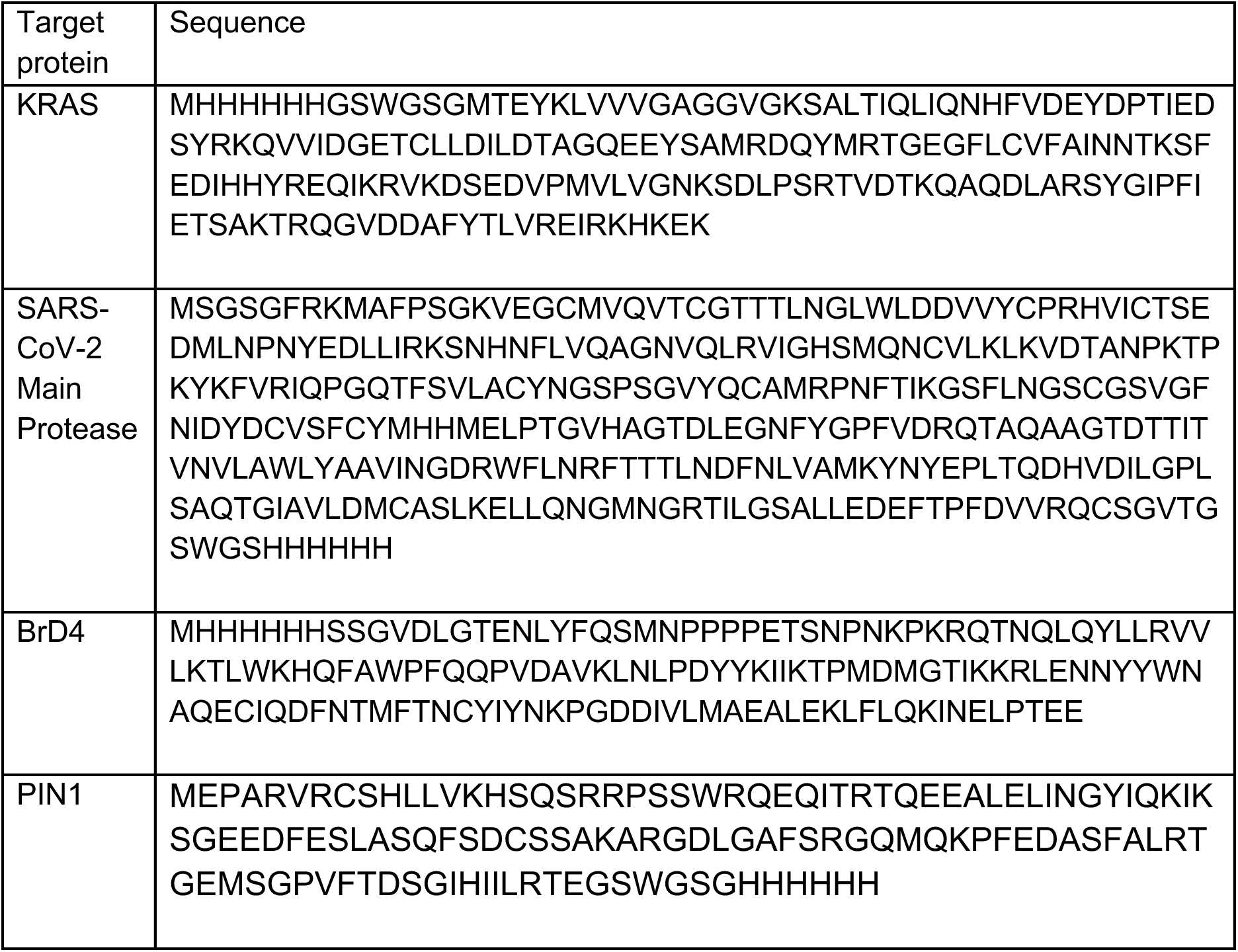
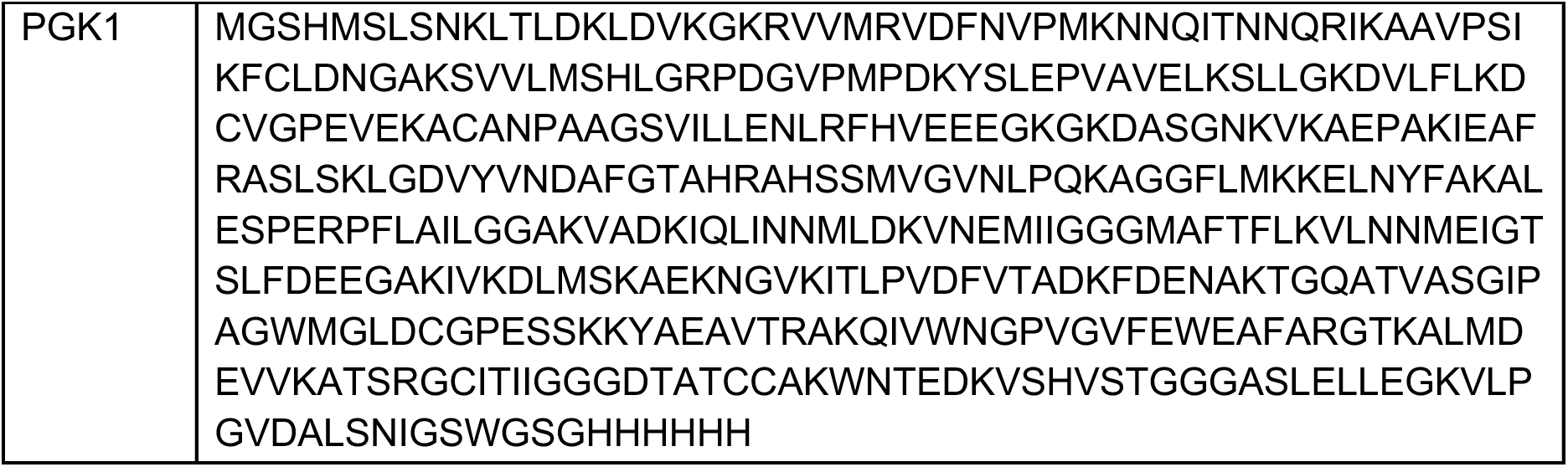
Target protein and sequence.

**Table 11.**
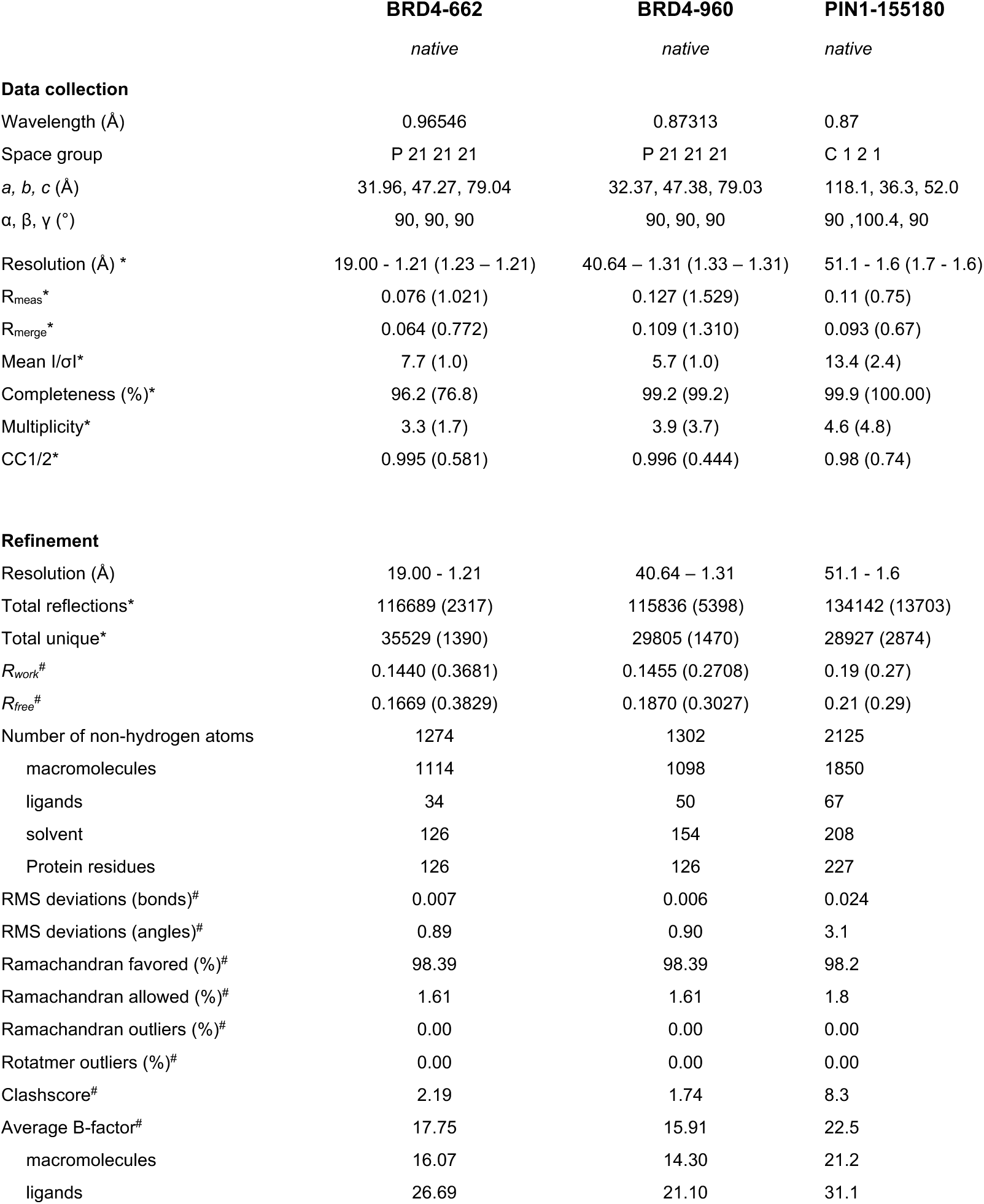
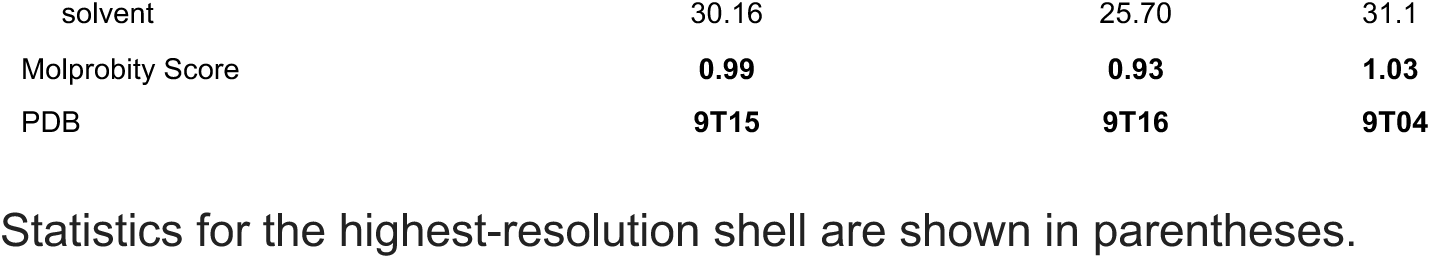
Data collection and refinement statistics.

### NMR

#### KRAS WT and PGK1 G185A protein expression and purification for NMR

A preculture was grown for 5 hours at 37 °C, 110 rpm in 1x Luria Broth with kanamycin and transferred to M9 Minimal Media: 1 L (3 g KH_2_PO_4_, 8.4 g Na_2_HPO_4_ · 2H_2_O, 0.5 g NaCl, 7 g D-glucose, 1 g ^15^NH_4_Cl, 10 mL MEM vitamin mix, 1 mL kanamycin, 2 mL MgSO_4_ (1 M)). Protein expression was induced at OD600 was 0.4 by addition of 1 mM IPTG with overnight incubation at 25 °C. Protein was harvested by spinning down the cells at 4000 g, 20 min at 4 °C and resuspending the cell pellet in 35 mL of lysis buffer: 50 mM Tris, 500 mM NaCI, 2 mM MgCI_2_, and one EDTA-free protease inhibitor cocktail tablet (Roche Diagnostics, Mannheim, Germany). The cells were lysed by passing them 3 times through a microfluidizer. The cell lysate was centrifuged for 20 min at 35000 g at 4 °C. The resulting supernatant was filtered before loading onto a 5 mL HisTrap FF Ni-column (GE Healthcare), equilibrated with Buffer A: 50 mM Tris, 500 mM NaCl, 5 mM MgCI_2_, 10mM imidazole and 2 mM BME, pH=7.4 with a flow rate of 2 mL/min with a diastolic pump. Once loaded with protein, the column was washed with 50 mL of Buffer A and the protein was eluted with an imidazole concentration of 150 mM. Dialysis was done in 20 mM Tris, 50 mM NaCl, 1 mM MgCI_2_, 2 mM BME pH=8 overnight to get rid of the imidazole and the His-tag was cleaved with TEV protease over 2 days at 4°C. The protein solution was filtered and passed over the His-Trap column in order to remove the tag, and concentrated in a 10 kDa concentrator (Merck Millipore Amicon® Ultra). A PD10 column was used to change to an NMR buffer: 20 mM HEPES, 100 mM NaCl, 5 mM MgCI_2_, 2 mM TCEP, 10 % D_2_O.

#### NMR experiments

All NMR experiments were performed on a Bruker Avance III HD 600 MHz spectrometer equipped with a TCI-cryoprobe and SampleXpress. The [^15^N, ^1^H]-HSQC experiments for determining affinities were measured with 128 (*t*_1, max_ (^15^N) = 52.6 ms) × 1024 (*t*_2, max_ (^1^H) = 122 ms) complex points data points with 8 scans per increment and 0.8 s interscan delay on a Bruker Avance IIII HD 600.

The ^15^N KRAS G12V concentration was 100 μM. The compounds were titrated at a concentration of 250, 500, 1000, 1500 and 2000 μM. Where no kd was determined a ligand concentration of 2 mM was used. The ^15^N PGK1 G185A concentration was 100 μM and the ligand concentration was 0.5 mM.

### Crystallization

#### BRD4

The BrD4-compound complexes (8 mg/ml) were crystallised using the sitting-drop vapour diffusion setup at 18 C with 1:1 ratio of BrD4 compound sample and crystallisation solution containing sodium formate 5-6 M, glycerol 10-12%. The protein was first mixed with 100 mM of compound in 1.3% final DMSO. Crystals were cryoprotected with 25% glycerol and flash-cooled in liquid nitrogen. Diffraction data were collected at a temperature of 100 K at the European Synchrotron Radiation Facility at beamlines MASSIF-1 and ID30B(ESRF Grenoble, France) [37]. Raw data were processed and scaled with XDS (VERSION Jan 19, 2025 BUILT=20250714) [37] and then processed using the autoPROC package [60],[61]. (GlobalPhasing, v.20250717).

Phases were obtained by molecular replacement using the Phaser module of the Phenix package (v.2.83) [61] and a model from PDB ID: 8B98. Manual model building and refinement were completed using COOT (v.0.9.8.96 EL) [62,63] and Phenix.refine [62,63] (v.2.0-5761). Details of the data collection and refinement statistics are shown in the Extended Data Table 11.

#### PIN1

##### Protein Expression, Purification, and Co-crystallization

The open reading frame of the *PIN1* PPiC domain containing the K77Q and K82Q mutations was subcloned into the pET-28a(+) vector (GenScript), which included an N-terminal 6×His tag and a thrombin cleavage site. The construct was transformed into *E. coli* BL21(DE3) cells (Novagen) using a standard heat shock protocol.

Transformed cells were plated on 2xTY agar supplemented with 50 µg/mL kanamycin and incubated at 37°C for 16 hours. A colony lawn was scraped and used to inoculate 1 L of sterile 2xTY liquid medium containing 50 µg/mL kanamycin. Cultures were grown at 37°C with vigorous shaking until reaching an optical density at 600 nm (OD₆₀₀) of 0.6. The culture was then cooled rapidly under tap water, and protein expression was induced by the addition of 0.5 mM IPTG (isopropyl β-D-1-thiogalactopyranoside). Expression continued for 16 hours at 18°C. Cells were harvested by centrifugation at 6,000 × g for 10 minutes at 4°C, washed with PBS, centrifuged again, and the resulting ∼6 g cell pellet was stored at –20°C.

Cell lysis was performed by resuspending the pellet in lysis buffer (50 mM HEPES pH 7.5, 500 mM NaCl, 20 mM imidazole, 1 mM DTT) at a 1:9 (w/v) ratio. An EDTA-free protease inhibitor cocktail (Merck) and DNase I were added. Cells were disrupted using a Sonics Vibra-Cell VC 130 ultrasonic homogenizer (10 cycles of 30 s pulses with 1 min intervals, 45% amplitude). The lysate was clarified by centrifugation at 60,000 × g for 45 minutes at 4°C (Beckman Coulter Avanti J-26 XP centrifuge). The supernatant was incubated with Ni-NTA resin (Qiagen) for 2 hours at 4°C.

The resin was washed with 25 column volumes (CV) of wash buffer (lysis buffer with 40 mM imidazole), and bound protein was eluted using an elution buffer containing 500 mM imidazole. Fractions containing the target protein were identified by SDS-PAGE, pooled, and subjected to thrombin digestion (Merck) for 16 hours at 4°C during dialysis against cleavage buffer (50 mM Tris-HCl pH 8.0, 150 mM NaCl, 10 mM CaCl₂).

The cleaved sample was re-applied to Ni-NTA resin to remove uncleaved protein and His-tagged contaminants. Thrombin was removed using a HiTrap Benzamidine FF column (Cytiva). The protein was concentrated to 5 mL using a 10,000 MWCO Vivaspin 20 centrifugal concentrator (Sartorius) and purified by size-exclusion chromatography on a HiLoad 16/600 Superdex 75 column (Cytiva) equilibrated in gel filtration buffer (10 mM HEPES pH 7.5, 100 mM NaCl, 1 mM DTT). Peak fractions were pooled, analysed by SDS-PAGE, and concentrated to 15 mg/mL.

For crystallization, a hanging-drop vapor diffusion method was employed. Drops consisting of 2 µL of protein solution and 2 µL of reservoir solution (100 mM HEPES pH 7.5, 200 mM ammonium sulphate, 1.1–1.3 M sodium citrate, 5 mM DTT) were equilibrated at 20°C. Crystals of apo PIN1 appeared within 48 hours and were used as seeds for further optimization. Seeded crystals were soaked for 24 hours in reservoir solution supplemented with 4% DMSO and 24 mM of compound 155180.

##### X-ray Data Collection, Structure Determination, and Refinement

Prior to data collection, crystals were cryoprotected by soaking in reservoir solution supplemented with 20% (v/v) glycerol, 5% DMSO, and 24 mM of compound 155. Crystals were flash-cooled in liquid nitrogen and shipped to the European Synchrotron Radiation Facility (ESRF, Grenoble, France). X-ray diffraction data were collected at beamline ID30B [64] using a PIXEL Eiger2_9M detector.

Diffraction data were indexed, integrated, and scaled to a resolution of 1,6 Å using the program AIMLESS within the [65] CCP4i2 software suite. The structure was solved by molecular replacement using Phaser [63] in Phenix, employing the PIN1 structure from PDB entry 4TYO as a search model.

Manual model building was conducted in Coot [66], and structural figures were generated using PyMOL [67]. Data collection and refinement statistics are summarized in Table S11.

### Small molecule design pipeline

To design small molecules based on the placed fragments the following pipeline was developed. In the first step, we search and place a set of fragments within the known target protein pocket using FragmentScope using the second pre-build database. In the second step we create a library comprising all possible pairs of the placed fragments without any clashes between them (3 Å distance between closest atoms). The next step involves linking pairs of fragments using the Difflinker model (1000 linkers per pair) [32]. Through a filtering step that assesses whether generated molecules contain the linked fragments, we define complete molecules, resulting in a library of potentially existing binding molecules.

The subsequent stage is to compare the generated molecules with a database. For this purpose, we selected the Enamine REAL database, one of the largest databases of synthetically accessible small molecules, comprising more than 48 billion compounds [33]. The Enamine software SpaceLight provided an ultra-fast comparison algorithm, using the Tanimoto coefficient on Extended-connectivity fingerprints (ECFPs) fingerprints as the similarity metric [68].

### Experimental validation of designed compounds

To evaluate the capability of our surface fingerprint-guided fragment placement pipeline, we tested designed compounds targeting SARS-CoV-2 Main Protease. Three fully de novo designed molecules using FragmentScope were structurally analysed and experimentally tested (Figure S7A). All three compounds were distant from any known small molecules bound to the target pocket on the protein; all-to-all tanimoto similarity values are presented on the histogram Figure S7A. The placement of the designed compounds was also aligned with the pose of one of the known inhibitors MG-131 PDB ID: 7Z0P bound to the S2 pocket. Even though the interaction profile of the designed compounds didn’t appear to be rather rich, some of the reference interactions were retrieved, such as hydrogen bond with HIS164 for Z559 compound. Additionally we docked the compounds to the targeted pocket, the best retrieved pose was observed for Z559 compound with the RMSD values 3.54 Å and 1.93 Å for the first pose and the closest from first five poses correspondingly (Supplementary Table 8).

GCI experiments were conducted and all compounds showed binding to the target at 2 mM concentration. The positive control measurements have also been done. As a control compound nirmatrelvir a reversible covalent inhibitor was chosen. Due to its covalent behaviour of binding usually instead of K_D_, the constant of inhibition K_i_ is reported in the literature with the value of 3.1 nM [41]. Experimental binding of nirmatrelvir to the SARS Mpro at low nanomolar concentration have shown reliable signal responses and kinetic behavior related to the reversible covalent binding, although the determination of precise K_D_ value is not accurate in this case.

Specificity of the compounds was assessed against KRAS and PIN1 proteins at both concentrations. Z104 showed specificity at 2 mM. NMR experiments of the Z104 compound have shown binding as well at 500 µM to the SARS-CoV-2 Main Protease target. Z106 compound showed binding to the target and one of the off-targets PIN1, however no binding was detected at 500 µM concentration. Z559 compound binding was observed to the target at both concentrations, however off-target binding was also detected to the KRAS and PIN1 at 2 mM concentration, but binding at 500 μM became more specific, so was not detected any binding to PIN1 off-target at 500 μM.

Our second target was phosphoglycerate kinase 1 (PGK1), an essential metabolic enzyme and a well-known target in cancer metabolism [69]. Two computationally generated molecules Z118 and Z123 were experimentally validated (Figure S7B). Both compounds were designed with the same pipeline described earlier by using the surface fingerprint similarity to guide fragment placement into the PGK1 binding site. Both compounds were structurally validated through docking (Supplementary Table 8). The poses of the design compounds Z118 and Z123 aligned closely well with key interaction hotspots in the active site. Interaction profiles of both compounds revealed that they formed consistent polar and hydrophobic interactions within the binding pocket (Figure S7B, interaction panel). Docking analysis indicated limited accuracy in reproducing predicted binding mode for the compound Z118 with RMSD values (4.21 Å for top-1/top-5 poses). Docking results for compound Z123 indicated a significant deviation from the predicted binding mode, reflected by higher RMSD values (7.65 Å for top-1, 5.66 Å for top-5 poses). This discrepancy suggests that while the computational placement captured initial surface signals, compound Z123 adopted an alternative binding pose upon docking probably due to more complexity of the compound, highlighting challenges in accurately predicting fragment interactions for certain cases.

Experimental testing did not show measurable binding of Z118 to PGK1 or to any off-targets (KRAS or PIN1) at either 2 mM or 500 µM, suggesting low or absent affinity (Figure S7B, heatmaps). In contrast, Z123 demonstrated binding to PGK1 at both concentrations tested. However, specificity analysis revealed cross-reactivity with KRAS, indicating that Z123 lacks target selectivity at that concentration. All the measurements were done with the reference measurement of the positive control. The K_D_ obtained with GCI of CHR-6494 positive control to PGK1 target was 30 μM, which is in correspondence to the published value of 37 μM [41]. This data underscores the need for further optimization of the compounds to achieve desired affinity and specificity of compounds.

## Data availability

The crystal structure of Pin1 in complex with compound 155180 deposited at the PDB under accession code 9T04. Crystal structures of BrD4 in complex with compound 662 deposited at the PDB under accession code 9T15 and with compound 960 at the PDB under accession code 9T16.

## Code availability

Fragmentscope scripts are available at GitHub: https://github.com/LPDI-EPFL/FragmentScope.git

## Acknowledgements

We acknowledge the European Synchrotron Radiation Facility (ESRF) for provision of synchrotron radiation facilities under proposal number MX2591 and we would like to thank Christoph Mueller-Dieckmann for assistance and support in using beamline ID30B. We acknowledge Amédé Noredine Larabi for setting up crystallisation plates.

## Supplementary information

**Figure S1.**
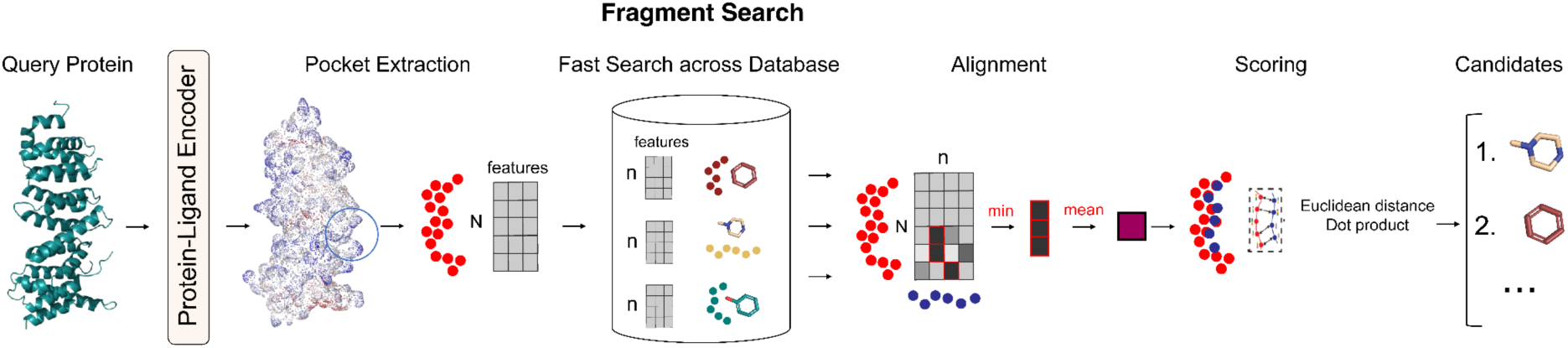
FragmentScope. Fragment search algorithm that consists of three consecutive steps: ultra-fast search, patch alignment and scoring.

**Figure S2.**
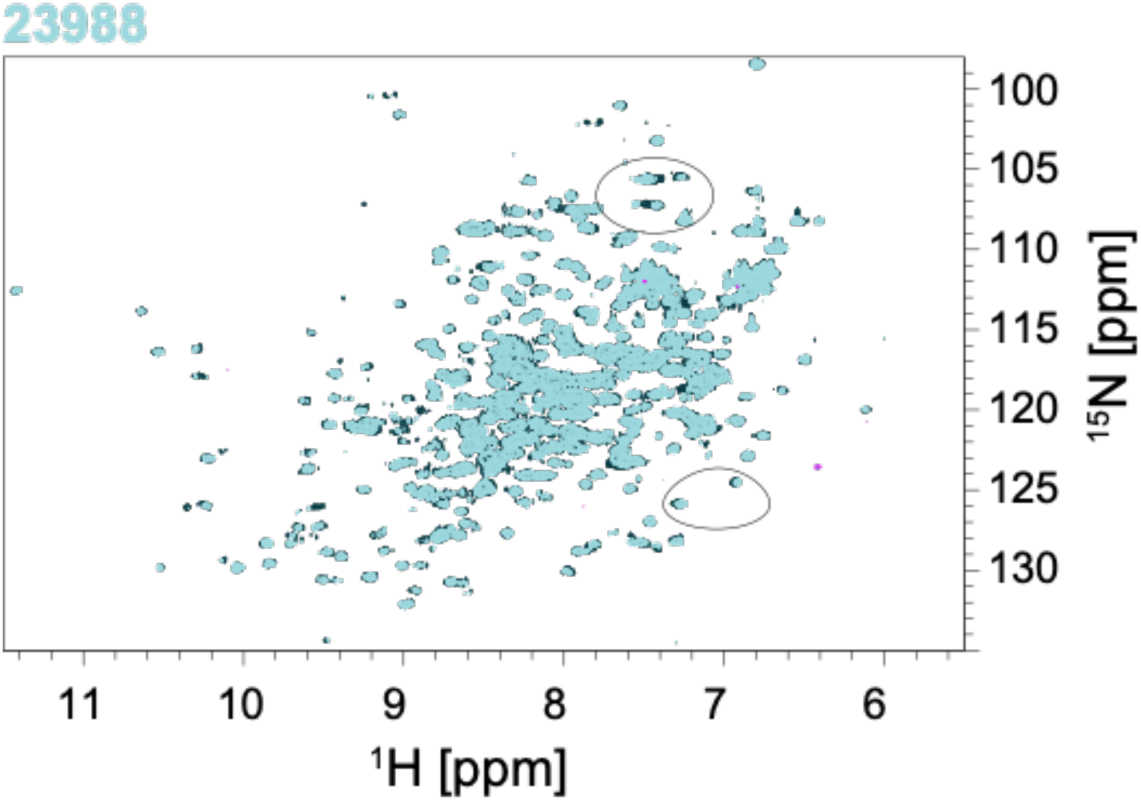
[15N,1H] -HSQC overlay of 100 µM apo-PGK1 G185A (dark blue) bound to 0.5 mM of the 23988 ligand (light blue) - noticeable shifts in the peak positions are visible and circled.

**Figure S3.**
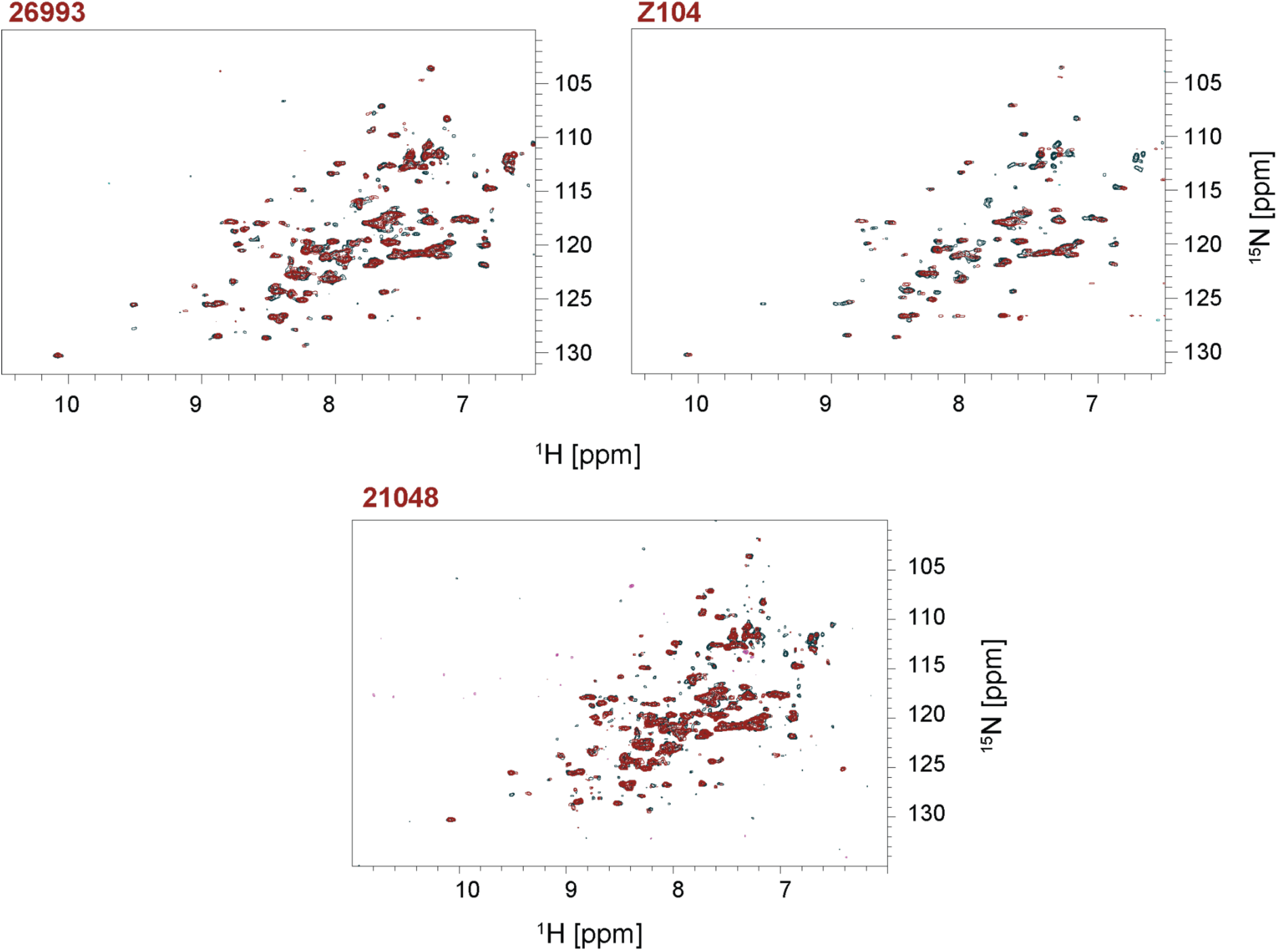
[15N,1H] -HSQC overlay of 50 µM apo-MPro (dark blue) bound to 0.5 mM of the fragments 21048 (red), 26993 (red), ligand Z104 (red) - noticeable shifts in the peak positions are visible throughout the spectrum.

**Figure S4.**
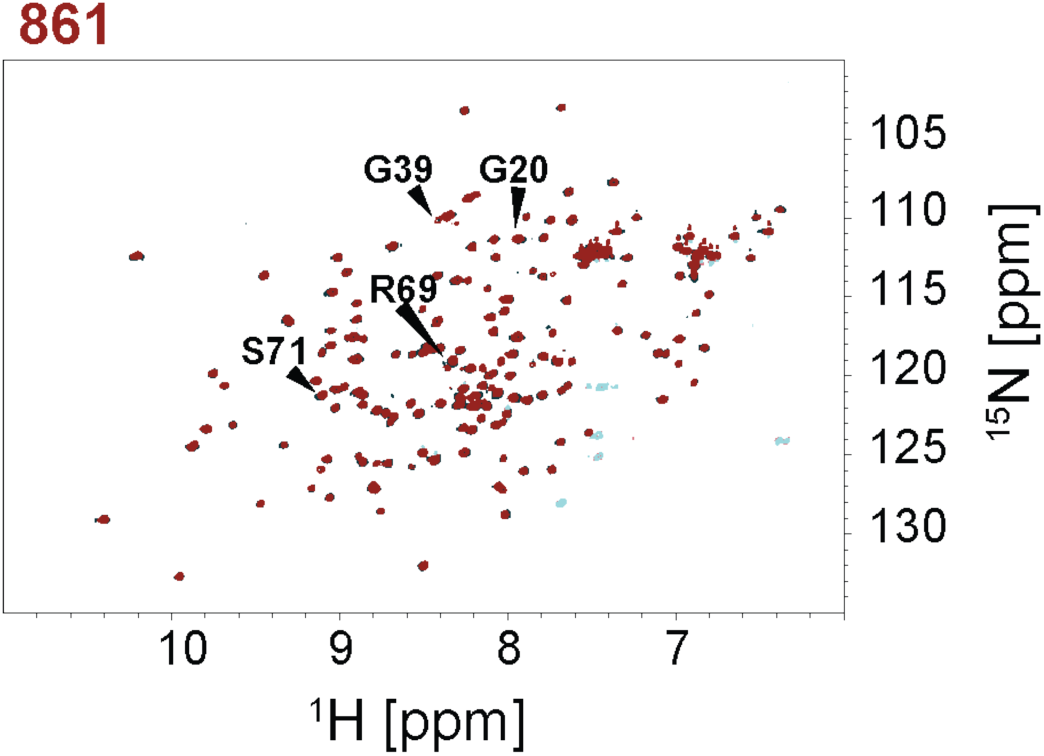
[15N,1H] -HSQC overlay of 200 µM apo-PIN1 (dark blue) bound to 0.5 mM of the fragment 861 (red), binding of the 861 fragment was confirmed, noticeable shifts in the peak positions are marked.

**Figure S5.**
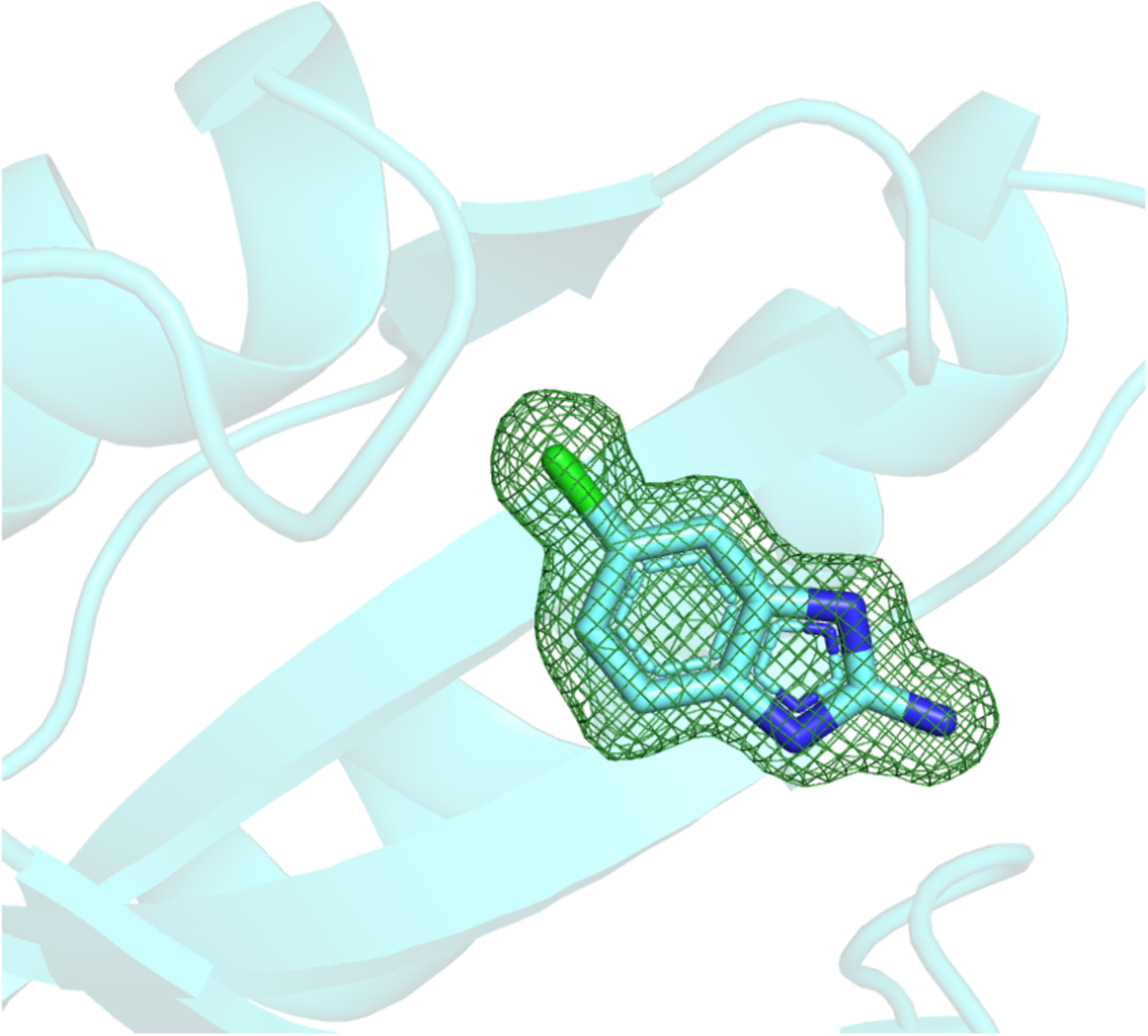
Omit map: PIN1 in a complex with 155 compound.

**Figure S6.**
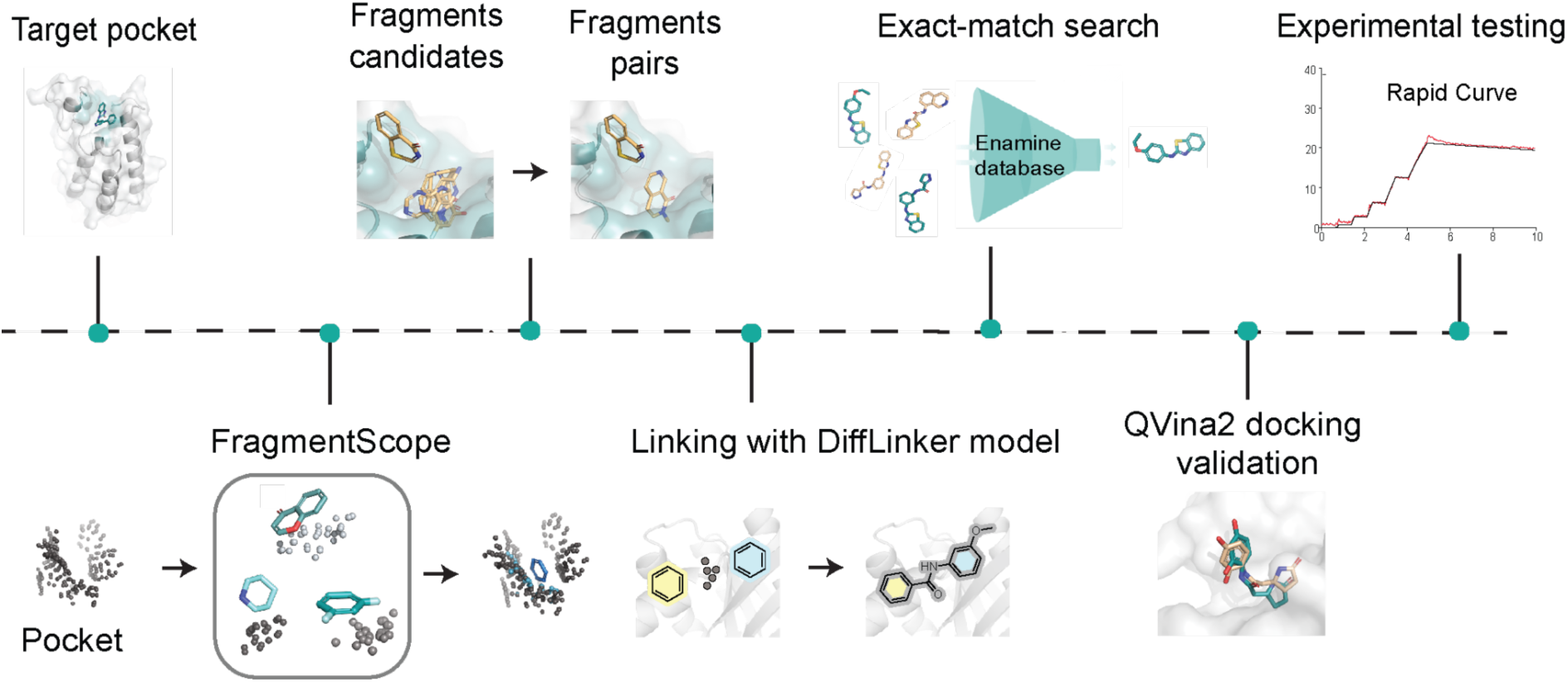
Overview of the fragment-based small molecule pipeline. Multiple steps are presented sequentially, starting from fragments placement with FragmentScope, following with generation of pairs of fragments. Linking step with DiffLinker, comparison of the generated molecules to Enamine Database [31], docking of the exact-match molecules and experimental validation.

**Figure S7.**
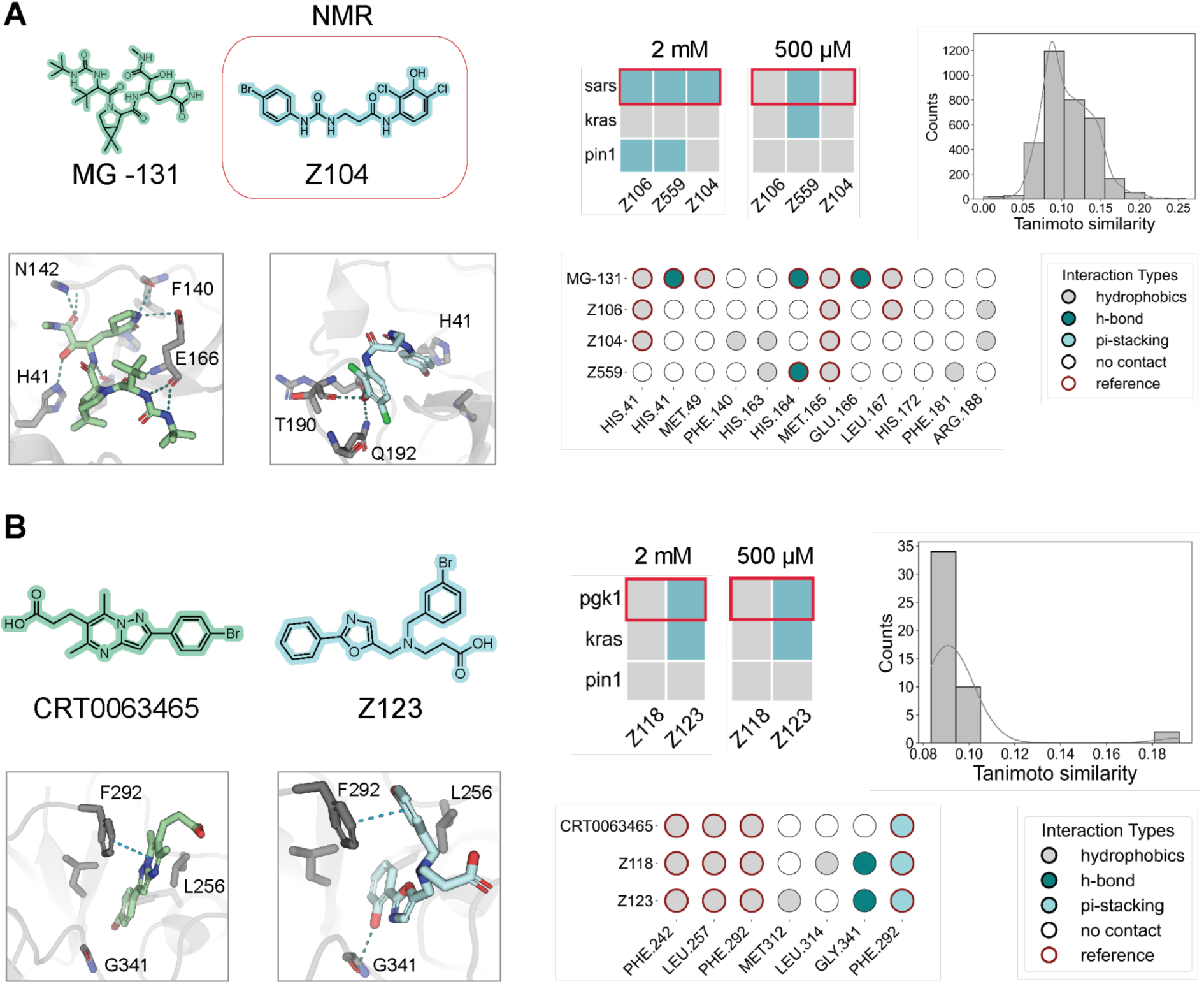
Experimental validation and structural analysis of fragment-based designed small molecules across three protein targets: SARS Mpro and PGK1. **A)** Reference compound MG-131 and designed compound Z104, confirmed by NMR. Shown are chemical structures and 3D placement within the binding pocket. Heatmaps indicate binding specificity (blue – binding, grey – no binding) at 2 mM and 500 µM across two off-targets (KRAS and PIN1). Tanimoto similarity histogram highlights chemical novelty compared to all known small molecules. Interaction maps summarize contact profiles of predicted protein-ligand complexes in comparison to the MG-131. **B)** Known ligand CRT0063465 and designed compound Z123 in PGK1. Chemical structures, crystal structure placement, and predicted binding poses are shown. Binding profiles to PGK1 and off-targets, interaction maps, and novelty assessment (Tanimoto similarity histogram) are presented as in panel A.

**Figure S8.**
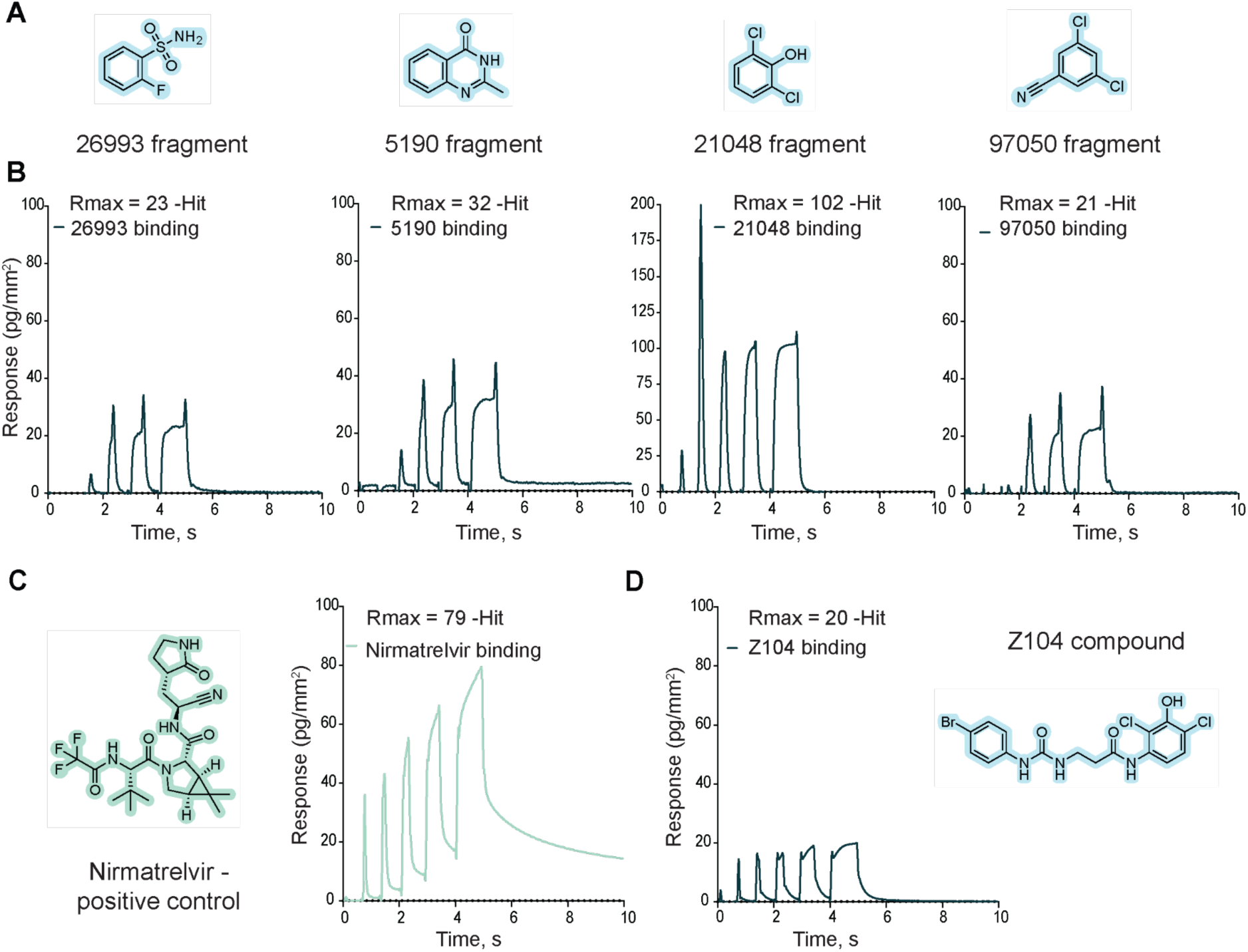
SARS-CoV-2 Main Protease case kinetic data: fragments and ligands. **A)** Structures of SARS fragments. **B)** Examples of binding kinetic sensorgrams (rapid kinetic) for hit fragments. **C)** SARS positive control: structure of Nirmatrelvir and kinetic data. **D)** Kinetic sensorgram and structure of SARS hit Z104 compound.

**Figure S9.**
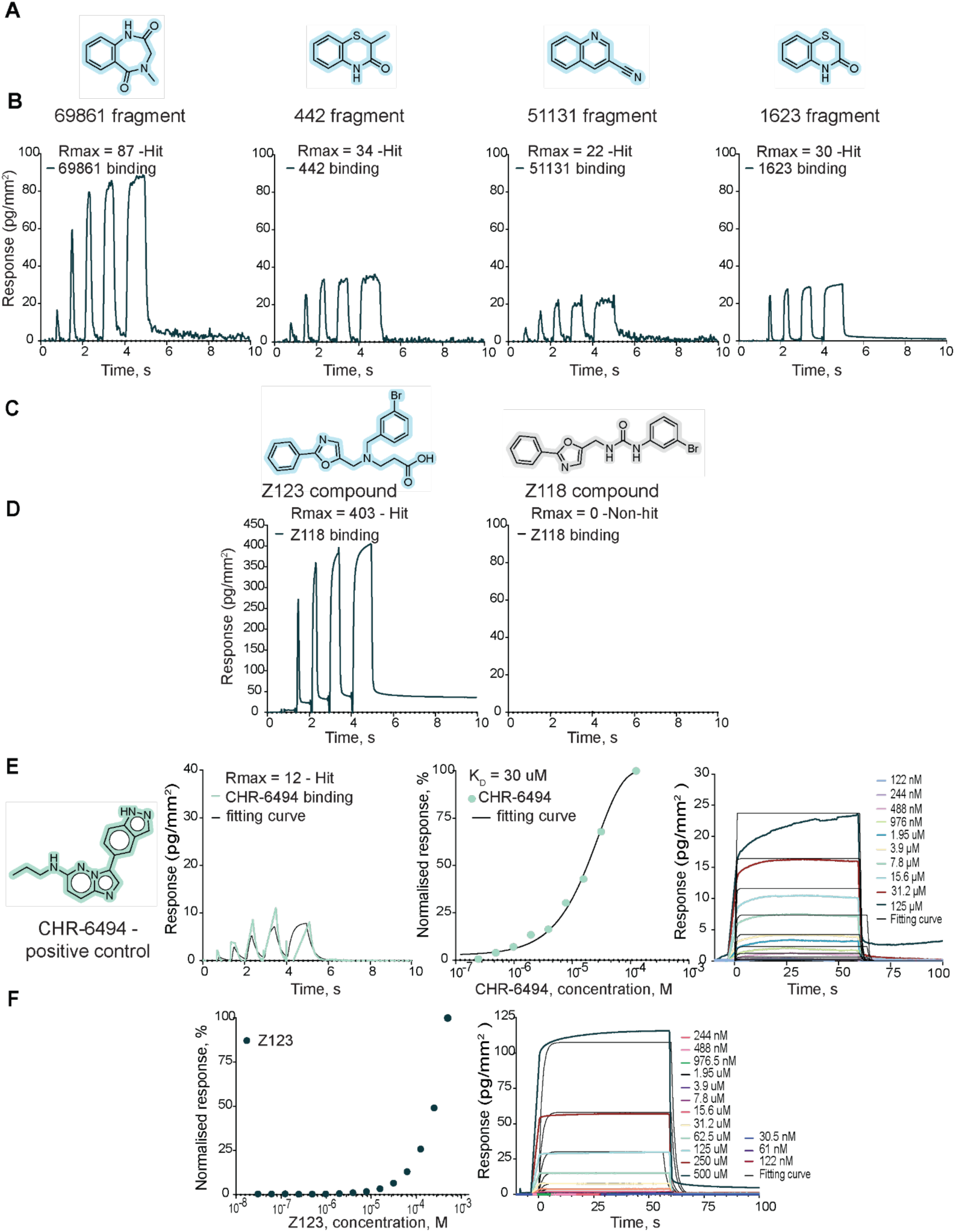
PGK1 case kinetic data: fragments and ligands. **A)** Structures of PGK1 fragments. **B)** Examples of binding kinetic sensorgrams (rapid kinetic) for hit fragments. **C)** Structures of PGK1 compounds. **D)** Kinetic sensorgrams of PGK1 compounds. **E)** Positive control CHR-6494 for PGK1: structure and kinetic data, K_D_ = 30 uM. **F)** Multicycle kinetic data and K_D_ determination for PGK1 compound Z123: K_D_ - mM* range binder (K_D_ - undefined).

**Figure S10.**
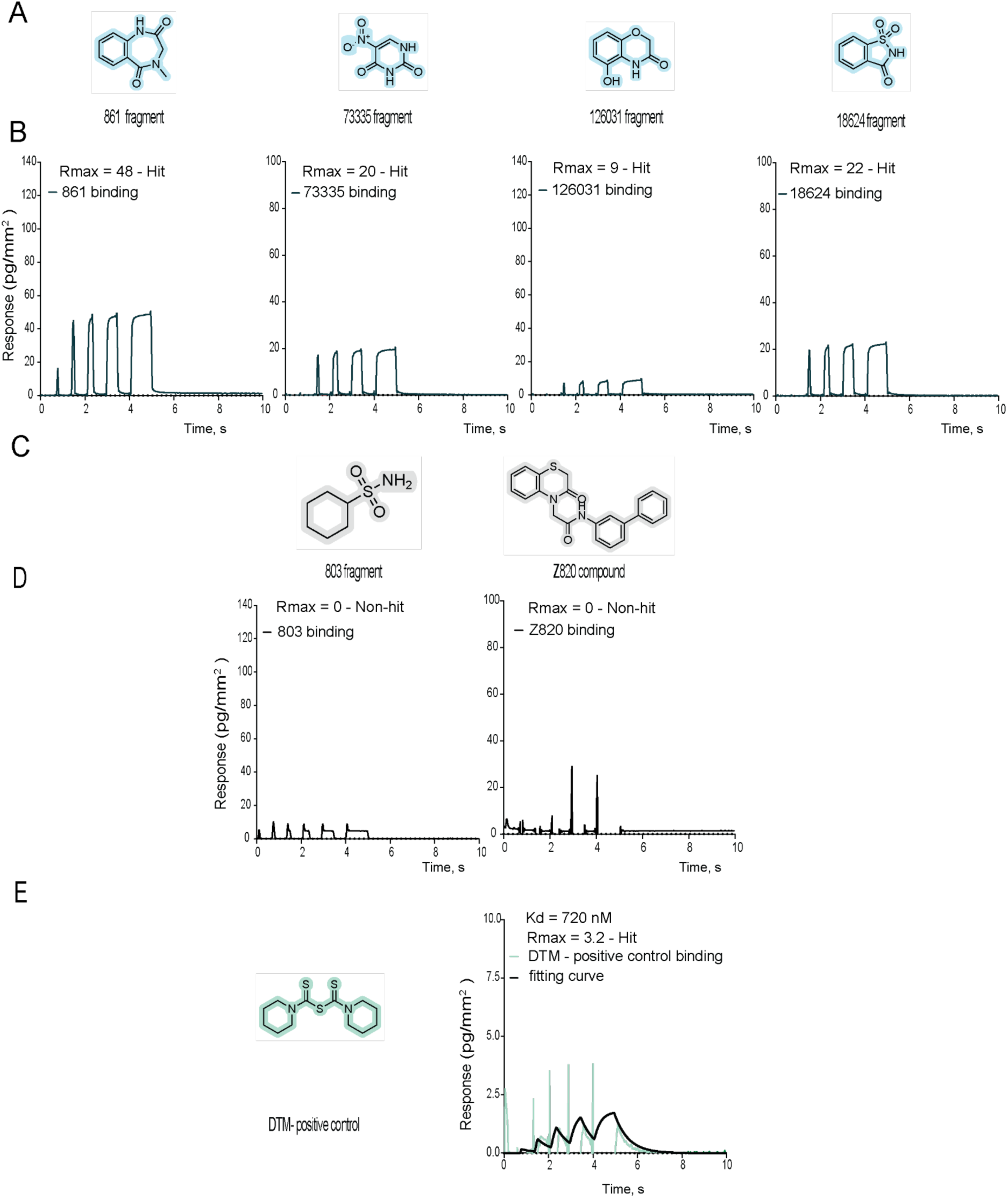
PIN1 case kinetic data: fragments and ligands. **A)** Structures of PIN1 fragments. **B)** Examples of binding kinetic sensorgrams (rapid kinetic) for hit fragments. **C)** Structures of PIN1 non-hit fragments/compounds. **D)** Kinetic sensorgrams of PIN1 non-hit fragments/compounds. E) Positive control DTM for PIN1: structure and kinetic data, K_D_ = 720 nM.

**Figure S11.**
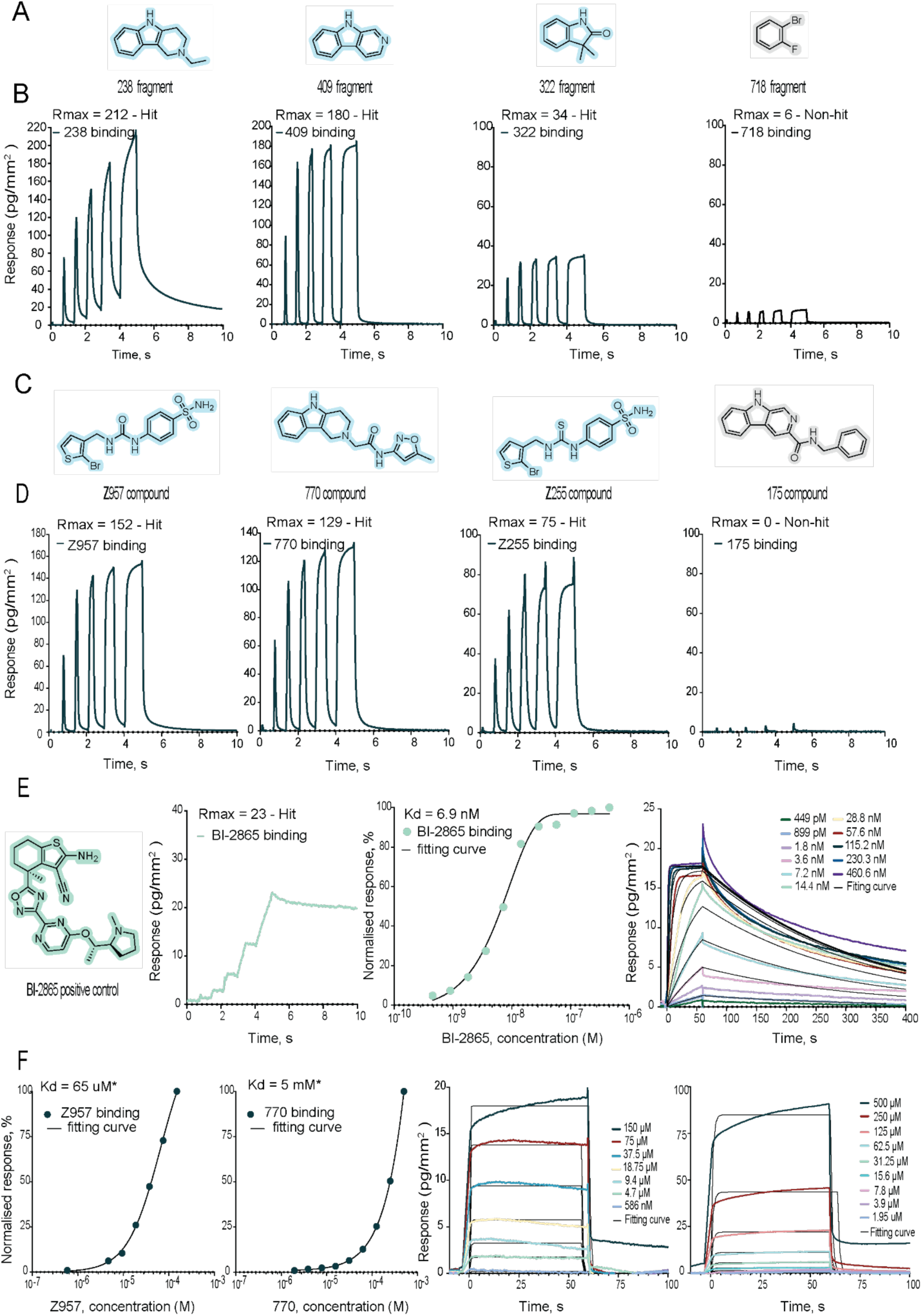
KRAS case kinetic data: fragments and ligands. **A)** Structures of KRAS fragments. **B)** Examples of binding kinetic sensorgrams (rapid kinetic) for hit and non-hit fragments. **C)** Structures of KRAS compounds. **D)** Kinetic sensorgrams of KRAS compounds. **E)** Positive control BI-2865 data for KRAS: structure and kinetic data, K_D_ = 6.9 nM. **F)** Multicycle kinetic data and K_D_ determination for KRAS compounds: K_D_(Z957)=65 uM*, K_D_(770)=5 mM*.

**Figure S12.**
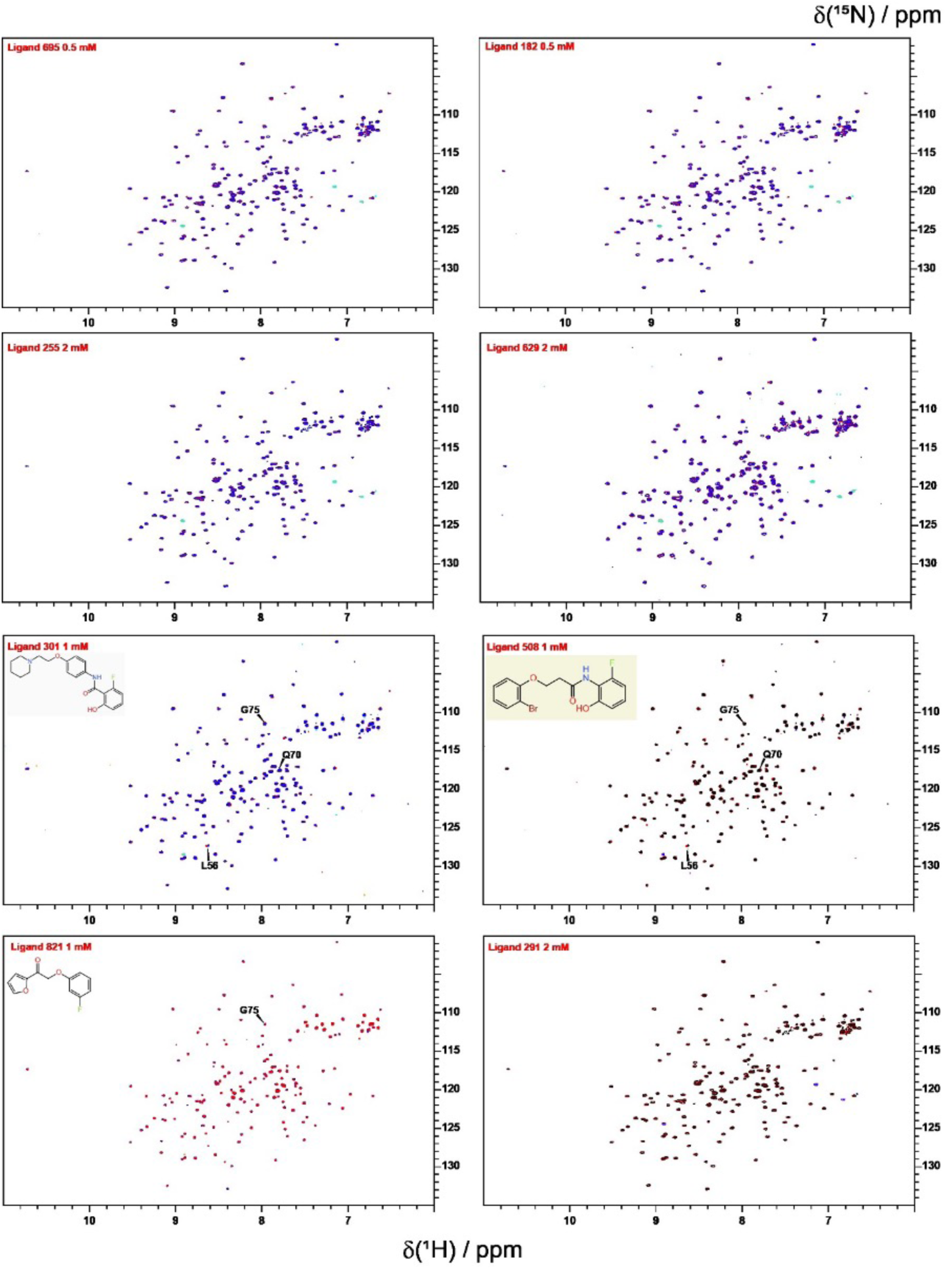
[15N,1H] -HSQC overlay of apo-KRAS G12V 100 µM (blue or black) with ligand (red) - for table 2 KRAS compounds (known fragment, novel linker).

**Figure S13.**
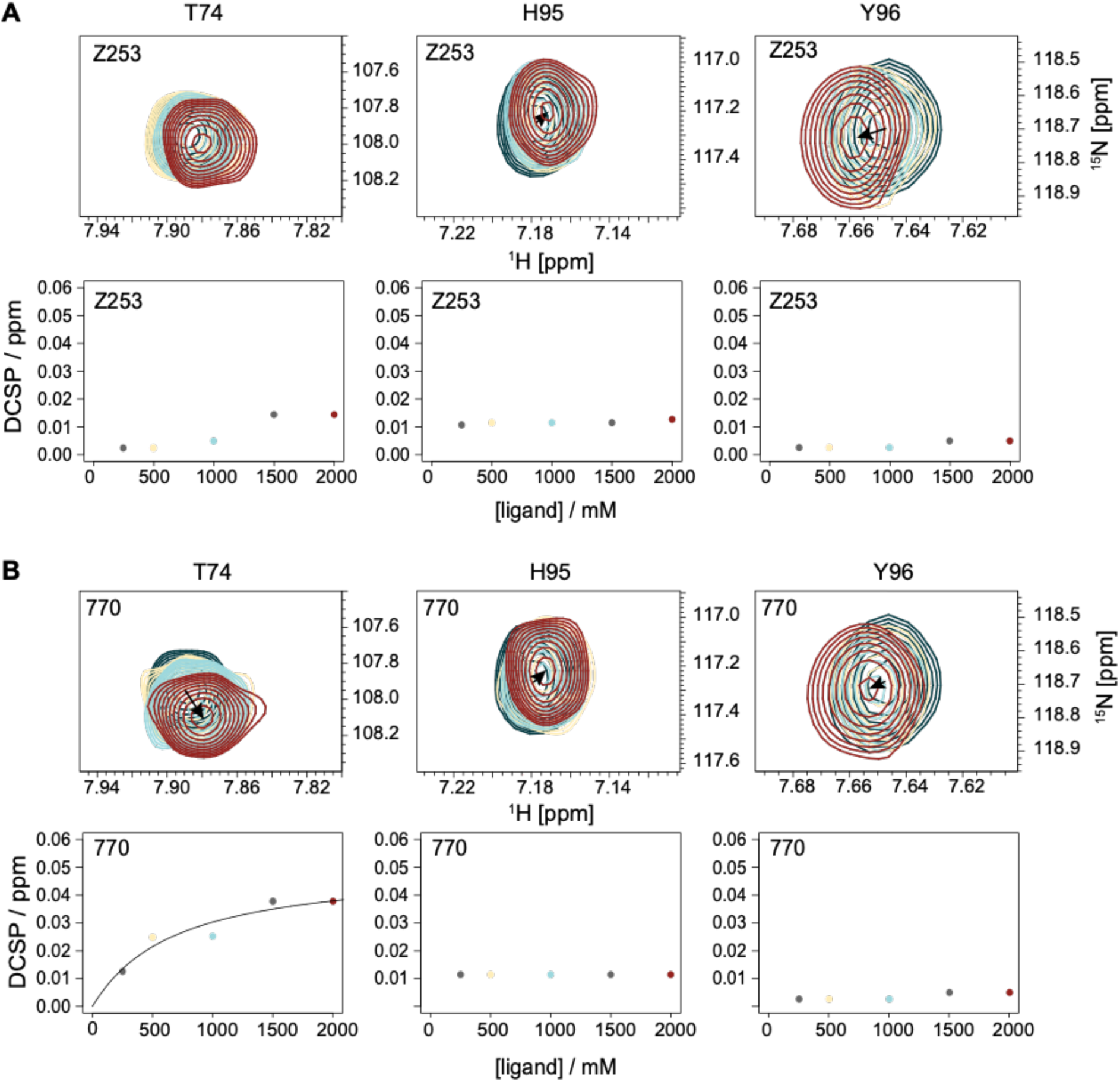
[^15^N,^1^H]-HSQC titrations of compound Z253 confirm binding to KRAS WT. **A)** The chemical shifts of the 3 residues T74, H95 and Y96 are shown with varying amounts of ligand Z253. **B)** The K_D_ titration curves, the color coding for the various concentrations of the ligand can be read from the K_D_ titration curves, which were done with 80 µM KRAS. Due to the weak binding the K_D_ value was not defined.

**Figure S14.**
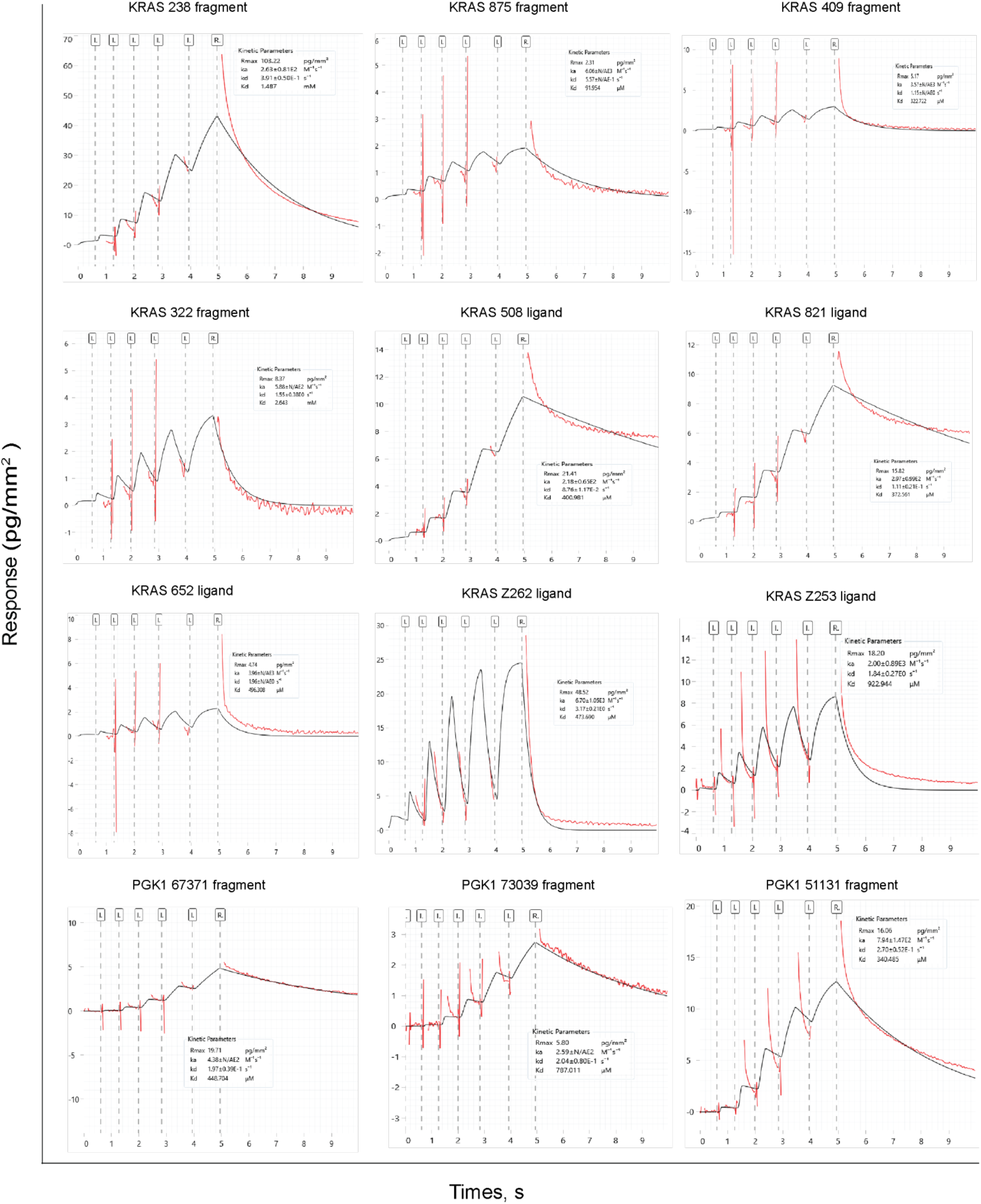
Representative rapid kinetic sensorgrams from the kinetic screening of library (fragments and ligands) per protein target. Corresponding values of K_D_ 1 : 1 interaction kinetic model was globally (black traces) fitted to the experimental sensorgrams (red traces) providing the information about the range of affinities.

**Figure S15.**
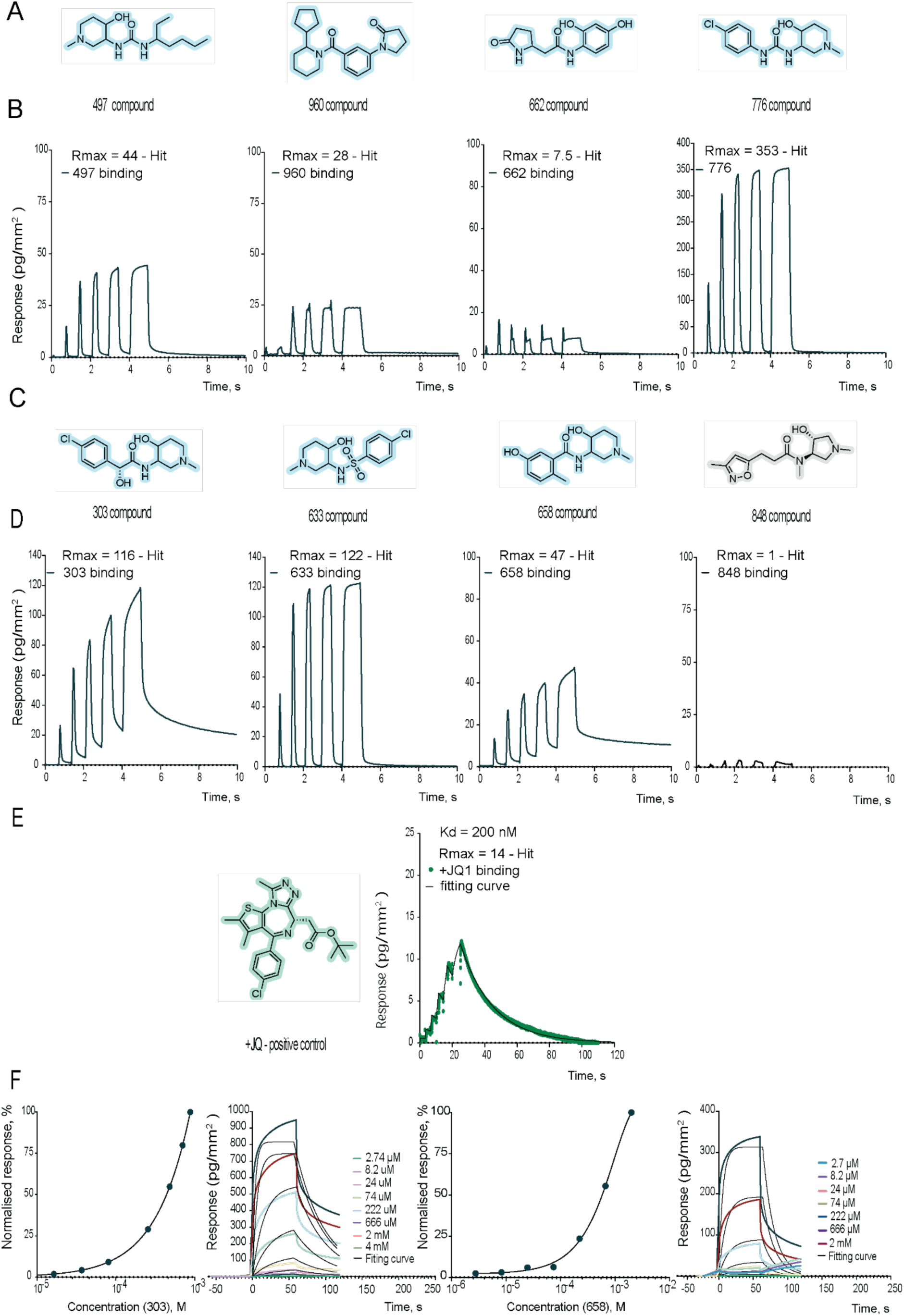
BrD4 case kinetic data: ligands. **A)** Structures of BrD4 ligands (first round of designs). **B)** Examples of binding kinetic sensorgrams (rapid kinetic) for hit ligands (first round of designs). **C)** Structures of BrD4 ligands (second round of designs). **D)** Examples of binding kinetic sensorgrams for hit and non-hit ligands (second round of designs). **E)** Positive control +JQ1 for BrD4: structure and kinetic data, K_D_ = 200 nM. **F)** Multicycle kinetic data and K_D_ determination for BrD4 ligands: 303 and 658 - K_D_ in mM* range (undefined).

**Figure S16.**
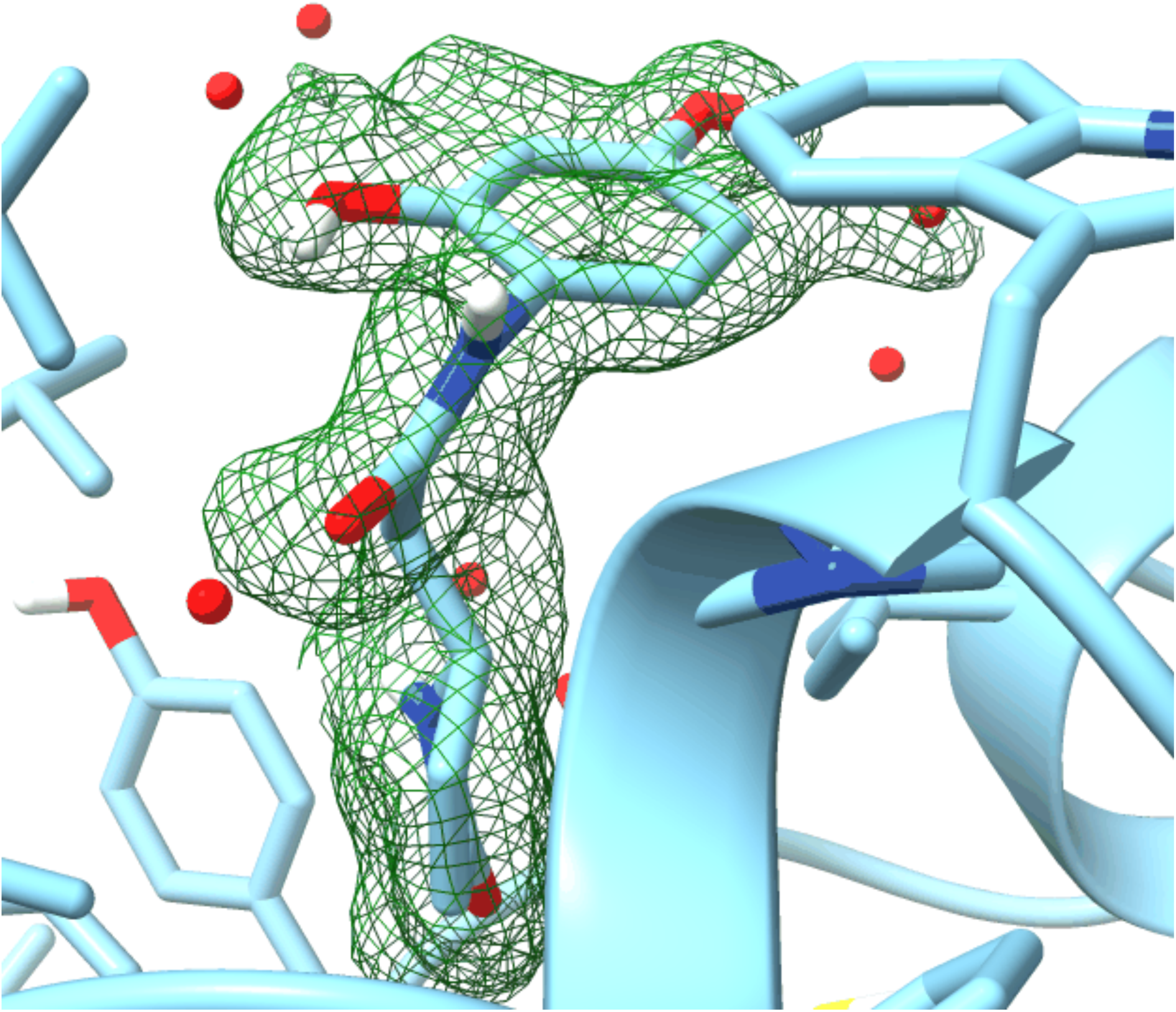
Omit map: BrD4 in a complex with 662 compound.

**Figure S17.**
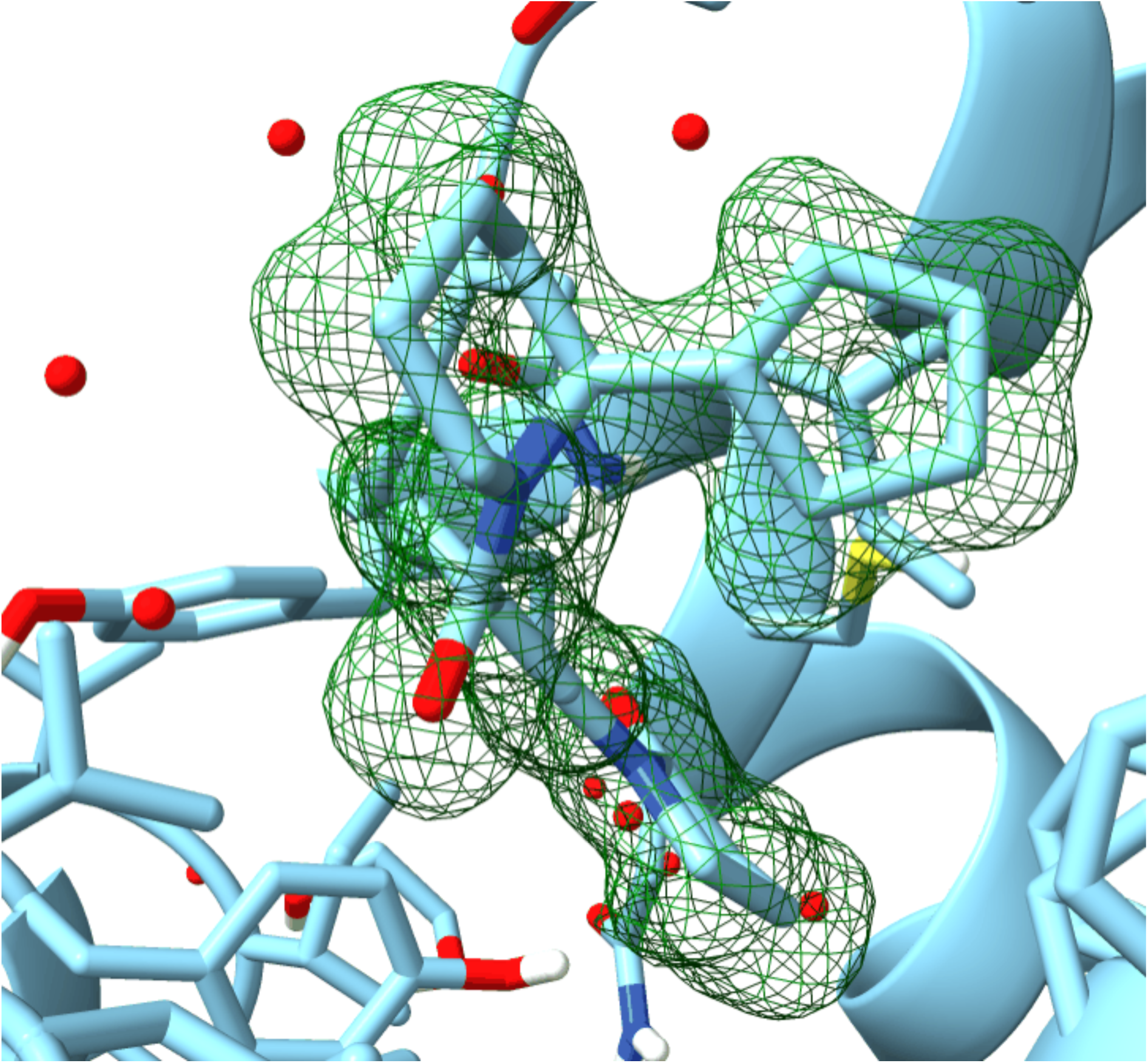
Omit map: BrD4 in a complex with 960 compound.

