## Supplementary Information for "FragmentScope - exploring the fragment space with learned surface representations"

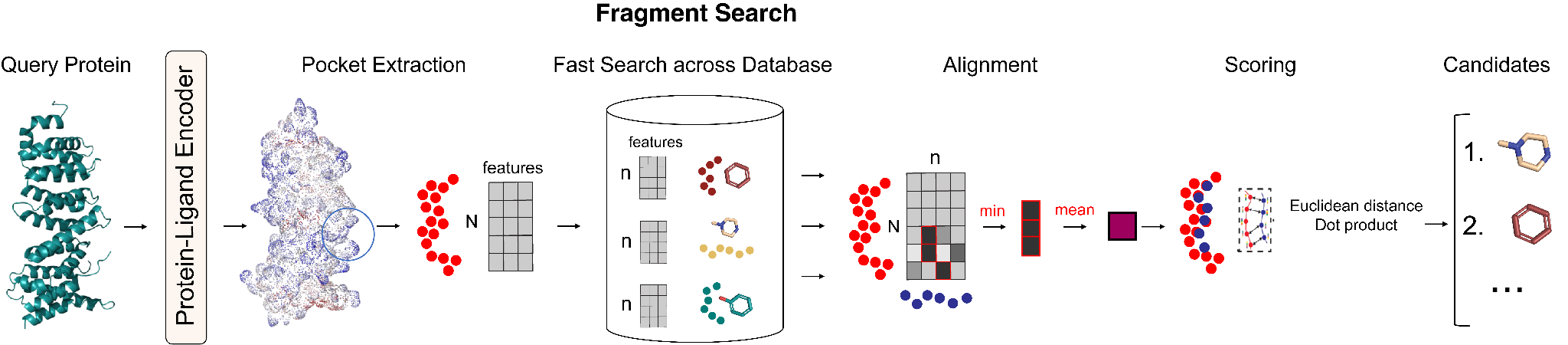

**Figure S1. FragmentScope.** Fragment search algorithm that consists of three consecutive steps: ultra-fast search, patch alignment and scoring.

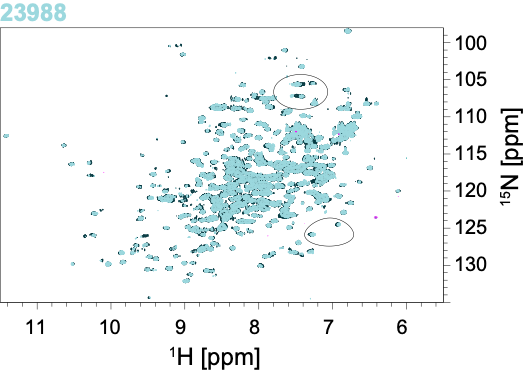

**Figure S2**. [15N,1H] -HSQC overlay of 100 µM apo-PGK1 G185A (dark blue) bound to 0.5 mM of the 23988 ligand (light blue) - noticeable shifts in the peak positions are visible and circled.

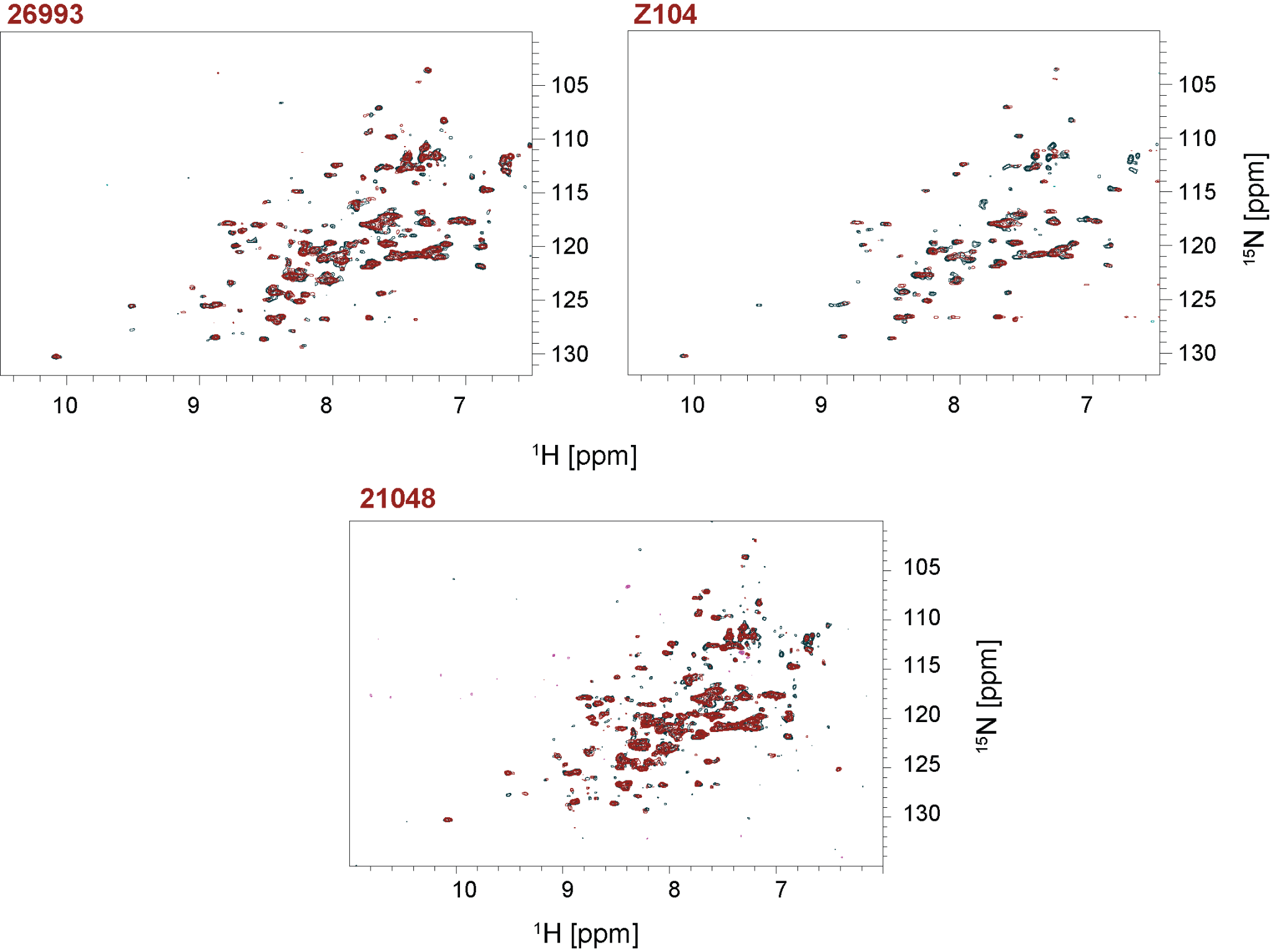

**Figure S3.** [15N,1H] -HSQC overlay of 50 µM apo-MPro (dark blue) bound to 0.5 mM of the fragments 21048 (red), 26993 (red), ligand Z104 (red) - noticeable shifts in the peak positions are visible throughout the spectrum.

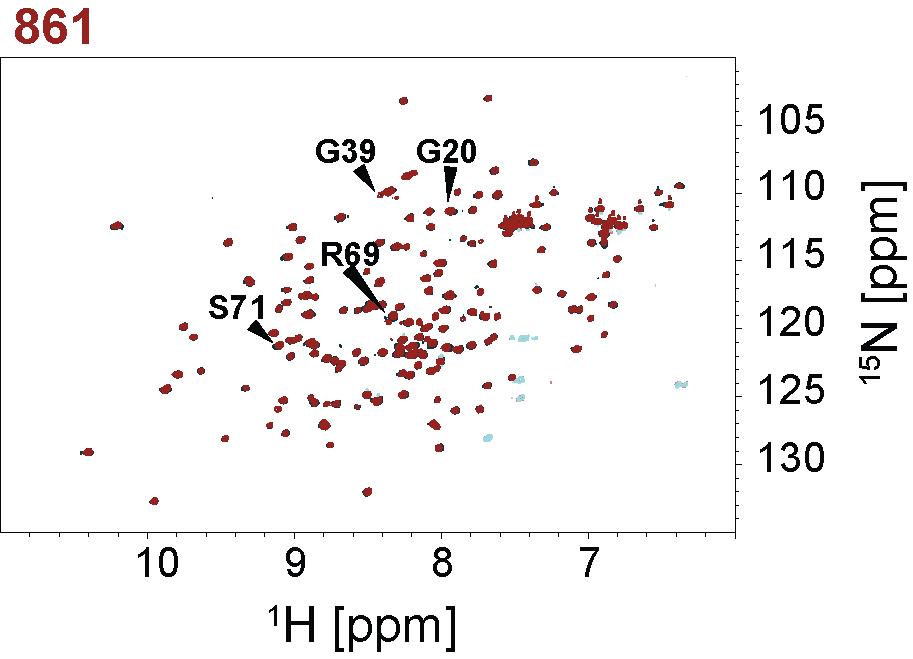

**Figure S4.** [15N,1H] -HSQC overlay of 200 µM apo-PIN1 (dark blue) bound to 0.5 mM of the fragment 861 (red), binding of the 861 fragment was confirmed, noticeable shifts in the peak positions are marked.

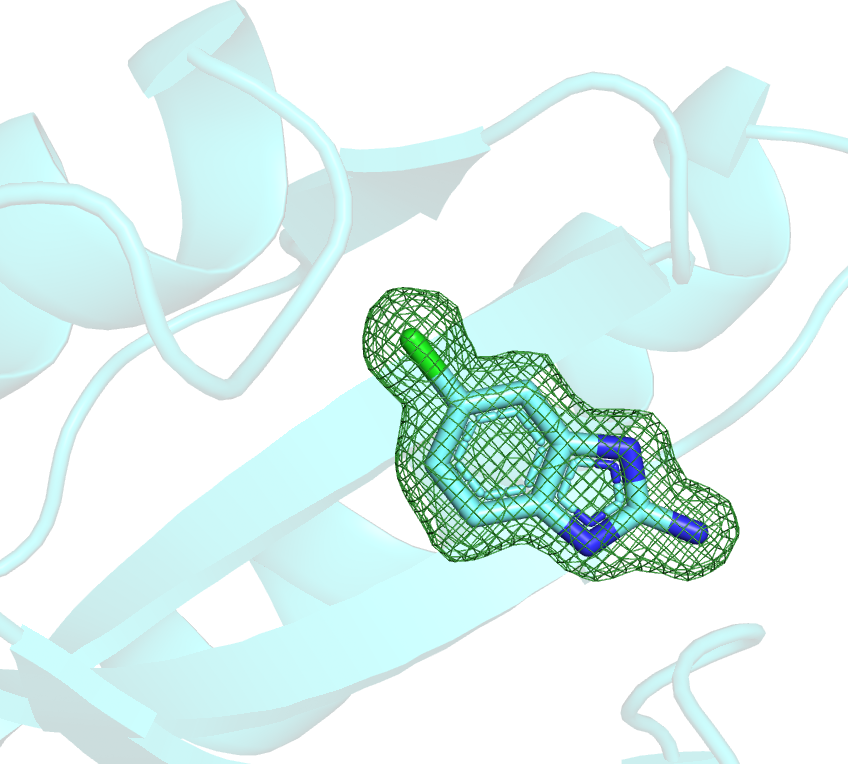

**Figure S5.** Omit map: PIN1 in a complex with 155 compound.

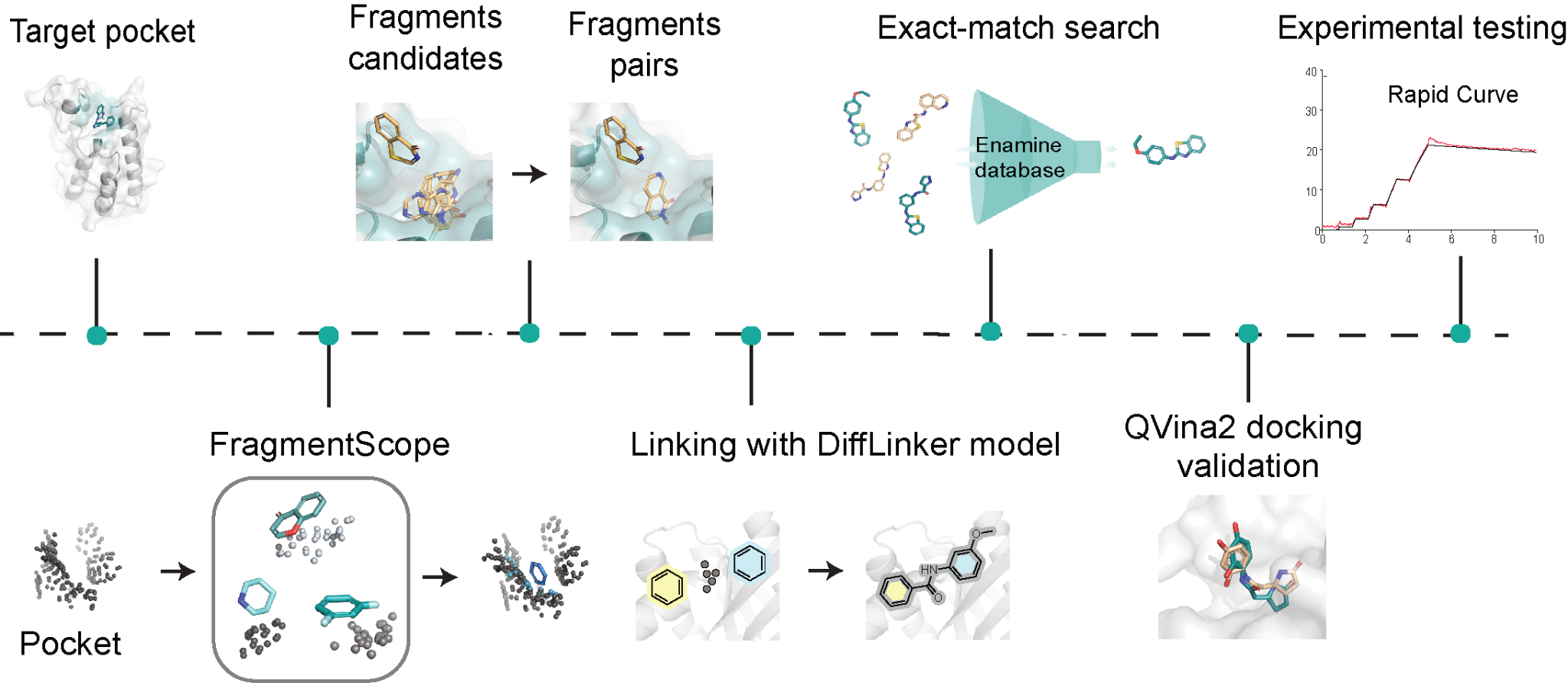

**Figure S6. Overview of the fragment-based small molecule pipeline**. Multiple steps are presented sequentially, starting from fragments placement with FragmentScope, following with generation of pairs of fragments. Linking step with DiffLinker, comparison of the generated molecules to Enamine Database [31], docking of the exact-match molecules and experimental validation.

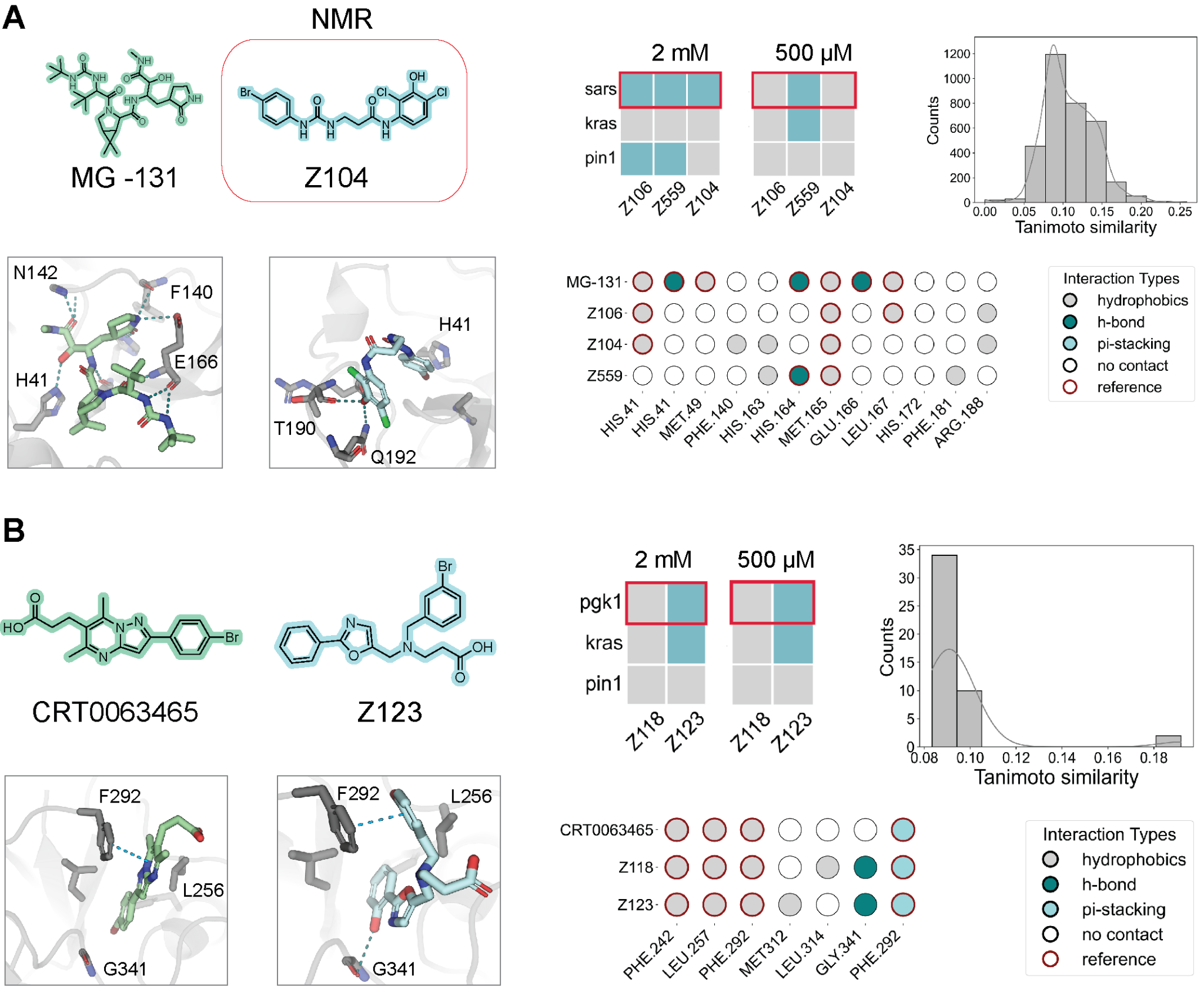

**Figure S7. Experimental validation and structural analysis of fragment-based designed small molecules across three protein targets: SARS Mpro and PGK1. A)** Reference compound MG-131 and designed compound Z104, confirmed by NMR. Shown are chemical structures and 3D placement within the binding pocket. Heatmaps indicate binding specificity (blue – binding, grey – no binding) at 2 mM and 500 µM across two off-targets (KRAS and PIN1). Tanimoto similarity histogram highlights chemical novelty compared to all known small molecules. Interaction maps summarize contact profiles of predicted protein-ligand complexes in comparison to the MG-131. **B)** Known ligand CRT0063465 and designed compound Z123 in PGK1. Chemical structures, crystal structure placement, and predicted binding poses are shown. Binding profiles to PGK1 and off-targets, interaction maps, and novelty assessment (Tanimoto similarity histogram) are presented as in panel A.

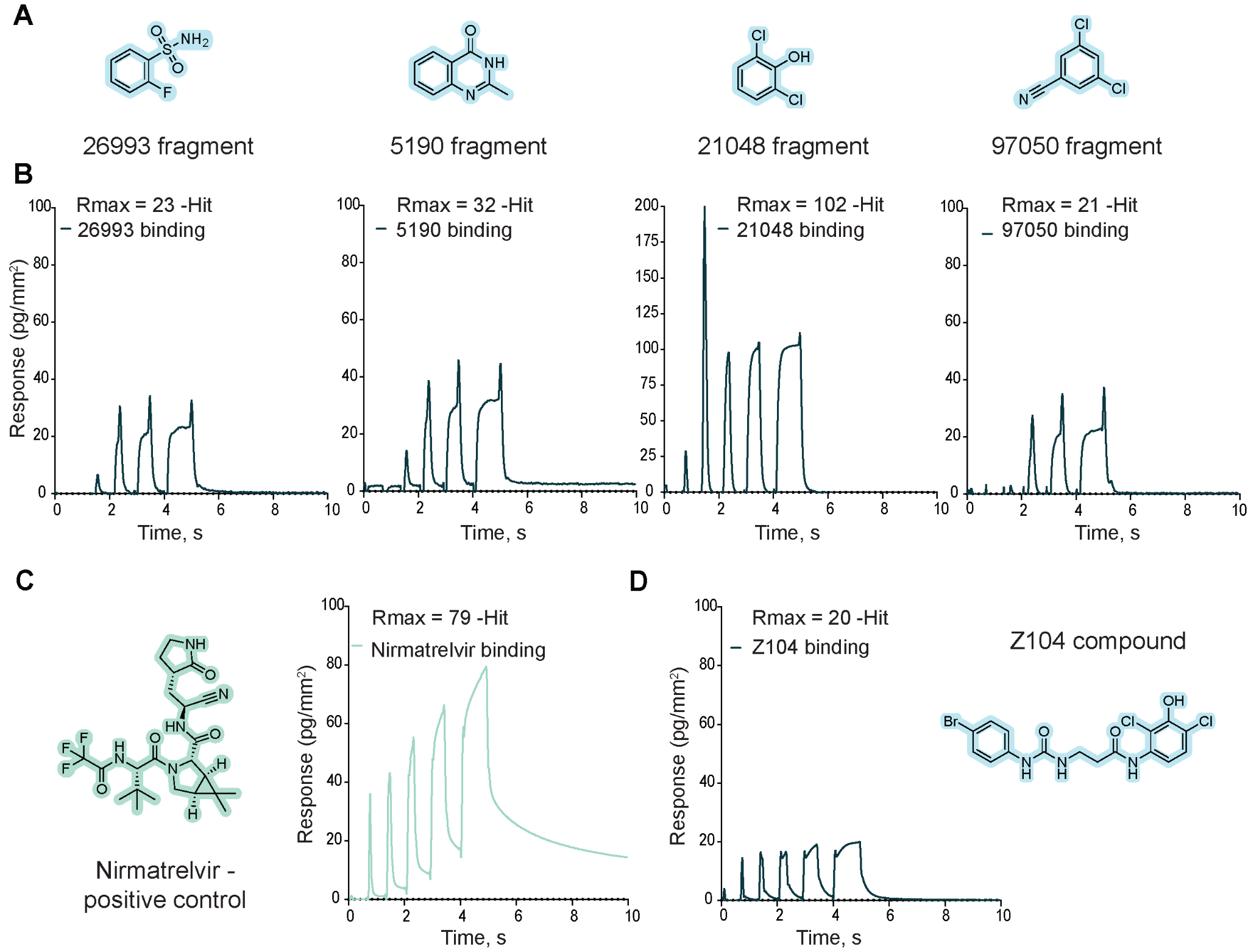

**Figure S8.** **SARS-CoV-2 Main Protease** **case kinetic data: fragments and ligands. A)** Structures of SARS fragments. **B)** Examples of binding kinetic sensorgrams (rapid kinetic) for hit fragments. **C)** SARS positive control: structure of Nirmatrelvir and kinetic data. **D)** Kinetic sensorgram and structure of SARS hit Z104 compound.

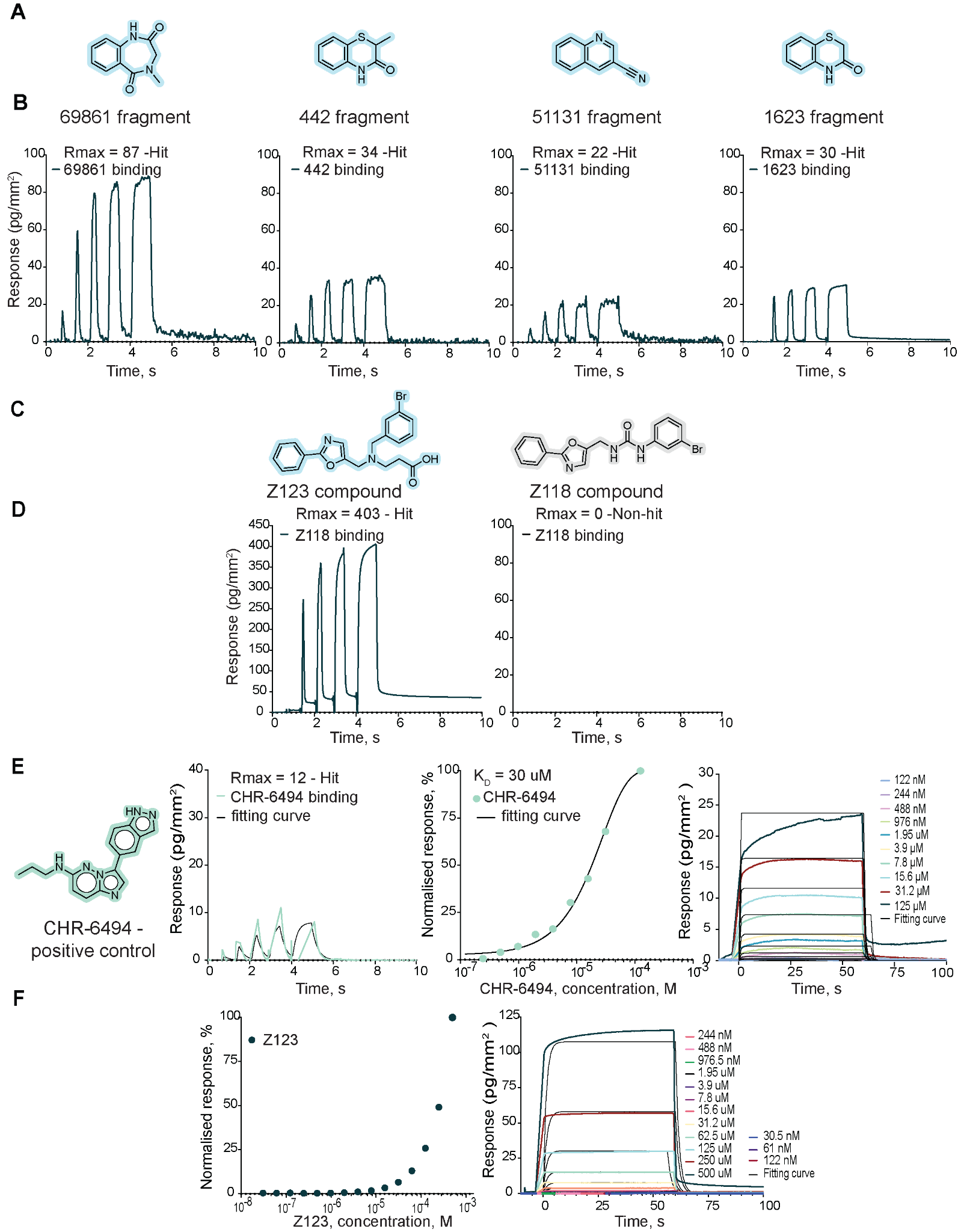

**Figure S9. PGK1 case kinetic data: fragments and ligands. A)** Structures of PGK1 fragments. **B)** Examples of binding kinetic sensorgrams (rapid kinetic) for hit fragments. **C)** Structures of PGK1 compounds. **D)** Kinetic sensorgrams of PGK1 compounds. **E)** Positive control CHR-6494 for PGK1: structure and kinetic data, K_D_ = 30 uM. **F)** Multicycle kinetic data and K_D_ determination for PGK1 compound Z123: K_D_ - mM* range binder (K_D_ - undefined).

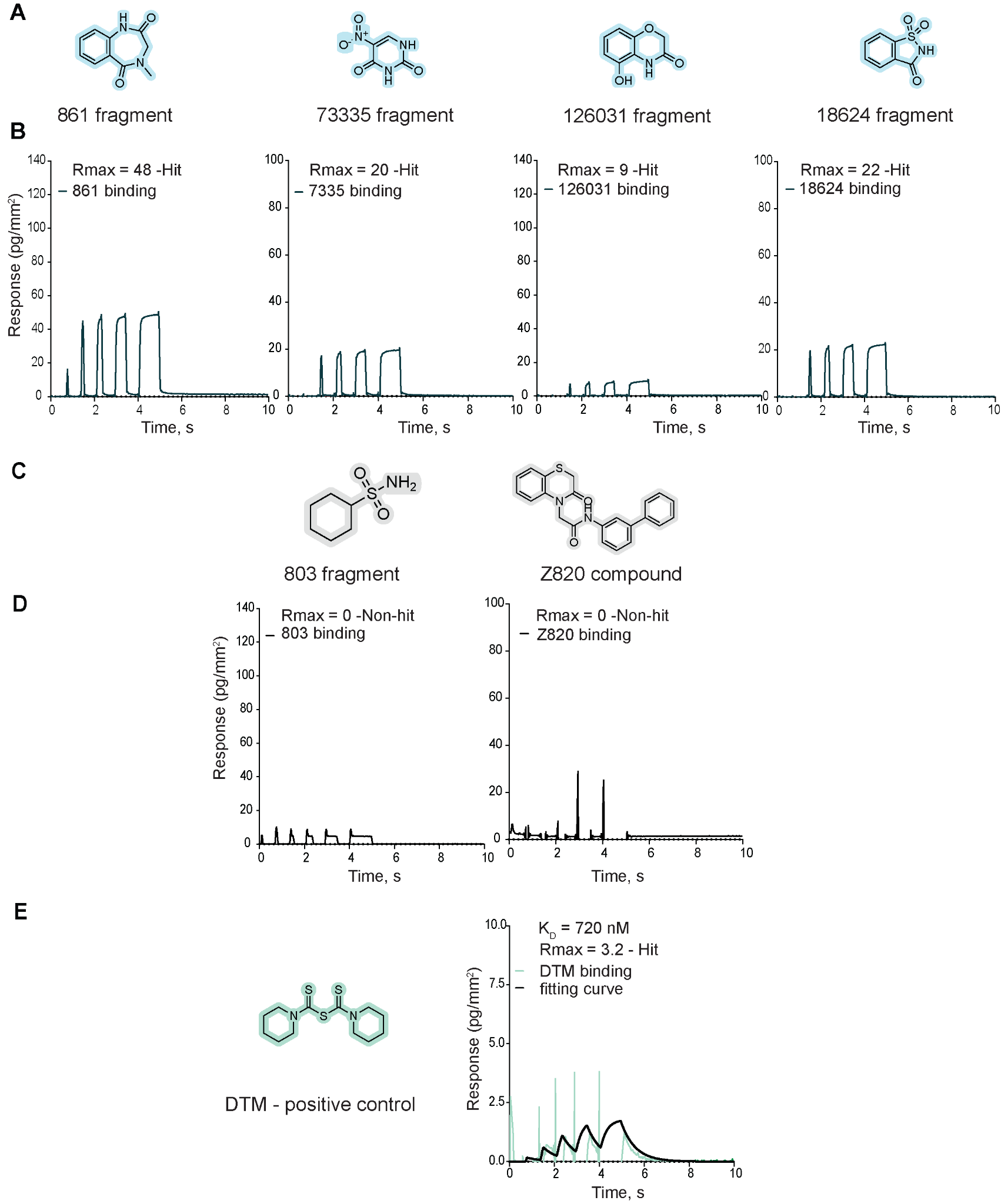

**Figure S10. PIN1 case kinetic data: fragments and ligands. A)** Structures of PIN1 fragments. **B)** Examples of binding kinetic sensorgrams (rapid kinetic) for hit fragments. **C)** Structures of PIN1 non-hit fragments/compounds. **D)** Kinetic sensorgrams of PIN1 non-hit fragments/compounds. E) Positive control DTM for PIN1: structure and kinetic data, K_D_ = 720 nM.

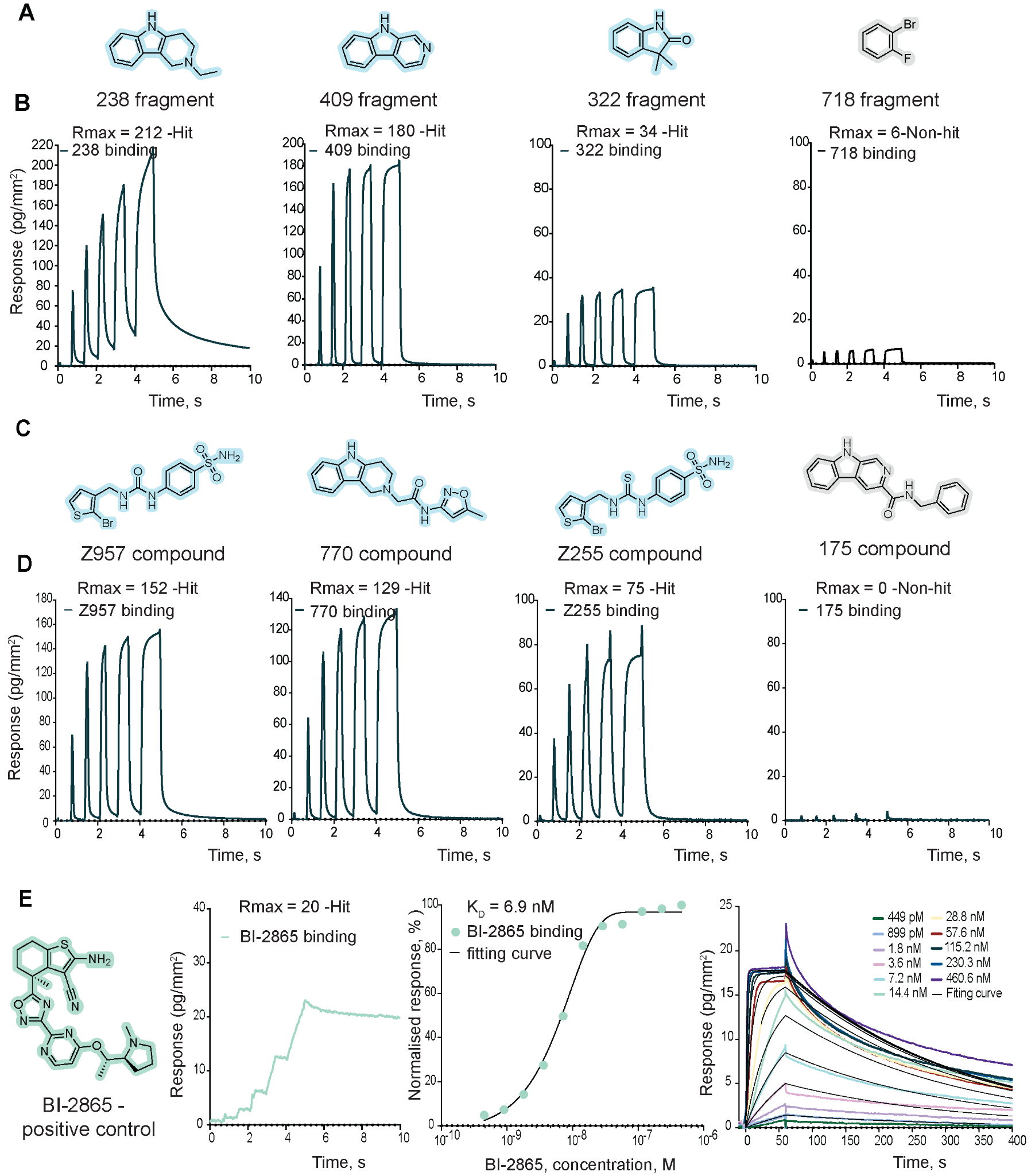

**Figure S11. KRAS case kinetic data: fragments and ligands.** **A)** Structures of KRAS fragments. **B)** Examples of binding kinetic sensorgrams (rapid kinetic) for hit and non-hit fragments. **C)** Structures of KRAS compounds. **D)** Kinetic sensorgrams of KRAS compounds. **E)** Positive control BI-2865 data for KRAS: structure and kinetic data, K_D_ = 6.9 nM.

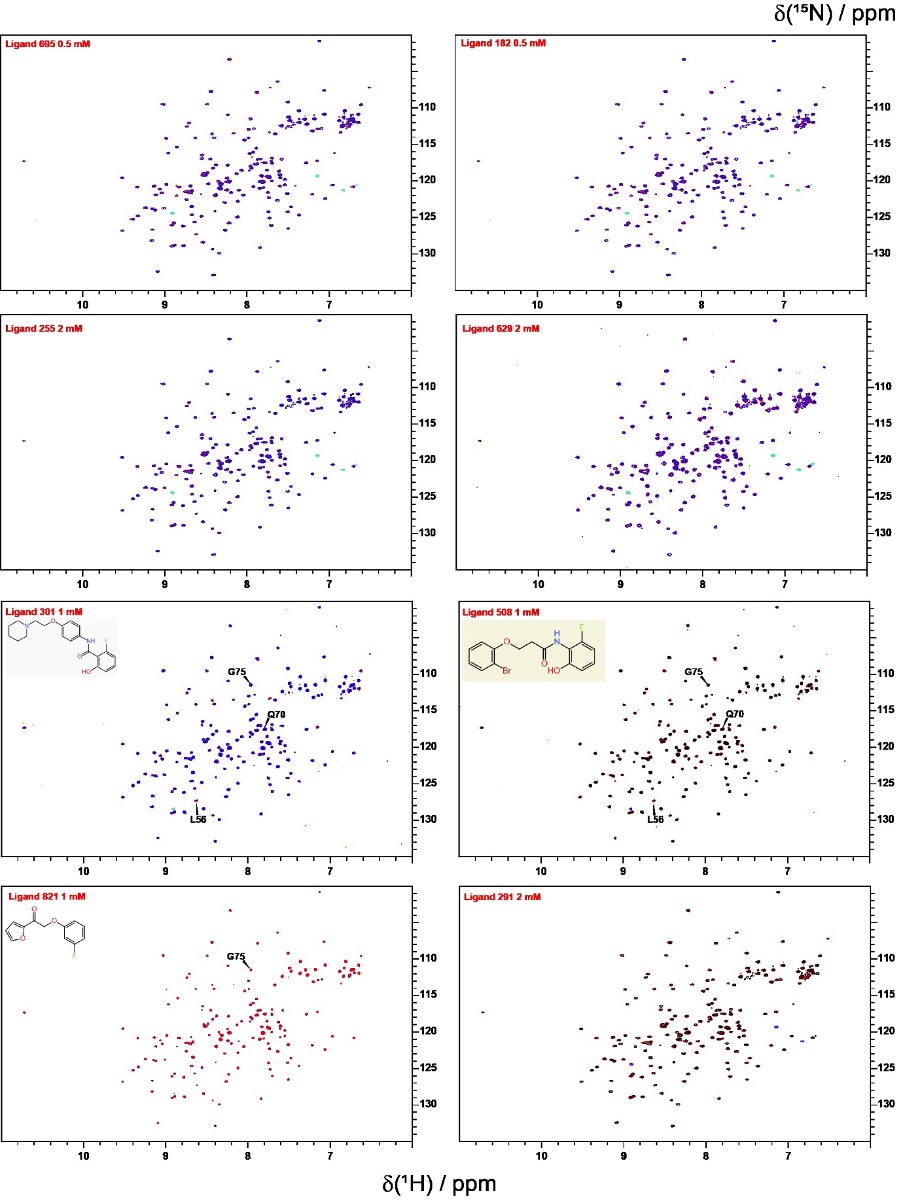

**Figure S12.** [15N,1H] -HSQC overlay of apo-KRAS G12V 100 µM (blue or black) with ligand (red) - for table 2 KRAS compounds (known fragment, novel linker).

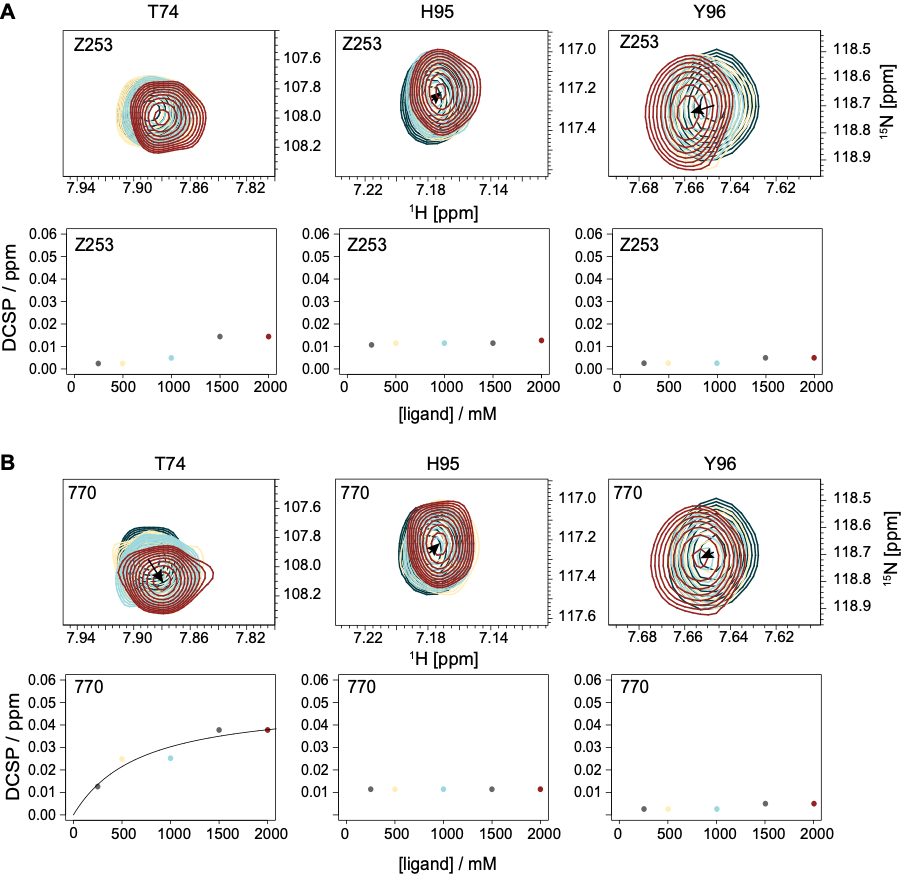

**Figure S13.** **[^15^N,^1^H]-HSQC titrations of compound Z253 confirm binding to KRAS WT. A)** The chemical shifts of the 3 residues T74, H95 and Y96 are shown with varying amounts of ligand Z253. **B)** The K_D_ titration curves, the color coding for the various concentrations of the ligand can be read from the K_D_ titration curves, which were done with 80 µM KRAS. Due to the weak binding the K_D_ value was not defined.

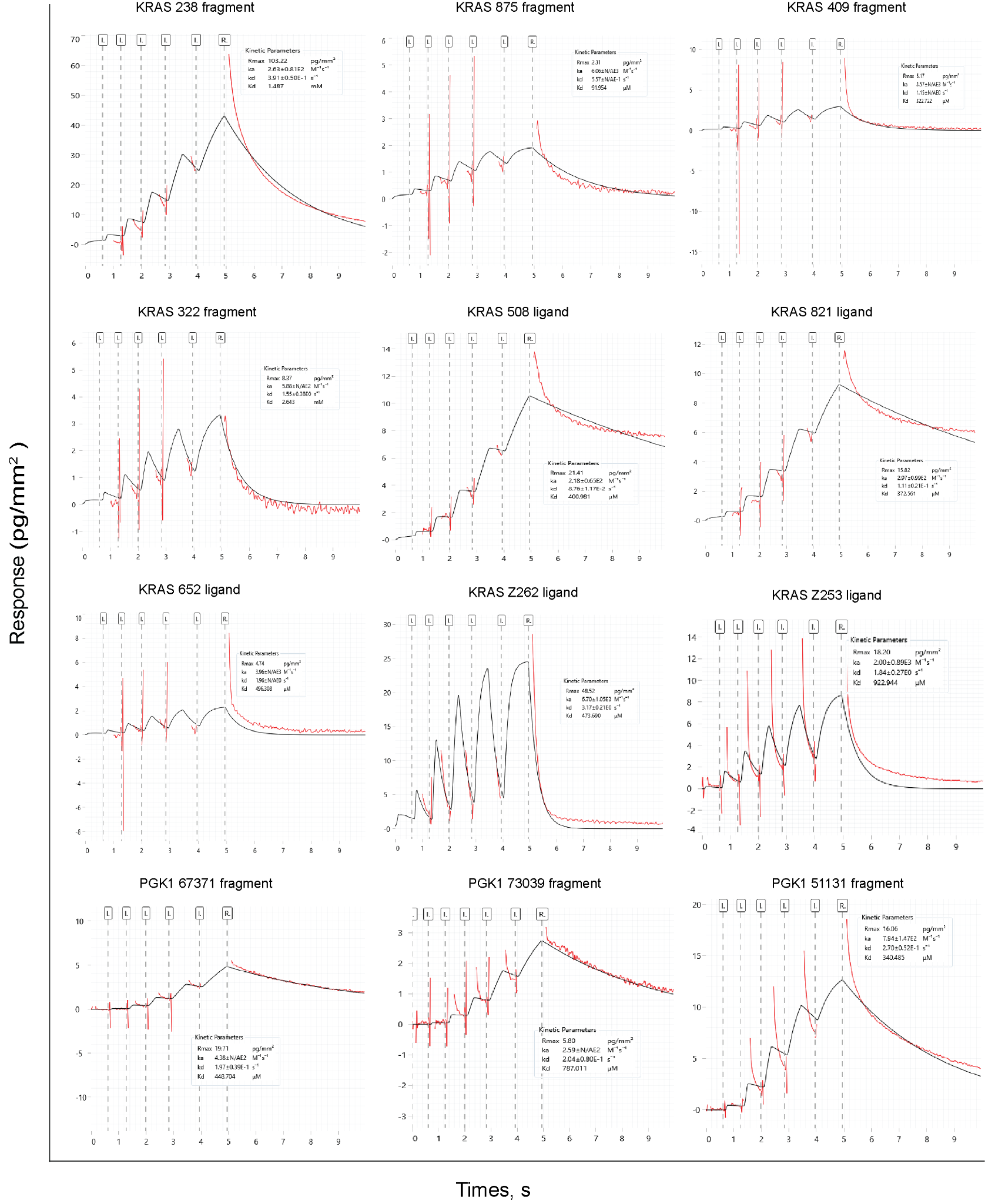

**Figure S14. Representative rapid kinetic sensorgrams from the kinetic screening of library (fragments and ligands) per protein target.** Corresponding values of K_D_ 1 : 1 interaction kinetic model was globally (black traces) fitted to the experimental sensorgrams (red traces) providing the information about the range of affinities.

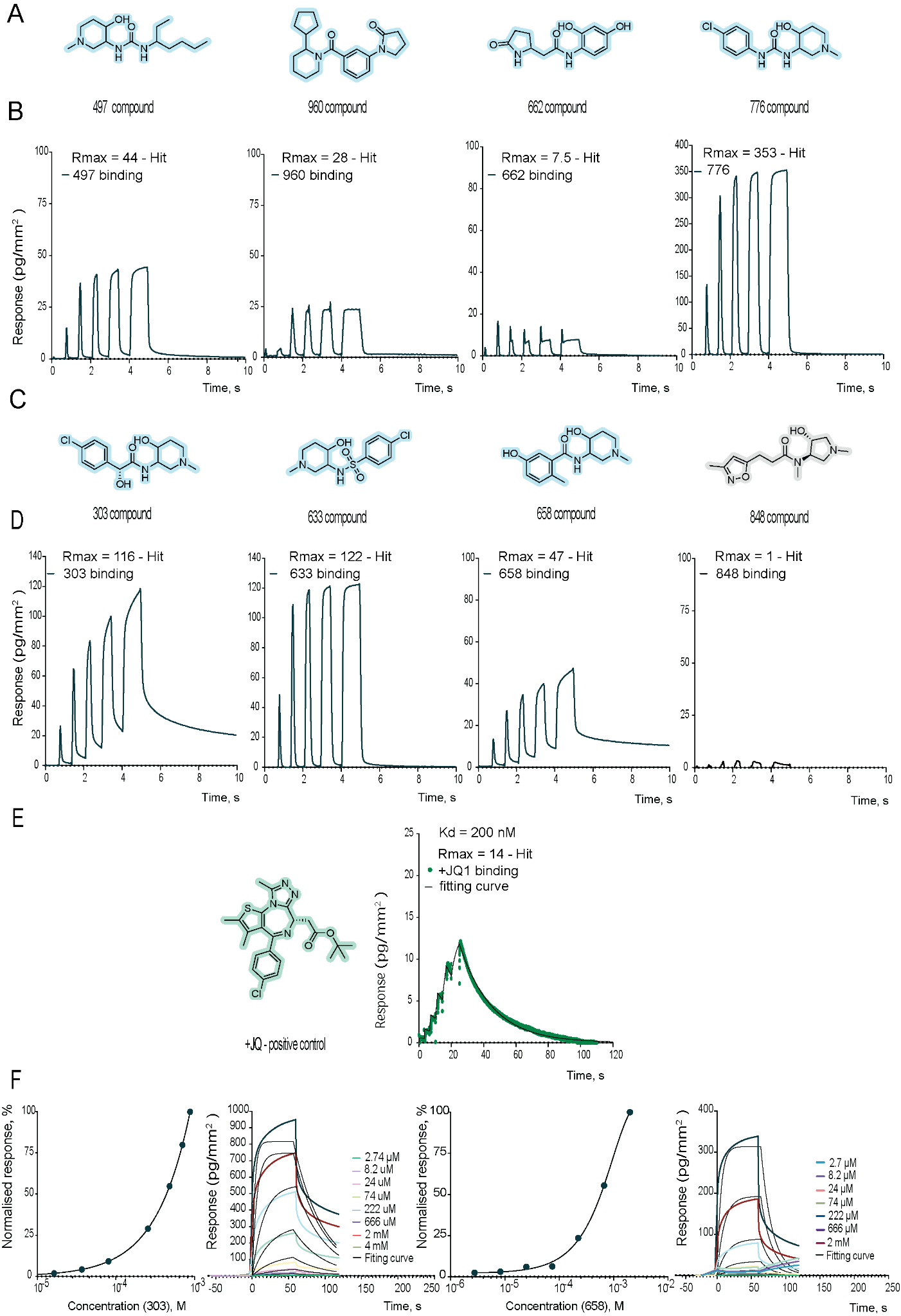

**Figure S15. BrD4 case kinetic data: ligands. A)** Structures of BrD4 ligands (first round of designs). **B)** Examples of binding kinetic sensorgrams (rapid kinetic) for hit ligands (first round of designs). **C)** Structures of BrD4 ligands (second round of designs). **D)** Examples of binding kinetic sensorgrams for hit and non-hit ligands (second round of designs). **E)** Positive control +JQ1 for BrD4: structure and kinetic data, K_D_ = 200 nM. **F)** Multicycle kinetic data and K_D_ determination for BrD4 ligands: 303 and 658 - K_D_ in mM* range (undefined).

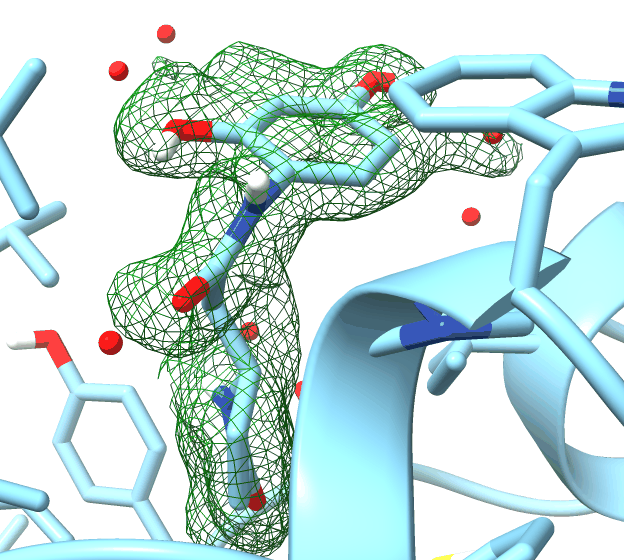

**Figure S16.** Omit map: BrD4 in a complex with 662 compound.

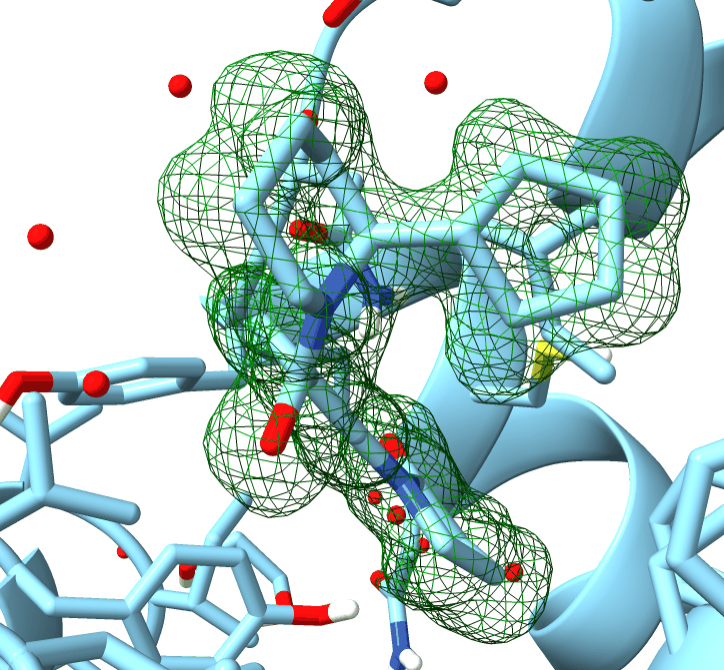

**Figure S17.** Omit map: BrD4 in a complex with 960 compound.

| Target | ID | Smiles | Note | Enamine ID |
| --- | --- | --- | --- | --- |
| KRAS | 770 | CC1=CC(NC(=O)CN2CCC=3NC=4C=CC=CC4C3C2)=NO1 | Derivative of the hit fragment | Z1130197770 |
| KRAS | 652 | O=C(NCCN1C=CC(=O)C=C1)C=2N=CC=C3C=4C=CC=CC4NC23 | Derivative of the hit fragment | Z6265411652 |
| KRAS | 175 | O=C(NCC=1C=CC=CC1)C=2C=C3C(=CN2)NC=4C=CC=CC43 | Derivative of the hit fragment | Z5193516175 |
| KRAS | 400 | O=C(N1CC=2NC(=O)NC(=O)C2C1)C=3N=CC=C4C=5C=CC=CC5NC34 | Derivative of the hit fragment | Z8998113400 |
| KRAS | 530 | CN1CCC=2NC=3C=CC(=CC3C2C1)C(=O)NC4=CNN=C4C(=O)OC(C)(C)C | Derivative of the hit fragment | Z2714949530 |
| KRAS | 935 | CC(=O)NCCOCCN1CCC=2NC=3C=CC(F)=CC3C2C1 | Derivative of the hit fragment | Z2379563935 |
| KRAS | 459 | OC(=O)/C=C/C=1C=C2C(=CN1)NC=3C=CC=CC32 | Derivative of the hit fragment | Z2313083459 |
| KRAS | 362 | CC=1N=CC=C2C=3C=C(O)C=CC3NC12 | Derivative of the hit fragment | Z1216815362 |
| KRAS | Z722 | NS(=O)(=O)C=1C=CC(=CC1)NC(=O)C(=O)NC=2C=CC=3OC(F)(F)OC3C2 | Novel | Z968867222 |
| KRAS | Z957 | NS(=O)(=O)C=1C=CC(=CC1)NC(=O)NCC=2C=CSC2Br | Novel | Z2011030957 |
| KRAS | Z265 | NS(=O)(=O)C=1C=CC=C(C1)CNC(=O)NC=2C=CC=C3OC(F)(F)OC23 | Novel | Z9368007265 |
| KRAS | Z263 | NS(=O)(=O)C=1C=CC=C(C1)NC(=O)NC=2C=CC=3OC(F)(F)OC3C2 | Novel | Z9368007263 |
| KRAS | Z261 | NS(=O)(=O)C=1C=CC=C(C1)CNC=2C=CC=3OC(F)(F)OC3C2 | Novel | Z9368007261 |
| KRAS | Z253 | NS(=O)(=O)C=1C=CC=C(C1)CN2CCC(CC2)S(=O)(=O)C=3C=CC(F)=CC3 | Novel | Z9368007253 |
| KRAS | Z255 | NS(=O)(=O)C=1C=CC(=CC1)NC(=S)NCC=2C=CSC2Br | Novel | Z9368007255 |
| KRAS | Z831 | NS(=O)(=O)C=1C=CC(=CC1)NC(=O)NCC=2C=CC=3OC(F)(F)OC3C2 | Novel | Z2113033831 |
| KRAS | Z124 | NS(=O)(=O)C=1C=CC(=CC1)NNC(=S)NC=2C=CC=3OC(F)(F)OC3C2 | Novel | Z821348724 |
| KRAS | Z262 | NS(=O)(=O)C=1C=CC(=CC1)NCC2=NC=3C=CC(F)=CC3S2 | Novel | Z9368007262 |
| KRAS | 301 | OC=1C=CC=C(F)C1C(=O)NC=2C=CC(OCCN3CCCCC3)=CC2 | Known fragments design setup | Z1724662301 |
| KRAS | 508 | OC=1C=CC=C(F)C1NC(=O)CCOC=2C=CC=CC2Br | Known fragments design setup | Z1576275508 |
| KRAS | 821 | FC=1C=CC=C(OCC(=O)C2=CC=CO2)C1 | Known fragments design setup | Z19755821 |
| KRAS | 291 | OC=1C=CC=C(F)C1C(=O)NCCC=2C=CC(=CC2)N3C=NN=C3 | Known fragments design setup | Z8534392291 |
| KRAS | 16 | FC=1C=CC=C(OCCCC=2C=CC=CC2)C1 | Known fragments design setup | Z54190016 |
| KRAS | 182 | OC=1C=CC=C(F)C1C(=O)NCCOC=2C=CC=CC2Br | Known fragments design setup | Z1724226182 |
| KRAS | 255 | FC=1C=CC=C(OCCNC(=O)C=2C=CC(OCC3CCCCC3)=CC2)C1 | Known fragments design setup | Z8847152255 |
| KRAS | 695 | OC=1C=CC=C(F)C1NC(=O)COC=2C=CC=CC2Br | Known fragments design setup | Z1547524695 |
| KRAS | 238 | CCN1CCC=2NC=3C=CC=CC3C2C1 | Novel fragment | EN300-9295238 |
| KRAS | 875 | O=C1CCC=2C=CC=C3CCN1C32 | Novel fragment | EN300-310875 |
| KRAS | 519 | ClC=1C=CC=2C(=O)NC=NC2C1 | Novel fragment | EN300-23519 |
| KRAS | 718 | FC=1C=CC=CC1Br | Novel fragment | EN300-52718 |
| KRAS | 322 | CC1(C)C(=O)NC=2C=CC=CC21 | Novel fragment | EN300-79322 |
| KRAS | 544 | Oc1cccc(F)c1 | Fragment from a known compound | EN300-19544 |
| KRAS | 409 | C=1C=CC2=C(C1)NC=3C=NC=CC32 | Novel fragment | Z1255372409 |
| BrD4 | 612 | CC(CN1CCCC1=O)NC(=O)C2=CC=NN2C | Known fragments design setup | Z973586612 |
| BrD4 | 280 | O=C(N1CCCC1CC2CCCC2)C=3C=CC(=CC3)N4N=CCC4=O | Known fragments design setup | Z4307830280 |
| BrD4 | 709 | O=C(N1CCCC1CC2CCCC2)C=3C=CC(=CC3)N4CCCC4=O | Known fragments design setup | Z2501079709 |
| BrD4 | 960 | O=C(N1CCCCC1C2CCCC2)C=3C=CC=C(C3)N4CCCC4=O | Known fragments design setup | Z8847151960 |
| BrD4 | 631 | BrC=1C=CC=C(CCNC(=O)N2CCCC2=O)C1 | Known fragments design setup | Z3471560631 |
| BrD4 | 662 | OC=1C=CC(NC(=O)CC2CCC(=O)N2)=C(O)C1 | Known fragments design setup | Z3364074662 |
| BrD4 | 474 | O=C1CCCN1C=2C=CC=C(C2)C=3C=CN=CC3 | Known fragments design setup | Z6739505474 |
| BrD4 | 366 | OC=1C=CC(C(=O)OCC2CCC(=O)N2)=C(O)C1 | Known fragments design setup | Z237684366 |
| BrD4 | 230 | CN1N=CC=C1C2(CC2)NC(=O)N3CCCC3=O | Known fragments design setup | Z4431190230 |
| BrD4 | 100 | OC=1C=CC(NC(=O)C[C@H]2COC(=O)N2)=C(O)C1 | Known fragments design setup | Z8847152100 |
| BrD4 | 008 | ClC=1C=CC(CN(CCN2CCCC2)C(=O)C(=O)C=3C=CC=CC3)=CC1 | Known fragments design setup | Z1941875008 |
| BrD4 | 192 | CN(CC(O)C1CCCC1)C(=O)NCC=2C=CC(=CC2)N3CCCC3=O | Known fragments design setup | Z8847152192 |
| BrD4 | 994 | CN(CCC1CCCC1)C(=O)NCC=2C=CC(=CC2)N3CCCC3=O | Known fragments design setup | Z8847151994 |
| BrD4 | 025 | O=C(NCC=1C=CC=C(C1)N2CCCC2=O)C=3C=CC=CC3C4CCCC4 | Known fragments design setup | Z8847152025 |
| BrD4 | 776 | CN1CCC(O)C(C1)NC(=O)NC=2C=CC(Cl)=CC2 | Known fragments design setup | Z8306448776 |
| BrD4 | 633 | CN1CCC(O)C(C1)NS(=O)(=O)C=2C=CC(Cl)=CC2 | Known fragments design setup | Z8202242633 |
| BrD4 | 492 | CN1CCC(O)C(C1)NC(=O)NCC=2C(C)=NOC2C | Known fragments design setup | Z9101503492 |
| BrD4 | 503 | O[C@@H]1CCCN(CC2CCCCN2C(=O)C=3C=CC=NC3)C1 | Known fragments design setup | Z5602870503 |
| BrD4 | 750 | CCCCC(CC)C(=O)NC1CN(C)CCC1O | Known fragments design setup | Z8424913750 |
| BrD4 | 497 | CCCCC(CC)NC(=O)NC1CN(C)CCC1O | Known fragments design setup | Z9101503497 |
| BrD4 | 956 | CN1CCC(O)C(C1)NC(=O)CC2=CN(C)N=N2 | Known fragments design setup | Z8424873956 |
| BrD4 | 303 | CN1CCC(O)C(C1)NC(=O)[C@H](O)C=2C=CC(Cl)=CC2 | Known fragments design setup | Z8424845303 |
| BrD4 | 496 | CN1CCC(O)C(C1)NCC=2C=CC=C(O)C2Cl | Known fragments design setup | Z9101503496 |
| BrD4 | 886 | CN1CC(CC1=O)C(=O)NC2CN(C)CCC2O | Known fragments design setup | Z8424913886 |
| BrD4 | 649 | CN1CCC(O)C(C1)NC(=O)C=2C=CC=C(O)C2Cl | Known fragments design setup | Z8424848649 |
| BrD4 | 805 | OC1CCCN(CCNC(=O)C=2C=CC=NC2)C1 | Known fragments design setup | Z5533626805 |
| BrD4 | 658 | CN1CCC(O)C(C1)NC(=O)C=2C=C(O)C=CC2C | Known fragments design setup | Z8424887658 |
| BrD4 | 848 | CN([C@@H]1CN(C)C[C@H]1O)C(=O)CCC2=CC(C)=NO2 | Known fragments design setup | Z3587156848 |
| PGK1 | 69861 | CN1CC(=O)NC=2C=CC=CC2C1=O | Novel fragment | EN300-69861 |
| PGK1 | 160442 | CC1SC=2C=CC=CC2NC1=O | Novel fragment | EN300-160442 |
| PGK1 | 16263 | O=C1CSC=2C=CC=CC2N1 | Novel fragment | EN300-16263 |
| PGK1 | 51131 | N#CC=1C=NC=2C=CC=CC2C1 | Novel fragment | EN300-51131 |
| PGK1 | 67371 | OC=1N=CN=CC1Br | Novel fragment | EN300-67371 |
| PGK1 | 229416 | FC(F)(F)C1CCNC2=CC=NN12 | Novel fragment | EN300-229416 |
| PGK1 | 73039 | O=C1CNC(=O)C2CCCN21 | Novel fragment | EN300-73039 |
| PGK1 | 18289 | O=C1NC(=O)C=2N=CNC2N1 | Novel fragment | EN300-18289 |
| PGK1 | 21473 | NC1=NC=2NC=NC2C(=O)N1 | Novel fragment | EN300-21473 |
| PGK1 | 23988 | O=C1NC=NC=2C=CSC12 | Novel fragment | EN300-23988 |
| PGK1 | Z118 | BrC=1C=CC=C(NC(=O)NCC2=CN=C(O2)C=3C=CC=CC3)C1 | Novel | Z9215344118 |
| PGK1 | Z123 | OC(=O)CCN(CC1=CN=C(O1)C=2C=CC=CC2)CC=3C=CC=C(Br)C3 | Novel | Z9215344123 |
| PIN1 | 81 | BrC=1C=CC=2NC=CC2C1 | Novel fragment | EN300-21081 |
| PIN1 | 215001 | NC(=N)C=1C=CC=2C=CC=CC2C1 | Novel fragment | EN300-215001 |
| PIN1 | 250472 | ClC=1C=CC=2CCNC(=O)C2C1 | Novel fragment | EN300-250472 |
| PIN1 | 861 | CN1CC(=O)NC=2C=CC=CC2C1=O | Novel fragment | EN300-69861 |
| PIN1 | 155180 | NC1=NC=2C=CC(Cl)=CC2N1 | Novel fragment | EN300-155180 |
| PIN1 | 18624 | O=C1NS(=O)(=O)C=2C=CC=CC12 | Novel fragment | EN300-18624 |
| PIN1 | 59957 | CN1C(=O)N(C)C=2C=CC=CC12 | Novel fragment | EN300-59957 |
| PIN1 | 497 | NC=1N=C(F)N=C2N=CNC12 | Novel fragment | EN300-697497 |
| PIN1 | 411 | O[C@@H]1[C@H]2CO[C@H](O2)[C@H](O)[C@H]1O | Novel fragment | EN300-7375411 |
| PIN1 | 126031 | OC=1C=CC=C2OCC(=O)NC12 | Novel fragment | EN300-126031 |
| PIN1 | 803 | NS(=O)(=O)C1CCCCC1 | Novel fragment | EN300-59803 |
| PIN1 | 18289 | O=C1NC(=O)C=2N=CNC2N1 | Novel fragment | EN300-18289 |
| PIN1 | 21473 | NC1=NC=2NC=NC2C(=O)N1 | Novel fragment | EN300-21473 |
| PIN1 | 117542 | O=C1CNC(=O)C=2C=CC=CC2N1 | Novel fragment | EN300-117542 |
| PIN1 | 73335 | [O-][N+](=O)C1=CNC(=O)NC1=O | Novel fragment | EN300-73335 |
| PIN1 | 46887 | CC=1C=CC=C2C(=O)NC=NC12 | Novel fragment | EN300-76887 |
| PIN1 | 21022 | CN1C(=O)NC(=O)C=2NC=NC12 | Novel fragment | EN300-212022 |
| PIN1 | Z126 | O=C1COC=2C=CC=C(OCC=3C=CC=C(Br)C3)C2N1 | Novel | Z1587591126 |
| PIN1 | Z400 | CN1C(=O)N(C)C=2C=C(C=CC21)NS(=O)(=O)C=3C=CC(=CC3)S(N)(=O)=O | Novel | Z955490400 |
| PIN1 | Z819 | O=C(CNS(=O)(=O)C=1C=CC=CC1)NC=2C=C(Cl)SC2Cl | Novel | Z9426861819 |
| PIN1 | Z821 | NS(=O)(=O)C1=NN=C(NC(=O)NCC=2C=CC=CC2C=3C=CC=CC3)S1 | Novel | Z9426861821 |
| PIN1 | Z820 | O=C(CN1C(=O)CSC=2C=CC=CC21)NC=3C=CC=C(C3)C=4C=CC=CC4 | Novel | Z955490400 |
| PIN1 | Z284 | CC(C)(C(=O)NC1=NC2=CC=C(Cl)C=C2N1)C=3C=CC(=CC3)C=4C=CC=CC4 | Novel | Z9588666284 |
| PIN1 | Z285 | O=C(CC=1C=CC(=CC1)C=2C=CC=CC2)NC3=NC=4C=CC(Cl)=CC4N3 | Novel | Z9588666285 |
| SARS | 21473 | NC1=NC=2NC=NC2C(=O)N1 | Novel fragment | EN300-21473 |
| SARS | 26993 | NS(=O)(=O)C=1C=CC=CC1F | Novel fragment | EN300-26993 |
| SARS | 18289 | O=C1NC(=O)C=2N=CNC2N1 | Novel fragment | EN300-18289 |
| SARS | 5190 | CC1=NC=2C=CC=CC2C(=O)N1 | Novel fragment | EN300-05190 |
| SARS | 21048 | OC=1C(Cl)=CC=CC1Cl | Novel fragment | EN300-21048 |
| SARS | 97050 | ClC=1C=C(Cl)C=C(C#N)C1 | Novel fragment | EN300-97050 |
| SARS | 111949 | OC(=O)C=1N=CC=C2C=CC=CC12 | Novel fragment | EN300-55949 |
| SARS | Z104 | OC=1C(Cl)=CC=C(NC(=O)CCNC(=O)NC=2C=CC(Br)=CC2)C1Cl | Novel | Z9215344104 |
| SARS | Z106 | NC=1C=CC(Br)=NC1C(=O)NCC(=O)NC=2C=CC(Br)=CC2 | Novel | Z9215344106 |
| SARS | Z559 | NC=1C=CC(Br)=CC1C(=O)NC=2C=C(O)C=CC2Br | Novel | Z4968271559 |

**Table 1. The list of compounds.**

| **Compound id** | **Concentration tested** | **Binder** |
| --- | --- | --- |
| 160442 | 0.5 mM | No |
| 21473 | 0.5 mM | No |
| 23988 | 0.5 mM | Yes |

**Table 2. NMR tested PGK1 novel fragments.**

| **Compound id** | **Concentration tested** | **Binder** |
| --- | --- | --- |
| 26993 | 0.5 mM | Yes |
| 21048 | 0.5 mM | Yes |
| Z104 | 0.5 mM | Yes |

**Table 3. NMR tested SARS Mpro novel fragments and ligands.**

| **Compound id** | **Concentration tested** | **Binder** |
| --- | --- | --- |
| 238 | 2 mM | Yes, large shifts at HSQC. |
| 875 | 2 mM | Yes, weak |
| 519 | 2 mM | No discernable shifts |
| 718 | 2 mM | No discernable shifts |
| 322 | 2 mM | Yes, weak |

**Table 4. NMR tested KRAS novel fragments.**

| **Compound id** | **Concentration tested** | **Binder** |
| --- | --- | --- |
| 861 | 0.5 mM | Yes |
| 126031 | 0.5 mM | Yes |

**Table 5. NMR tested PIN1 novel fragments.**

| **Compound id** | **Concentration tested** | **Binder** |
| --- | --- | --- |
| 695 | 0.5 mM | Small shift |
| 182 | 0.5 mM | No |
| 255 | 2 mM | No |
| 629 | 2 mM | No |
| 301 | 1 mM | Yes |
| 508 | 1 mM | Yes, Kd> 2.5 mM |
| 821 | 1 mM | Yes, very weak |
| 291 | 2 mM | No |

**Table 6. NMR tested KRAS compounds (known fragment, novel linker).**

| **Compound id** | **Concentration tested** | **Binder** |
| --- | --- | --- |
| Z253 | 2 mM | Yes |
| 362 | 2 mM | Yes |
| Z261 | 2 mM | No |
| Z957 | 2 mM | No |
| 770 | 2 mM | Yes |
| Z265 | 2 mM | No |
| Z262 | 2 mM | No |
| 695 | 2 mM | No |
| Z722 | 2 mM | No |
| 175 | 2 mM | No |

**Table 7. NMR tested KRAS derivatives of novel fragments, de novo molecules.**

| Compound | RMSD Top1 | RMSD Best Top5 |
| --- | --- | --- |
| z559 | 3.54 | 1.926 |
| z104 | 6.76 | 6.172 |
| z106 | 4.574 | 4.24 |

**Table 8. RMSD values for docking of compounds.**

| **Target** | **Compound** | **Rmax** | **Ass. Error** | **Diss. Error** | **HIT?** |
| --- | --- | --- | --- | --- | --- |
| KRAS | Z957 | 152 | 27 | 7 | YES |
| KRAS | 770 | 129 | 21 | 6 | YES |
| KRAS | Z255 | 75 | 14 | 5 | YES |
| KRAS | 175 | 0 | 480 | 2.2×10^8^ | NO |
| KRAS | BI-2865 | 23 | 16 | 20 | YES |
| KRAS | 238 | 212 | 34 | 22 | YES |
| KRAS | 409 | 180 | 21 | 6 | YES |
| KRAS | 322 | 34 | 25 | 9.8 | YES |
| KRAS | 718 | 6 | 63 | 5.2×10^7^ | NO |
| BrD4 | 497 | 44 | 17 | 93 | YES |
| BrD4 | 960 | 28 | 4.6 | 11 | YES |
| BrD4 | 662 | 7.6 | 0 | 0 | YES |
| BrD4 | 956 | 0 | 6.2×10^3^ | 4×10^8^ | NO |
| BrD4 | +JQ1 | 14 | 0 | 0 | YES |
| BrD4 | 303 | 116 | 16 | 10 | YES |
| BrD4 | 633 | 122 | 48 | 17 | YES |
| BrD4 | 658 | 47 | 17 | 16 | YES |
| BrD4 | 776 | 353 | 29 | 6.7 | YES |
| BrD4 | 848 | 1 | 1.1×10^3^ | 8×10^7^ | NO |
| PGK1 | Z123 | 403 | 43 | 14 | YES |
| PGK1 | 442 | 34 | 36 | 9.8 | YES |
| PGK1 | 51131 | 22 | 74 | 24 | YES |
| PGK1 | CHR-6494 | 12 | 16 | 5.1 | YES |
| PGK1 | 69861 | 87 | 75 | 11 | YES |
| PGK1 | Z118 | 0 | 1.1×10^3^ | 52 | NO |
| SARS | 26993 | 23 | 25 | 11 | YES |
| SARS | 5190 | 32 | 25 | 34 | YES |
| SARS | 21048 | 102 | 47 | 12 | YES |
| SARS | 97050 | 22 | 27 | 7.1 | YES |
| SARS | 18289 | 0 | 5.9×10^3^ | 9.9×10^2^ | NO |
| SARS | Nirmatrelvir | 79 | 42 | 42 | YES |
| SARS | Z104 | 20 | 68 | 32 | YES |
| PIN1 | 861 | 48 | 23 | 29 | YES |
| PIN1 | 73335 | 20 | 78 | 26 | YES |
| PIN1 | 803 | 4 | 1.3×10^3^ | 4.5×10^2^ | NO |
| PIN1 | 18624 | 22 | 55 | 18 | YES |
| PIN1 | DTM | 3.2 | 39 | 52 | YES |
| PIN1 | 126031 | 9 | 76 | 89 | YES |
| PIN1 | Z820 | 0 | 1.1×10^3^ | 52 | NO |

**Table 9. Hit criteria table for all targets: KRAS, BrD4, PGK1, SARS Mpro, PIN1.**

The experimental hits are defined on two criteria: 1) Observed R_max_ > 50% of R_max_ (positive control), 2) dissociation and association errors are ≤ 100%. The relative diss. and ass. errors are calculated as the 95% confidence intervals of the dissociation rate (K_D_) and association rate (K_A_) estimates divided by their values.

| Target protein | Sequence |
| --- | --- |
| KRAS | MHHHHHHGSWGSGMTEYKLVVVGAGGVGKSALTIQLIQNHFVDEYDPTIEDSYRKQVVIDGETCLLDILDTAGQEEYSAMRDQYMRTGEGFLCVFAINNTKSFEDIHHYREQIKRVKDSEDVPMVLVGNKSDLPSRTVDTKQAQDLARSYGIPFIETSAKTRQGVDDAFYTLVREIRKHKEK |
| SARS-CoV-2 Main Protease | MSGSGFRKMAFPSGKVEGCMVQVTCGTTTLNGLWLDDVVYCPRHVICTSEDMLNPNYEDLLIRKSNHNFLVQAGNVQLRVIGHSMQNCVLKLKVDTANPKTPKYKFVRIQPGQTFSVLACYNGSPSGVYQCAMRPNFTIKGSFLNGSCGSVGFNIDYDCVSFCYMHHMELPTGVHAGTDLEGNFYGPFVDRQTAQAAGTDTTITVNVLAWLYAAVINGDRWFLNRFTTTLNDFNLVAMKYNYEPLTQDHVDILGPLSAQTGIAVLDMCASLKELLQNGMNGRTILGSALLEDEFTPFDVVRQCSGVTGSWGSHHHHHH |
| BrD4 | MHHHHHHSSGVDLGTENLYFQSMNPPPPETSNPNKPKRQTNQLQYLLRVVLKTLWKHQFAWPFQQPVDAVKLNLPDYYKIIKTPMDMGTIKKRLENNYYWNAQECIQDFNTMFTNCYIYNKPGDDIVLMAEALEKLFLQKINELPTEE |
| PIN1 | MEPARVRCSHLLVKHSQSRRPSSWRQEQITRTQEEALELINGYIQKIKSGEEDFESLASQFSDCSSAKARGDLGAFSRGQMQKPFEDASFALRTGEMSGPVFTDSGIHIILRTEGSWGSGHHHHHH |
| PGK1 | MGSHMSLSNKLTLDKLDVKGKRVVMRVDFNVPMKNNQITNNQRIKAAVPSIKFCLDNGAKSVVLMSHLGRPDGVPMPDKYSLEPVAVELKSLLGKDVLFLKDCVGPEVEKACANPAAGSVILLENLRFHVEEEGKGKDASGNKVKAEPAKIEAFRASLSKLGDVYVNDAFGTAHRAHSSMVGVNLPQKAGGFLMKKELNYFAKALESPERPFLAILGGAKVADKIQLINNMLDKVNEMIIGGGMAFTFLKVLNNMEIGTSLFDEEGAKIVKDLMSKAEKNGVKITLPVDFVTADKFDENAKTGQATVASGIPAGWMGLDCGPESSKKYAEAVTRAKQIVWNGPVGVFEWEAFARGTKALMDEVVKATSRGCITIIGGGDTATCCAKWNTEDKVSHVSTGGGASLELLEGKVLPGVDALSNIGSWGSGHHHHHH |

**Table 10. Target protein and sequence.**

|  | **BRD4-662** | **BRD4-960** | **PIN1-155180** |
| --- | --- | --- | --- |
|  | *native* | *native* | *native* |
| **Data collection** |  |  |  |
| Wavelength (Å) | 0.96546 | 0.87313 | 0.87 |
| Space group | P 21 21 21 | P 21 21 21 | C 1 2 1 |
| *a, b, c* (Å) | 31.96, 47.27, 79.04 | 32.37, 47.38, 79.03 | 118.1, 36.3, 52.0 |
| α, β, γ (°) | 90, 90, 90 | 90, 90, 90 | 90 ,100.4, 90 |
| Resolution (Å) * | 19.00 - 1.21 (1.23 – 1.21) | 40.64 – 1.31 (1.33 – 1.31) | 51.1 - 1.6 (1.7 - 1.6) |
| R_meas_* | 0.076 (1.021) | 0.127 (1.529) | 0.11 (0.75) |
| R_merge_* | 0.064 (0.772) | 0.109 (1.310) | 0.093 (0.67) |
| Mean I/σI* | 7.7 (1.0) | 5.7 (1.0) | 13.4 (2.4) |
| Completeness (%)* | 96.2 (76.8) | 99.2 (99.2) | 99.9 (100.00) |
| Multiplicity* | 3.3 (1.7) | 3.9 (3.7) | 4.6 (4.8) |
| CC1/2* | 0.995 (0.581) | 0.996 (0.444) | 0.98 (0.74) |
| **Refinement** |  |  |  |
| Resolution (Å) | 19.00 - 1.21 | 40.64 – 1.31 | 51.1 - 1.6 |
| Total reflections* | 116689 (2317) | 115836 (5398) | 134142 (13703) |
| Total unique* | 35529 (1390) | 29805 (1470) | 28927 (2874) |
| *R_work_^#^* | 0.1440 (0.3681) | 0.1455 (0.2708) | 0.19 (0.27) |
| *R_free_^#^* | 0.1669 (0.3829) | 0.1870 (0.3027) | 0.21 (0.29) |
| Number of non-hydrogen atoms | 1274 | 1302 | 2125 |
| macromolecules | 1114 | 1098 | 1850 |
| ligands | 34 | 50 | 67 |
| solvent | 126 | 154 | 208 |
| Protein residues | 126 | 126 | 227 |
| RMS deviations (bonds)^#^ | 0.007 | 0.006 | 0.024 |
| RMS deviations (angles)^#^ | 0.89 | 0.90 | 3.1 |
| Ramachandran favored (%)^#^ | 98.39 | 98.39 | 98.2 |
| Ramachandran allowed (%)^#^ | 1.61 | 1.61 | 1.8 |
| Ramachandran outliers (%)^#^ | 0.00 | 0.00 | 0.00 |
| Rotatmer outliers (%)^#^ | 0.00 | 0.00 | 0.00 |
| Clashscore^#^ | 2.19 | 1.74 | 8.3 |
| Average B-factor^#^ | 17.75 | 15.91 | 22.5 |
| macromolecules | 16.07 | 14.30 | 21.2 |
| ligands | 26.69 | 21.10 | 31.1 |
| solvent | 30.16 | 25.70 | 31.1 |
| Molprobity Score | **0.99** | **0.93** | **1.03** |
| PDB | **9T15** | **9T16** | **9T04** |

Statistics for the highest-resolution shell are shown in parentheses.

**Table 11. Data collection and refinement statistics.**
